# Modeling causal signal propagation in multi-omic factor space with COSMOS

**DOI:** 10.1101/2024.07.15.603538

**Authors:** Aurelien Dugourd, Pascal Lafrenz, Diego Mañanes, Victor Paton, Robin Fallegger, Anne-Claire Kroger, Denes Turei, Yunfan Bai, Yuxin Li, Michael Trogdon, Drew Nager, Shibing Deng, Chen Shen, John D. Lapek, Blerta Shtylla, Julio Saez-Rodriguez

## Abstract

Understanding complex diseases requires approaches that jointly analyze omics data across multiple biological layers, including signaling, gene regulation, and metabolism. Existing data-driven multi-omics analysis methods, such as multi-omics factor analysis (MOFA), can identify associations between molecular features and phenotypes, but they are not designed to integrate existing mechanistic molecular knowledge, which can provide further actionable insights. We introduce an approach that connects data-driven analysis of multi-omics data with systematic integration of mechanistic prior knowledge using COSMOS+ (Causal Oriented Search of Multi-Omics Space). We show how factor analysis output can be used to estimate activities of transcription factors and kinases as well as ligand-receptor interactions, which in turn are integrated with network-level prior-knowledge to generate mechanistic hypotheses about paths connecting deregulated molecular features. We apply this approach on a novel multi-omics dataset of cell line models of breast cancer resistance to evaluate the ability of such mechanistic hypotheses to identify resistance drivers, as well as a breast cancer patient cohort. Our approach offers an interpretable framework to generate actionable insights from multi-omic data particularly suited for high dimensional datasets.

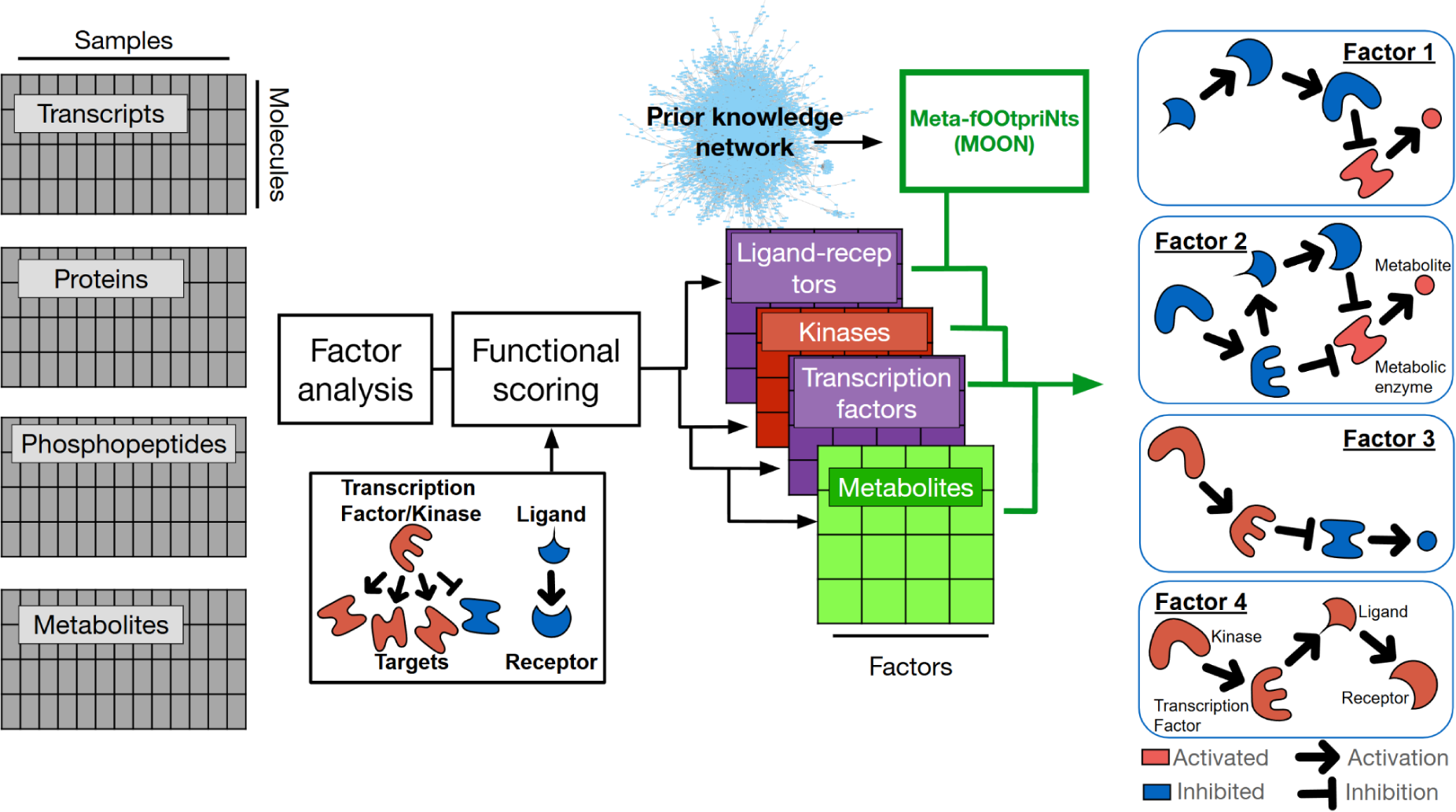

## 1. Introduction

Many diseases are the result of complex deregulations of several interconnected biological mechanisms that span over cell-cell communication mediated by ligands and receptors, intracellular signaling, gene regulation, and metabolism. Consequently, measuring multiple types of omics data, such as transcriptomics, proteomics, and metabolomics, is becoming increasingly popular (Chen *et al*, 2023; Hasin *et al*, 2017; Subramanian *et al*, 2020; Conesa & Beck, 2019) to understand such diseases (Yugi *et al*, 2016; Terakawa *et al*, 2022).

There are many analysis methods that can help researchers extract relevant insights from such complex datasets. Among them, factor analysis is a popular type of dimensionality reduction method for multi-omics data analysis. It provides a data driven estimation of the different axis of variability across samples and modalities (Argelaguet *et al*, 2020; Thurstone, 1931; Velten & Stegle, 2023; Brown *et al*, 2023; Velten *et al*, 2022; Frankhouser *et al*, 2024). Each omic feature is associated with weights related to latent factors, effectively untangling the sources of variability of the dataset. Such an unsupervised approach is particularly powerful when multi-omic datasets are generated from a large number of samples, such as cell line collections and patient cohorts (Quintero *et al*, 2020; Barretina *et al*, 2012; Yang & Michailidis, 2016; Rau *et al*, 2022; Freeman-Cook *et al*, 2021a; Argelaguet *et al*, 2018; Su *et al*, 2011; Wang *et al*, 2021; Garnett *et al*, 2012; Knowles *et al*, 2019), where group comparison is not trivial as there can be many different groups in a cohort, and the groups themselves are rarely homogeneous.

While very useful, the resulting statistical association in multi-omics factors and their associated weights are purely data-driven. They can be further analyzed to provide molecular mechanistic context, and this is typically done by pathway enrichment analysis and/or association with clinical features (Argelaguet *et al*, 2019; Consiglio *et al*, 2020; Gonçalves *et al*, 2022; Schlechte *et al*, 2023; Li *et al*, 2022; Monaco *et al*, 2022; Hamsanathan *et al*, 2022; Park *et al*, 2022; Kwok *et al*, 2023; Gambacorta *et al*, 2022; Mangiante *et al*, 2023). Besides pathways, Ligand-Receptor (LR) mediated cell-cell communication events can also be explored in the context of such factor weights (Armingol *et al*, 2021), by scoring the co-association of ligands and their receptors within factors, using a similar statistical framework as pathway analysis (Dimitrov *et al*, 2024; Efremova *et al*, 2020). However, pathway enrichment and ligand receptor analysis methods are limited in their ability to provide functional and mechanistic insights - they score by analyzing the expression level (or in this case, factor weights) of their genes, which does not always reflect activity (that is, their actual influence on the biological processes they are involved in) (Szalai & Saez-Rodriguez, 2020). Like the enrichment of expression of pathway components, LR co-expression scoring is only an approximation of cell-cell communication since LR co-expression is not equivalent to an LR biochemical activation; it rather allows us to prioritize which of them is more likely to induce intracellular signaling changes in a given condition (Armingol *et al*, 2021; Schäfer *et al*, 2024).

Intracellular processes can be estimated from the affected molecules e.g. the regulation of abundance of target transcripts by a transcription factor. These so-called footprint methods (Dugourd & Saez-Rodriguez, 2019) share the statistical framework of pathway enrichment analysis (Badia-I-Mompel *et al*, 2022) and their output is typically more biochemically accurate than classic pathway scoring (Szalai & Saez-Rodriguez, 2020). Therefore, footprint methods can potentially help us characterize factor analysis results at a biological level as it brings complementary biological insights to pathways and LR analyses.

Nonetheless, such analyses only output sets of disconnected feature scores (e.g. individual TFs or LR pair scores). TFs and LRs can be further connected together using networks derived from existing knowledge about interactions among molecules (prior knowledge networks) (Liu *et al*, 2019; Dugourd *et al*, 2021; Cancer Genome Atlas Research Network, 2013; Bradley & Barrett, 2017; Chowdhury *et al*, 2022; Massacci *et al*, 2023; Rosenberger *et al*, 2024; Garrido-Rodriguez *et al*, 2022) using methods that rely on various formalisms (such as Integer Linear Programming or network diffusion) to find paths in the network that can explain the orchestrated changes in individual processes (Gjerga *et al*, 2021; Venafra *et al*, 2024). They are particularly useful to propose mechanistic and testable hypotheses that can link alterations such as stimulation via a ligand or perturbation by a drug with downstream intracellular signaling, gene regulation, and metabolism. These network approaches currently suffer from several limitations. First, their computational complexity can limit their use in contexts where potentially many single samples, contrasts or factors need to be analyzed. Second, they often provide many mechanistic hypotheses, making it difficult for researchers to prioritize them for validation and organize them into a coherent narrative. Third, these methods inherently rely on the quality of the prior knowledge networks that are used. Errors in the prior knowledge can lead to false molecular interactions being highlighted as relevant to explain a given experimental result. Another limit of such prior knowledge based mechanistic hypothesis generation methods is that it remains unclear how useful those hypotheses are to answer specific biological questions such as the connection between drug treatment mechanistic response and resistance drivers.

In this work, we provide an integrated solution to these challenges by developing an approach to bridge multi-omic data-driven analysis with methods that can put them into the context of biological mechanisms beyond classic pathway enrichment analysis, including footprint analysis and mechanistic network integration. We first demonstrate how multi-omic factor weights can be used to characterize functional features associated with factors, such as Transcription Factors (TF) activity (Müller-Dott *et al*, 2023) and LR mediated cell-cell communication events (Dimitrov *et al*, 2024). We also developed a lightweight alternative network scoring procedure for COSMOS called Meta-fOOtprint aNalysis (MOON) that can efficiently contextualize prior knowledge networks into mechanistic hypotheses, therefore allowing the use of footprints and network methods with large datasets without necessitating explicit experimental structures. MOON can generate mechanistic hypothesis spanning over signaling and metabolism, effectively connecting perturbations observed at the level of cells kinase receptors with downstream transcription factor and metabolic de-regulations, as well as interactions between metabolic ligands and chemicals with downstream receptor and signaling cascades. We also assessed the ability of such network scoring procedures to capture relevant regulators of gene expression by applying it to a dataset of transcriptomic changes upon cytokine stimulation ( Jiang *et al*, 2021), and provide functionalities to facilitate the interpretation of such networks by end users, as well as making it easier to pinpoint errors in prior knowledge. We refer to the new version of COSMOS that incorporates the MOON function and the interface to Factor Analysis as COSMOS+. We used this updated COSMOS+ method to extract coherent mechanistic insight such as a crosstalk between the JAK-STAT pathway, Citrate metabolism and MYC inhibition that was specifically associated with leukemic cell lines in the NCI60 dataset (Su *et al*, 2011).

We then applied the COSMOS+ framework to a novel multi-omics/multi-timepoint dataset comprising transcriptomic and phosphoproteomic measurements of three breast cancer cell lines with different resistance profiles treated with various regimens of Cyclin-dependent kinase (CDK) inhibitors. We show that mechanistic hypotheses of signaling deregulation in resistant and sensitive cell lines treated with CDK inhibitors are correlated with resistance markers identified through knock-out screening. Finally, we apply the COSMOS+ framework to a breast cancer patient cohort, and we demonstrate the complementarity of the resulting features with omic data to predict patient outcomes.

To facilitate the use of the open-source COSMOS+ methods to any multi-omics dataset, we provided the software both in R and python (as part of the networkcommons (Paton *et al*, 2024) framework) along with its code and tutorials (R: https://github.com/saezlab/cosmosR, python: https://github.com/saezlab/networkcommons).

## 2. Results

### 2.1 Presentation of COSMOS+

COSMOS+ contextualizes generic resources of prior knowledge spanning protein/protein and protein/metabolite mechanistic interactions. It uses omic features resulting from different types of statistical analysis, such as factor weights resulting from the variance decomposition of multi-omic datasets. COSMOS+ is tailored to work with transcriptomic, phospho-proteomic or multi-omics data sets that contain at least two of the three following types of data: transcriptomics, phospho-proteomics and metabolomics. It can also be adapted to additional data types, as long as they can be mapped to functional alterations of proteins, e.g. protein conformation changes (Burtscher *et al*, 2024).

First, once a multi-omics dataset has been decomposed into a latent factor by a given factor analysis method, such as MOFA (Argelaguet *et al*, 2020), it can be further processed through the COSMOS+ method to generate factor-specific mechanistic hypotheses as follows. For each factor, transcription factor activities and ligand-receptor scores are estimated from transcriptomic data, and kinase activities are estimated from phospho-proteomic data. The activities are estimated by modeling each TF and kinase as a linear regression of their target measurements as function of their mode of regulations (see methods). Potential cell-cell interaction events are scored as a function of the co-regulation of pairs of ligands and receptors, normalized by the background weights of all other genes of the factor considered. To do so, the scores are estimated by modeling each pair of potential ligand and receptor (according to prior knowledge resource of ligand-receptor pairs) as a linear regression of the transcriptomic and/or proteomic measurements as function of their belonging to a given ligand-receptor pair (Figure 1A).

**Figure 1:**
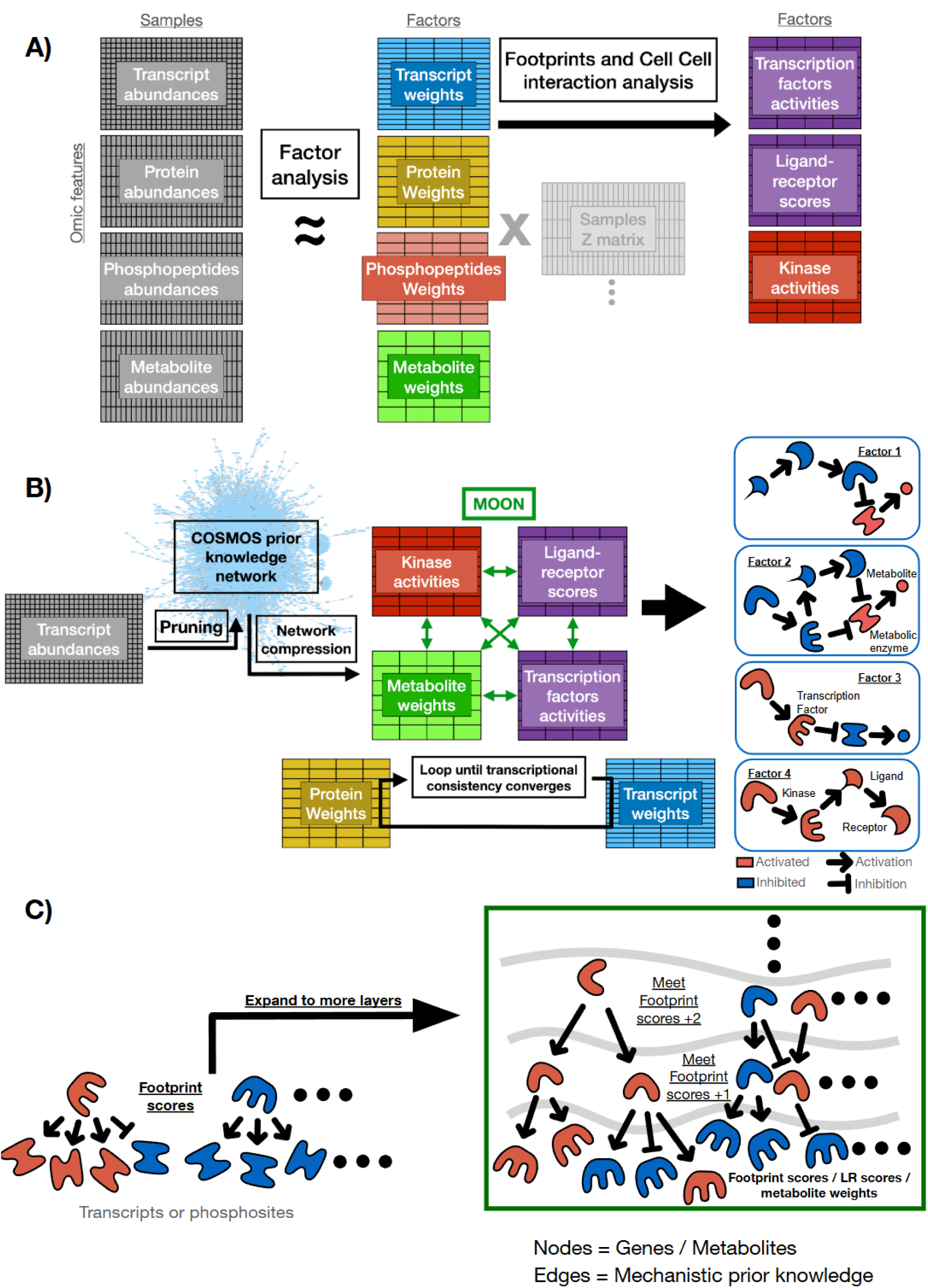
Schematic representation of the COSMOS+ method. A) footprint and CCC interactions in factor space. Representation of the types of data that are currently primarily handled by COSMOS+, and how they can be processed through factor analysis to generate corresponding footprint (TFs and kinases) and ligand-receptor scores. B) Mechanistic network hypothesis generation. Schematic representation of the integration of the feature weights, TF, kinase and LR score with a prior knowledge network. C) Meta-footprint. Schematic representation of the concept of the Meta-fOOtprint aNalysis (MOON), which expands the concept of footprint to upstream layers. Scores are usually estimated with decoupleR (Badia-I-Mompel et al, 2022).

The next steps connect the top contributing ligands, receptors, kinases, transcription factors and metabolites of a given factor in a consistent network of mechanistic hypotheses. It does so using the COSMOS prior knowledge spanning over ligand-receptor interactions, intracellular signaling, transcriptional regulation and metabolic reactions. The prior knowledge network consists of curated molecular interactions coming from the Omnipath database, STITCH database and recon3D human reaction network (Szklarczyk *et al*, 2016; Türei *et al*, 2021; Brunk *et al*, 2018), and its building process is now automated and integrated within Omnipath. If transcriptomic data is available, it can be used to perform an initial pruning. This pruning aims at removing mechanisms (e.g. post-translational modification or metabolic reactions) that are mediated by genes that are not expressed in any of the analyzed samples. Then, the network is pruned to keep only mechanisms that are in-between the top (based on user defined threshold, see Methods) contributing ligands, receptors, kinases, transcription factors and metabolites of a given factor within a set number of steps. This ensures the removal of elements that have no connecting path to perturbed (approximate non-controllable) or measured nodes (approximate non-observable) (Saez-Rodriguez *et al*, 2009). Then, redundant paths in the network (identified from parent nodes that share the exact same set of children nodes) are compressed (Supplementary Figure S1A/B).

Next, the new network meta-footprint method (MOON) is used to score the most consistent mechanisms of the prior knowledge network (PKN) connecting the ligands, receptors, kinases, transcription factors and metabolites (Figure 1B). Briefly, MOON builds upon footprint activity scoring and network diffusion (Wu *et al*, 2008; Leiserson *et al*, 2015) to perform an iterative footprint scoring (hence called ‘meta-footprint’) over a given prior knowledge network, starting from a downstream layer composed of the input nodes (Figure 1C). For example, if we consider a set of scored TFs mapped on a prior knowledge network, we can score nodes directly upstream of those TFs in the network, based on the consistency of the sign of TF activity score they directly regulate. This process can be repeated iteratively over all the nodes of the PKN (provided a node has a path downstream that can reach the input nodes). Thus, to run MOON, the user specifies which set of inputs will be considered as downstream layer and, optionally, upstream layer. This decision depends on the biological question that is considered. For example, in the case of studying the connections between receptor, kinases and TFs, the downstream layer can be set as the TFs and the upstream layer as receptors (from ligand-receptor pairs) and kinases (estimated from phospho-proteomic data). MOON will iteratively score nodes upstream of the TFs until it reaches the upstream receptor/kinase layer (if the later is provided, else until it reaches the maximum number of steps allowed by the user), and then compare the MOON score of those receptors/kinases with the upstream input scores (receptors scores and kinase activities). Any receptor/kinase that shows a sign incoherence between its MOON score and the input score/measurement is pruned out along with all incoming and outcoming edges. A transcriptional consistency check can be performed that removes any interaction between TF and inconsistently regulated direct downstream targets (incoherence between sign of the TF activity score and the sign of the downstream measurement/factor weight input). After the interactions are removed, the MOON scoring procedure can be repeated with the new pruned network, until no more interactions are removed. Finally, after scoring the nodes of the network, MOON also specifically prunes out any interaction from the prior knowledge network that would be inconsistent with the MOON node scores (e.g. two nodes with opposite scores connected by a positive interaction) (see methods). The output of MOON is a multi-omics network representing a set of potential mechanistic interactions consistent with the input data.

### 2.2 MOON cytokine prediction with the Cytosig dataset

We used MOON to score the nodes of the COSMOS network for each of the 1,359 Cytosig ( Jiang *et al*, 2021) ligand expression signatures. Each of the Cytosig ligand expression signatures corresponds to the transcriptional change observed after the application of a given ligand on a cell population. These signatures have been obtained from systematic query and automated processing of several large RNA data repositories. For each of Cytosig’s ligand expression signatures, the MOON scores of the corresponding ligand could be estimated in 549 out of 1,359 expression signatures across 63 unique ligands (31 ligands were not reachable upstream of transcription factors in the COSMOS prior knowledge within the specified number of steps, therefore their score couldn’t be estimated). Out of 63 unique ligands, 47 (75%) of them had a positive MOON score on average across their corresponding signatures (16 of the unique ligands significantly different from 0, t.test p-value <= 0.05) and the rest a negative score on average (4 of the unique ligands significantly different from 0, t.test p-value <= 0.05) (Figure 2A). To assess the impact of the time after perturbations as a potential confounding factor, we calculated the correlation between the MOON score of Cytosig ligands and the time delay of RNAseq measurement after perturbation for each signature, as well as between the MOON score and the shortest distance between estimated TF scores and upstream nodes of the network. The correlation between MOON score and time delay was -0.12 (Kendall’ Tau coefficient, p-value = 0.002). There was no significant correlation (Kendall’s tau = 0.03, p-value = 0.33) between the score of the ligand and the number of steps that separated them from their informative TFs (referred to as “level”). Thus, the time of measurement, but not the steps of the network, seem to influence the MOON score of the Cytosig ligands.

**Figure 2:**
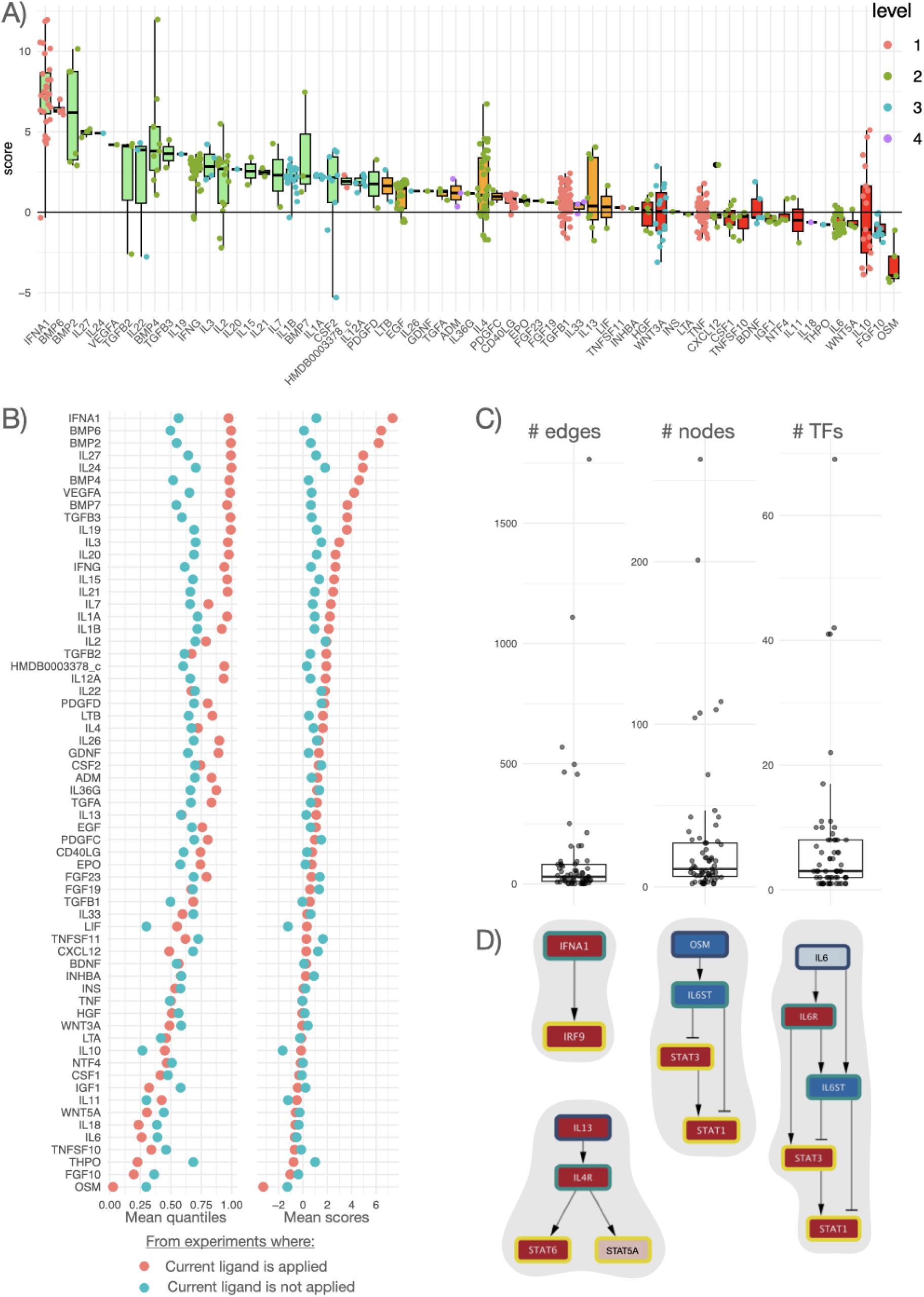
Estimation of the activity of ligands via MOON score on Cytosig data. A) MOON score of ligands on the corresponding Cytosig experiment. The x-axis represents the ligand, the y-axis represents the MOON score, therefore each dot represents the score of a given ligand MOON score in each experiment where it was applied. When a given ligand is applied in an experiment, its corresponding MOON score is expected to be positive. The color of the dots represents the number of steps between the ligand and the TFs in the PKN. The color of the box plot denotes whether the average MOON score for the corresponding ligands is above 1.7 (green), between 1.7 and 0.3 (orange) or below 0.3 (red). B) mean quantiles and scores of ligands in experiments where they were applied compared to experiments where they were not applied. C) Number of edges, number of nodes and number of downstream TF participating in the MOON score estimation of the ligands. Each dot represents a ligand. D) Schematic representation of several ligand scoring networks. Nodes filling in red indicate up-regulation of activity while blue represents down-regulation of activity, in a given experiment. Color of the node border represents the distance from TFs, from yellow (level 0, TF themselves) to dark blue (level 2, 2 steps upstream of the closest TF). Flat arrowheads represent inhibitory interactions, pointy arrowheads represent activatory interactions.

To evaluate the specificity of the ligand scores, we compared their average quantile (within a given contrast) in experiments where cells were treated with them to their average quantile in experiments where cells were treated with other ligands (Figure 2B). For 44 out of 63 unique ligands, average scores were higher in experiments where they were applied compared to experiments where they were not applied, while the 19 others were equal or lower compared to experiments where they were not applied. To facilitate the interpretation of the MOON scores, we extracted a subnetwork for each ligand that contains only the nodes that connect the ligand (upstream regulator) to the nodes of the downstream layers (the most downstream layer being TFs) that were used as targets during each iterative scoring round of MOON. This showed that the ligand scores were estimated from an average of seven TFs (Figure 2C) in the most downstream layer of each respective ligands. The complexity of the MOON scoring network across the ligands varied from very simple networks (2 nodes, 1 edge) to networks with > 1,000 of edges in some cases (max = 263 nodes, max = 1,766 edges, Figure 2C). Half of the scoring networks of ligands comprised no more than 30 edges, 11 nodes and 3 TFs. As an example, IFNA1 ligand had a very high MOON score on average (7.3). IFNA1 has a single TF, IRF9, as a direct downstream target, (Figure 2D). The MOON scores are defined from their most direct TF targets, to capture first/early effects; other TFs further downstream than IRF9 are not used for the computation of the MOON score. The high MOON scores indicate that treating cell lines with IFNA1 will consistently lead to the activation of IRF9.

The ligand OSM seemed to perform very poorly, with an average score of -3.25. OSM’s MOON score depends on the activity estimation of STAT3 and STAT1 (Figure 2D). We found that there was an error in the prior knowledge downstream of OSM, as IL6ST was annotated as an inhibitor of STAT3 and STAT1. IL6ST and OSM are in fact activators of JAK kinases (Mosly *et al*, 2023), explaining why OSM’s score was the opposite of what would be expected. This mistake was corrected in the version of Omnipath of 2024.03.19, and the average score of OSM in this new version is 4.4. This erroneous annotation also explained the seemingly poor MOON score estimation of the IL6 ligand (Figure 2D), which has an average score of 2.0 in the newer version.

An example of a seemingly random performer is IL13 (0.59 average score quantile in experiments where it’s applied and 0.59 as well where it’s not, Figure 2D), which had relatively heterogeneous MOON scores across experiments, despite having a positive score on average (IL13: mean = 1.06, SD = 2.26). IL13 is estimated mainly from the activity scores of STAT5A and STAT6 (Figure 2D), and the latter is a canonical outcome of IL13 (Rolling *et al*, 1996). While it has a high score in three experiments where it is applied (MOON score > 3), it has a negative score in 4 other experiments, leading to poor average score quantiles. The score did not correlate with either time after collection or cell type, however three out of four of the poorly scored experiments were from the same study (GSE43515). Additional examples with more complex structures are detailed in Supplementary Figure 2, showing cases where larger networks (e.g. TGFB1 and EGFR) may lead to lesser impact of prior knowledge errors and the impact of potentially context specific molecular interactions.

### 2.3 Mechanistic analysis of multi-omic factor weights

To demonstrate its use, we applied COSMOS+ to the NCI60 dataset, consisting of transcriptomics, proteomics and metabolomics across 58 cell lines (Su *et al*, 2011; Gholami *et al*, 2013) (Figure 3A). Preprocessing of the NCI-60 data yielded 6,000 most variable transcripts, 1,897 most variable proteins and 139 metabolites measured to be used as inputs for MOFA (see methods, Supplementary Figure S3). While 6,000 transcripts only represent a fraction of all the genes that were measured, they are enough to robustly estimate the activity of >100 transcription factors (Supplementary Figure S4A). Next, we ran MOFA while progressively constraining the maximum number of factors between 4 and 15, and assessed how robust the results were. We saw that at 9 factors and above, the optimal number of factors that MOFA chose didn’t change anymore, consistently settling on 9 factors, regardless of whether the upper factor limit was further increased. (Supplementary Figure S5). The MOFA model with 9 factors could reconstruct different percentages of the data variance for each type of omic data (Figure 3B). 60% of the variance was reconstructed for the RNA data, 19% for the metabolomic data and 23% for the proteomic data. The reconstructed variance is not much higher for the proteomic data compared to the metabolomic data, despite having more than 10 times more quantified features (1897 proteins vs 139 metabolites). This is likely due to the fact that, in general, metabolic abundances are more correlated between each other than protein abundances, due to the entangled metabolic reactions that connect them (Camacho *et al*, 2005; Saccenti *et al*, 2015; Steuer, 2006), making it easier to capture coordinated patterns despite having less features available to fit the MOFA model (Supplementary Figure S6). Factor 2 and 4 could explain variance in the data across all three omic views.

**Figure 3:**
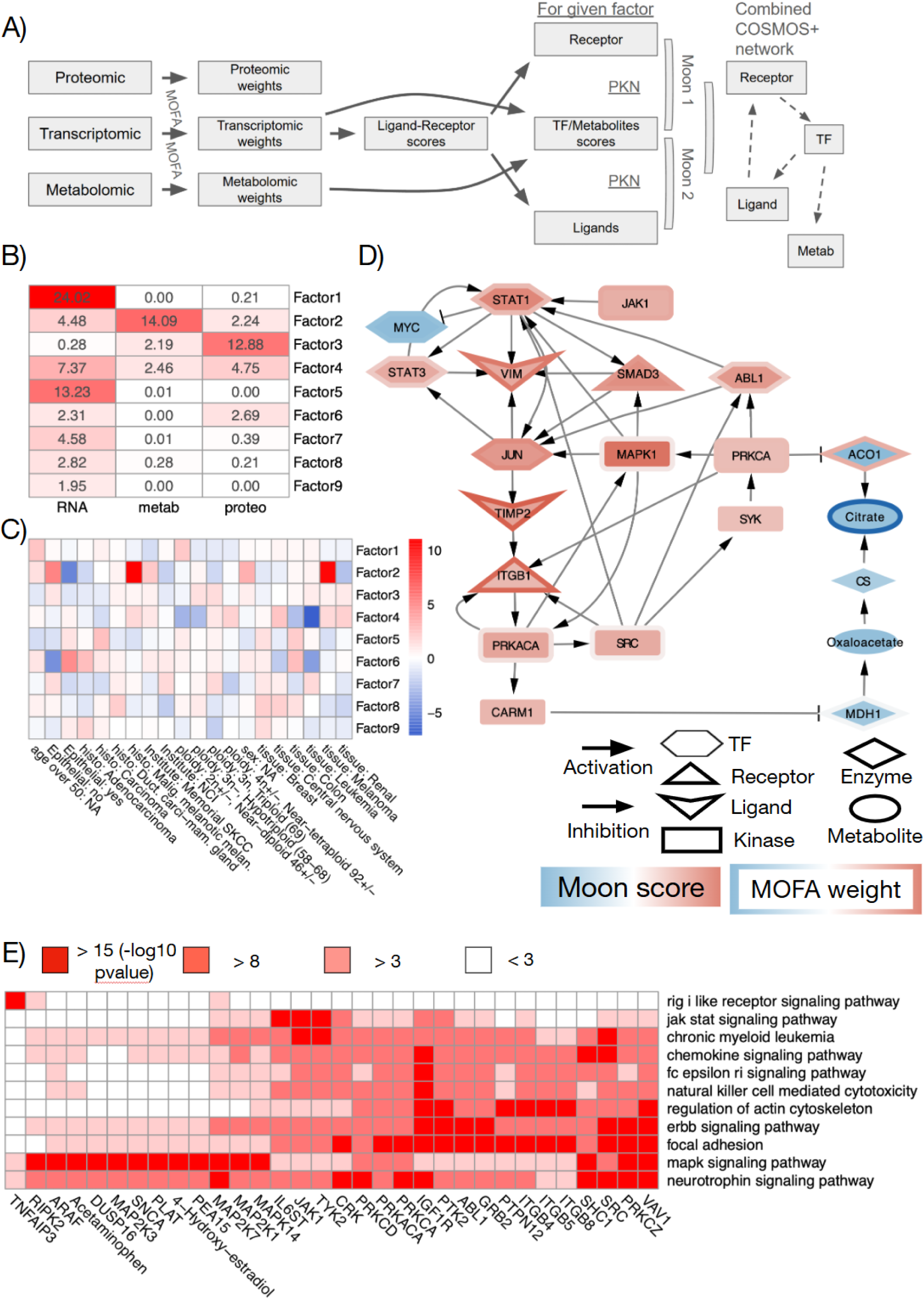
Mechanistic interpretation of a MOFA factor associated with blood cancer cell lines, such as MOLT-4 cell lines. A) Schematic representation of the data flow from NCI60 multi-omic data to COSMOS+ network B) Heatmap of the % of variance explained per omic view and per MOFA factor. **C**) Enrichment analysis of cell line tissue of origin metadata in each of the MOFA factors. Enrichment scores are computed using the ULM method of decoupleR (t-value of linear regression between the MOFA Z values in the background and each metadata class, such as tissue of origin, age of donor, pathology, etc..). C) MOON Network connecting the top deregulated TFs and LR interactions of factor 4 based on a signed directed prior knowledge network. D) Heatmap of the top results of the Pathway control analysis. We represent pathways that are significantly over-represented downstream of given nodes of the MOON thresholded network

We then checked which meta-data features of the NCI60 cell lines were enriched in each factor. MOFA decomposes the data into a Weight matrix and a Z matrix of cell line coordinates in the factor space. The latter can be analyzed to find meta-data features that are associated with the Z matrix coordinates. We used linear regressions (see methods) to model the coordinate of each cell line in the MOFA Z matrix as a function of each binarized clinical feature category (such as Tissue: renal = 0 or 1). Factor 4 was showing a significant negative association with samples of Leukemic origin. We also saw that factor 2 was significantly associated with a Melanoma origin, and negatively associated with an Epithelial origin (Figure 3C).

To get a general sense of the relationship between RNA and protein MOFA weights, we first explored how correlated the weights of transcripts and protein abundances were (Supplementary Figure S4B). The correlation between RNA and protein weights was close to 0 for factors 1, 3, 5, 7, 8 and 9. For factor 2, 4 and 6, the correlation was respectively 0.41, 0.42 and 0.29. This showed that the variance reconstructed for factor 2, 4 and 6 across the RNA and proteomic view is, at least partly, capturing the relationship between RNA and the corresponding protein abundance. Of note, the correlation between RNA and protein weights for those factors is of a similar magnitude as usual RNA/protein correlation at the level of bulk data (Shankavaram *et al*, 2007; Gry *et al*, 2009) (mean correlation for each cell-line across genes between RNA and protein value = 0.41 +- 0.04 s.d.; mean correlation for each gene across cell-lines between RNA and protein value = 0.29 +- 0.24 s.d.) (Supplementary Figure S7).

Then, we set to characterize mechanistically the factors using footprint and LR analysis methods. To do so, we used linear regressions to model RNA weights (Supplementary Figure S4C) as a function of their status as targets of each TF of the CollecTRI (Müller-Dott *et al*, 2023) database. This allowed us to obtain scores that represent how consistently high or low RNA weights of the transcriptional targets of given TFs compared to the background of RNA weights are. We can interpret such scores as TFs that are responsible for the transcriptional programs that are captured by given factors. The TF scores were especially significant for factor 4 (48 out of 284 TFs with p-values < 0.001; <u>Supplementary</u> <u>Figure S4</u>D). This indicates that the variance captured by factor 4 is consistent with our available prior knowledge about transcriptional regulations. The most expressed TF scores for factor 4 were members of the JAK-STAT, NFKB and hypoxic pathways (STAT1, NFKB, SMAD3, HIF1A). Since factor 4 was negatively associated with cells of Leukemic origins, this may indicate that those pathways are specifically less active in those cells than in cell lines of other origins. Then, we used decoupleR’s ULM function again to model RNA weights as a function of their status as members of ligand-receptor (LR) pairs. This allowed us to obtain LR scores that represent how consistently higher or lower the weights of transcripts coding for LR pairs compared to the weights of other transcripts in each factor are. Interestingly, factor 4 also showed the most consistency between RNA weights and prior knowledge about LR interactions (<u>Supplementary</u> <u>Figure S4</u>E). The most significant LR pairs of factor 4 were involved in processes such as cell adhesion (e.g. CD44, ITGBs, VIM, etc…).

To complement the TFs and LR interactions associated with factor weights, we used MOON to find network-level hypotheses supported by the RNA, protein and metabolite weights of the MOFA factors. We specifically focused on factor 4, since it captured variance at the level of all three omic views (RNA, proteomic and metabolomic), and was especially consistent with TF-target and LR interaction prior knowledge (Figure 3A, Supplementary Figure S4D,E).

The resulting MOON network of factor 4 shows which of the mechanistic interactions available in prior knowledge resources (e.g. the COSMOS PKN) are consistent with the feature weights of factor 4 (Figure 3D). For example, we can see that VIM has a high weight (0.7) in factor 4, and the transcription factors SMAD3, STAT1, STAT3 and JUN have many transcriptional targets that have a high weight in factor 4 (targets mean = 0.13, including VIM). Furthermore, we can also see that MAPK1 (MOON score = 3.2), PRKCA (MOON score = 2.4), ABL1 (MOON score = 2.8) and JAK1 (MOON score = 2.8) have all high MOON scores, which indicates that they are directly upstream of a high number of consistently up-regulated TFs (such as SMAD3, STAT1, STAT3 and JUN).

Complementarily, their corresponding transcripts also have high weights in factor 4 (MAPK1 = 65th quantile, PRKCA 92nd quantile, ABL1 = 86th quantile, JAK1 = 97th quantile of factor 4 weight). Taken together, factor 4 appears to be characterized by a consistent network of interactions connecting VIM as a downstream transcriptional target of SMAD3, STAT1, STAT3 and JUN, themselves regulated through post-translational regulations by MAPK1, PRKCA, ABL1 and JAK1. Other transcription factors, ligands and receptors also show activities and weights consistent with this signaling and transcriptional regulation, such as MYC, TIMP2 and ITGB1 (Figure 3D). MOON network also points at potential metabolic regulations: ITGB1 and MAK1 can activate PRKACA, which can itself activate CARM1, an inhibitor of the MDH1 metabolic enzyme through methylation. The inhibition of MDH1 could indirectly explain the downregulation of citrate (citrate = -0.38 factor 4 weight). Finally, PRKCA activation by SYK and SRC downstream of PRKACA can also lead to the inhibition of the citrate conversion activity of the metabolic enzyme ACO1 (Fillebeen *et al*, 2005; Pitula *et al*, 2004) (Figure 3D).

To find which biological processes are captured in the moon network, we perform a pathway over-representation analysis with sets of nodes down-stream of top deregulated MOON score nodes (absolute MOON score > 1.5) Then, we can represent pathways that are significantly over-represented downstream of given top scoring nodes of the MOON thresholded network in a heatmap (Figure 3E). Reassuringly, this analysis found expected control mechanisms, such as JAK1 or IL6ST very significantly controlling the JAK-STAT signaling pathway or ITGB (integrins) family members controlling focal adhesion. It also allows us to propose chemicals that can potentially control signaling pathways such as Acetaminophen or 4-hydroxy-oestradiol controlling the MAPK pathway.

### 2.4 Mechanistic analysis of breast cancer cell line multi-omics data in response to CDK inhibitors

We examined whether activities estimated by MOON can offer information regarding drug response and resistance mechanisms to cell cycle inhibitors in breast cancer. To address this, we created and analyzed a multi-omics perturbation dataset comprising transcriptomic and phosphoproteomic data. This dataset was structured to investigate resistance to cyclin-dependent kinase (CDK) 4 and 6 inhibitors, such as palbociclib—an approved oral CDK 4 and 6 inhibitor in breast cancer cell lines (refer to Table 1 and Methods). This dataset was appealing for two reasons. First it provided a high complexity multi-perturbation and multi-omics dataset well suited for COSMOS+. Second, the cell lines in the dataset could be compared to a closely related clinical breast cancer dataset (Paloma3 study) which we analyze with COSMOS in the subsequent section.

**Table 1:**
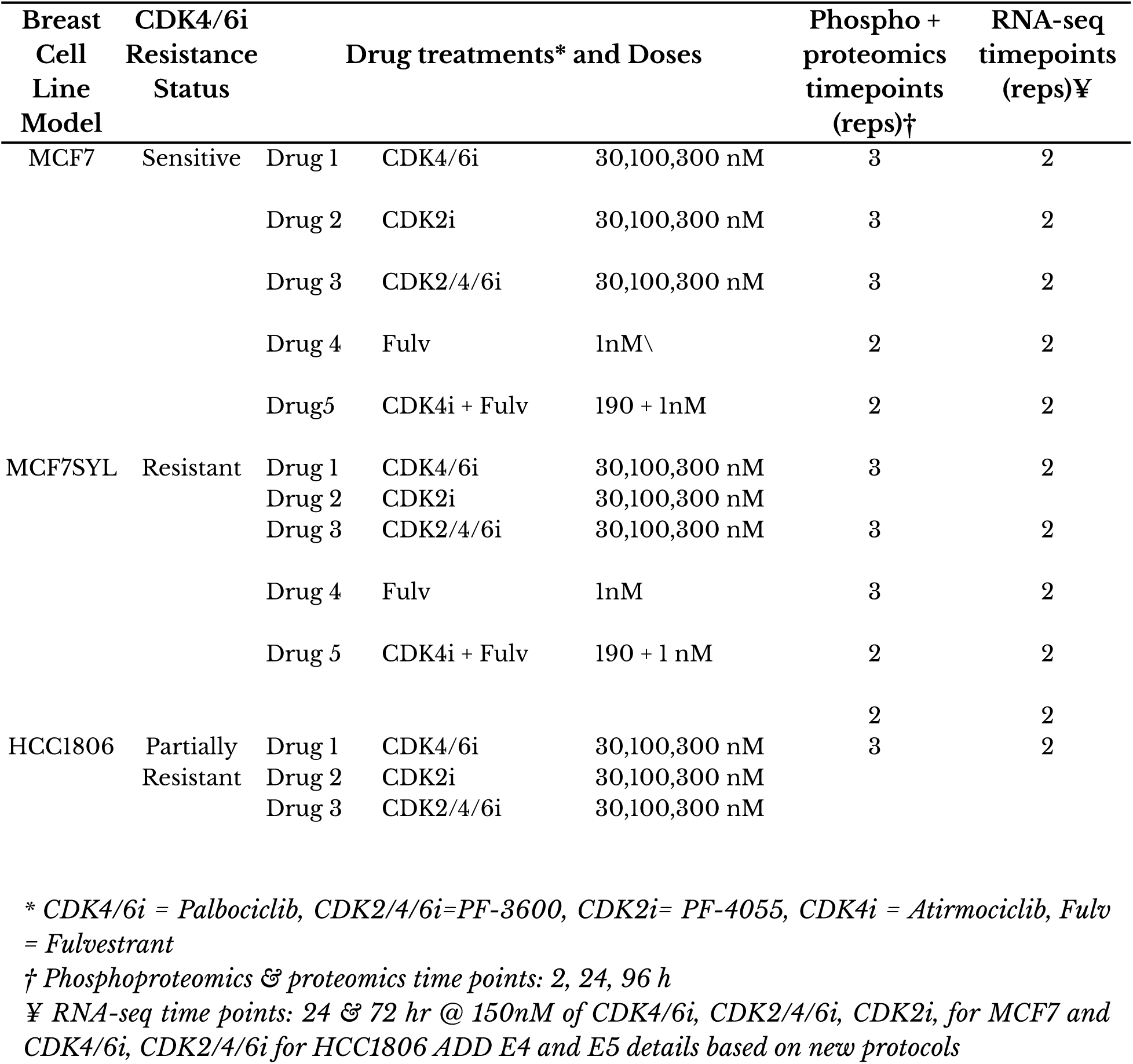
Summary of the cell-line RNA and phospho-proteomic dataset.

We used the COSMOS+ pipeline to explore potential mechanistic connections that could connect deregulated phosphosites regulators (e.g. kinase/phosphatases) and downstream transcription factors. Across all the conditions where we could match both RNA and phosphorylation measurements (refer to multi-omic time-point matching supplementary Text S1 and Supplementary Table S1), 22 differential analysis contrasts between the different treatments and matched time-point (early: 2-4 hours; medium: 24 hours; late: 72-96 hours) untreated controls could be performed. Series of differential analysis were performed rather than a multi-omic factor analysis in this context because the dataset was clearly structured and different conditions could be explicitly compared. Therefore, COSMOS+ was applied on each contrast individually. TF activity scores were obtained as previously described using the CollecTRI regulons and univariate linear models of decoupleR. To estimate kinase scores, we obtained a kinase-substrate network from Omnipath. However, very few targets of CDK2, 4 and 6 in the kinase-substrate network were detected in the phospho-proteomic dataset (65, 2 and 1 targets, respectively). To obtain more reliable kinase scores, we applied kinasePhos3 tool (Ma *et al*, 2023), a machine learning-based framework to infer specific kinase phosphorylation site targets from protein sequences, to infer kinase-substrate relationships on all human Swissprot protein sequences. The combination of the Omnipath curated kinase-substrate network (26090 curated kinase-substrate relationships) with the kinasePhos3 network (2533817 inferred kinase-substrate relationships) increased the number of detected targets of CDK2, 4 and 6 to 820, 362 and 380 targets, respectively. It was used with the univariate linear model method of decoupleR to estimate kinase activity scores. Then, the Omnipath prior knowledge network of protein-protein interactions was scored with the MOON algorithm using TFs as downstream inputs and kinase activities as upstream inputs.

In the resulting scored network, we first assessed if the activity of canonical CDK pathway members (CDK2,4, and 6, RB1 and E2F4) (Wang *et al*, 2024) was estimated in an expected manner following CDK4/6 inhibition with CDK4/6i at 24 hours in sensitive (MCF7) and partially resistant (HCC1806) cell lines. CDK2,4,6 and E2F4 were consistently down-regulated in the sensitive cell line MCF7 while RB1 was up-regulated, but they were not or only mildly affected in the resistant cell line HCC1806 (CDK2 = -2.25/1.41; CDK4 = -1.61/0.59; CDK6 =-0.67/1.56; RB1 = 3.85/-0.31 E2F4 = -18.35/2.07 MOON scores in MCF7/HCC1806, respectively). As this dataset was designed to explore mechanisms related to CDK2 mediated resistance, we also assessed the activity differences in MCF7SYL (fully resistant cell line) treated with the CDK4/6 inhibitor versus with a CDK2/4/6 inhibitor. As expected, at the early timepoints (2h phospho/4h RNA), CDK2 activity was more down-regulated following CDK2/4/6 inhibitor treatment compared to CDK4/6 inhibitor treatment (CDK2 = -2.1 MOON score). CDK2/4/6 inhibitor treatment also appeared to inhibit the cell cycle progression as E2F1,2,3 and 4 were the top down-regulated activities (E2F1 = -16.2; E2F3 = -14.8, E2F4= -12.4; E2F2 = -9.1 MOON scores). The 5^th^ most down-regulated activity was that of FOXM1, a direct target of CDK2. These results suggest that this dataset, when analyzed with COSMOS+, allows to explore both differences between cell lines and between target-specific inhibitors.

While each contrast between conditions could be explicitly computed, the high number of comparisons still made it difficult to intuitively interpret this series of contrasts in order to extract high level patterns, such as general mechanisms of resistance. To extract such patterns of resistance to CDK inhibitors rather than exploring each contrast one by one, we performed a PCA on the resulting MOON scores matrix (across the 22 contrasts; Figure 4B). The first Principal Component (PC1) was positively associated with MCF7 cells (t-value = 3.45) and negatively associated with both resistant cell lines (MCF7SYL t-value = -1.29 and HCC1806 t-value = -2.21), suggesting an association with general mechanisms of resistance shared across both resistant cell lines. Furthermore, the first component was not confounded by CDK4/6i, CDK2/4/6i or the fulvestrant + CDK4i combo treatment or time points (that is, samples treated with e.g. CDK4/6i or CDK2/4/6i appeared to be distributed evenly across PC1), which would indicate that this factor represents a general separation of MCF7 versus MCF7SYL and HCC1806 drug responses (Figure 4D). Due to the separation of resistant and sensitive cell lines on this component, we focused our analysis further on the factor loadings of PC1, which captured differentially active (based on MOON scores) features between the sensitive and resistant cell lines.

**Figure 4:**
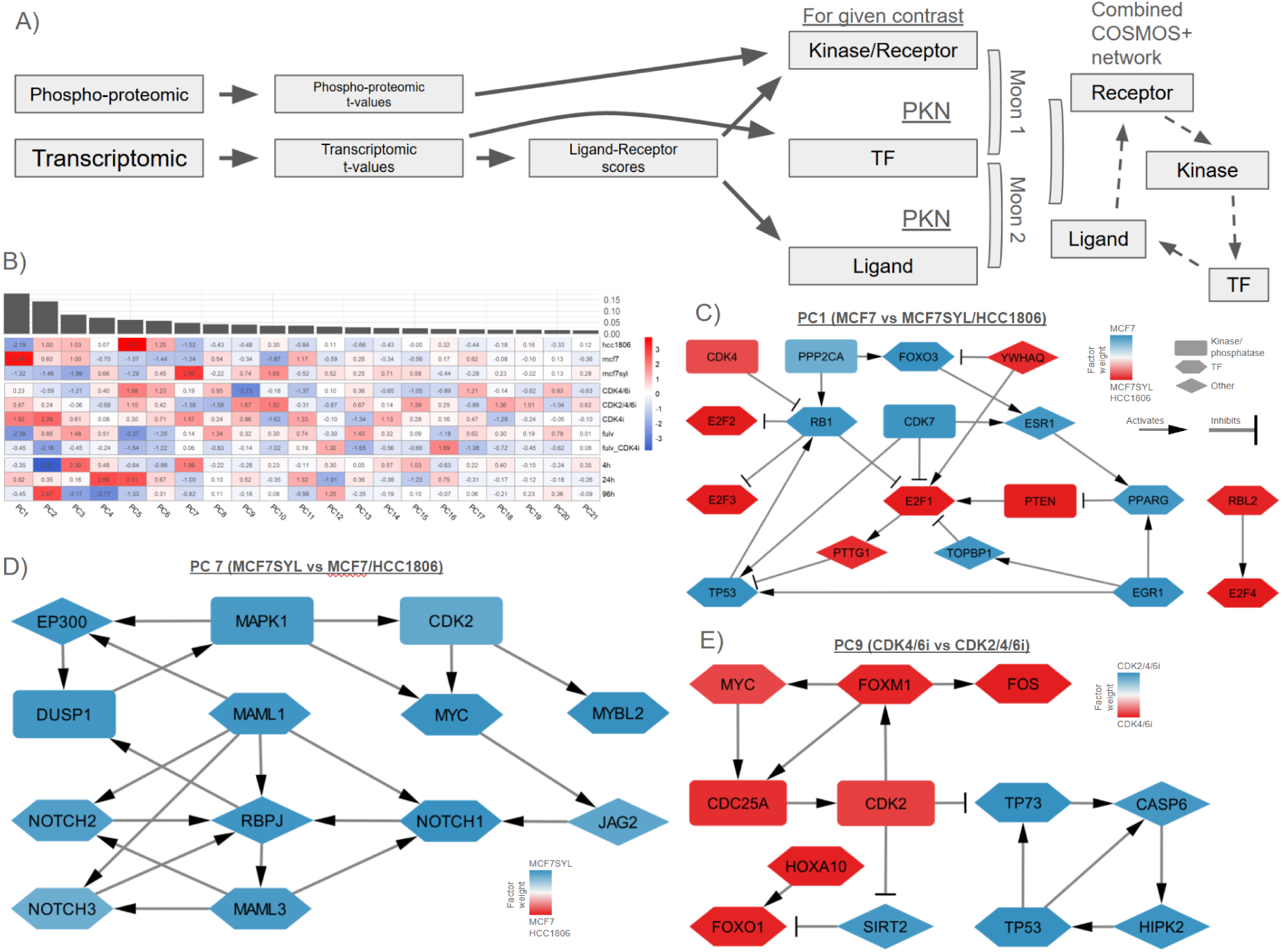
Mechanistic analysis of multi-omic time series dataset of breast cancer cell line response to several CDK inhibitors. A) Schematic representation of the data flow from breast cancer cell line experiemtal data to COSMOS+ network. B) T-values of linear modeling of PCA scores as a function of meta-data feature (cell line, drug, timepoint). The PCA was performed on a MOON score matrix estimated from the differential analysis results of all the comparisons that were made (e.g. MCF7 treated with CDK4/6i at 24 hours against control MCF7) and could be overlapped between phospho and RNA data. Therefore, each value can be interpreted as the association of a given meta-data feature with a given PCA component, while the PCA space itself is a dimensionality reduction of all the MOON scores calculated across all comparisons. C) Network illustration of a select few top PCA loadings (computed from MOON scores) from PC1. MCF7 (sensitive) is well separated from MC7SYL and HCC1806 (resistant) on this component. It highlights the differential regulation of E2Fs. Red/Blue color indicates positive/negative association of PCA weights (based on MOON scores) with MCF7 (blue) and HCC1806/MCF7SYL (red) cell lines D) Network illustration of a select few top PCA loadings (computed from MOON scores) from PC7. MCF7SYL (resistant) is well separated from MC7 (sensitive) and HCC1806 (resistant) on this component. It highlights the crosstalk of CDK2 and NOTCH signaling associated with this factor. Red/Blue color indicates positive/negative association of PCA weights (based on MOON scores) with MCFSYL (blue) or MCF7 and HCC1806 (red, none selected in this specific subnetwork) cell lines. E) Network illustration of a select few top PCA loadings (computed from MOON scores) from PC9. CDK4/6i (CDK4/6i) is well separated from CDK2/4/6i (CDK2/4/6i) on this component. It highlights the differential regulation of CDK2 and its neighbors. Red/Blue color indicates positive/negative association of PCA weights (based on MOON scores) with CDK2/4/6i (blue) or CDK2/4i (red).

To identify potential mechanisms associated with PC1, we filtered the original prior knowledge network to keep only consistent interactions between the top MOON features (that is, nodes that have loading signs that are consistent with the sign of the edges of the prior knowledge network that connect them together, and are in the 90th percentile (90p nodes) of factor weights or in the 66th percentile (66p nodes) of factor weights but connect two 90p nodes together, see Methods). This yielded a network composed of 229 nodes and 569 edges. Figure 4E shows a subset of this network focusing on upstream regulators of the E2Fs transcription factors, which are important mediators of CDK activities through RB1. The network shows a potential feedback loop between E2F1, PTTG1, TP53 and RB1. CDK7 may further activate TP53 while inhibiting E2F1. ESR1 and EGR1 could be responsible for PPARG activation in MCF7, leading to an inhibition of PTEN and E2F1. YWHAQ may also be involved in the activation of E2F1 (through its stabilization) while inhibiting FOXO3 in MCF7SYL and HCC1806 cell lines. This network thus summarizes various potentially differentially regulated mechanisms following treatment between sensitive MCF7 cells and the resistant MCF7SYL and HCC1806.

While PC1 was the only component where the sensitive cells (MCF7) were clearly separated from both resistant cells, other components with strong association with one or the other cell line were PC5 and PC7 (Figure 4B). PC7 was associated with the acquired resistance cell line MCF7SYL. Filtering the original prior knowledge network with the top MOON features of PC7 yielded a network of 239 nodes and 457 edges. CDK2 has a positive factor weight in the network, indicating a higher activity in MCF7SYL following CDKi treatment, consistent with the hypothesis that the acquired resistance of MCF7SYL to CDK4/6 inhibition is mediated by CDK2 compensation. One of the top weights in this factor is MYC, directly downstream of CDK2 activation. Furthermore, MYC regulates the expression of JAG2, a ligand of NOTCH1, a critical regulator of cell proliferation. NOTCH1,2, and 3, as well as MALM1,3, RBPJ and EP300 are all connected downstream of MYC in this network. EP300 itself can activate DUSP1, potentially cycling back to CDK2 through MAPK1 (Figure 4D). Therefore, the network associated with MCF7SYL recapitulates well the CDK2 mediated acquired resistance to treatment while highlighting a potential positive feedback loop through MYC and the NOTCH pathway.

In addition to factors associated with differences between cell lines, PC9 was the component with the highest difference of association between the CDK4/6 inhibitor and CDK2/4/6 inhibitor (Figure 4B). We filtered the original prior knowledge network to keep the top associated factor weights. The resulting network contained 197 nodes and 378 edges. It shows CDK2 as one of the top negatively associated activities with this factor. This is expected since the inhibition of CDK2 is the main difference between the two treatments that are associated with each extremity of this factor. Focusing on the close neighborhood of CDK2 shows a potential positive feedback loop around CDK2 mediated by FOXM1, MYC and CDC25A. The FOS and FOXO1 TFs are also potentially deregulated following the specific inhibition of CDK2, as well as TP53 and TP73 in a positive feedback loop through CAPS6 and HIPK2, important mediators of cell death (Figure 4H). Therefore, this network represents potential mechanisms that could connect the specific inhibition of CDK2 with increased apoptosis through activation of CASP6 and HIPK2 while down-regulating FOX and FOS TFs.

### 2.5 Comparison of deregulated features in resistant cells with genome wide CRISPR KO sensitization assays

Mechanistic hypotheses help explore the potential signaling response from the three cell lines (MCF7, MCF7SYL and HCC1806) to various CDK inhibitors. We sought to evaluate their accuracy and functional relevance with respect to resistance to treatment by investigating whether feature weights of the PCA on MOON scores were associated with mechanisms of resistance to the CDK inhibitors CDK4/6i (CDK4/6) and CDK2/4/6i (CDK2/4/6). For this purpose, a genome wide crispR KO essentiality/sensitization dataset was generated with MCF7 and HCC1806 cell lines in control, palbociclib and pf3600 treatment conditions.

First, we assessed how comparable the gene essentiality or sensitization signatures were between the two cell lines. Hierarchical clustering on the crispR KO profiles showed that their differences were driven more by condition (e.g. control or sensitization assay) than by cell line (Figure 5A). This indicated that resistant and sensitive cell lines still appeared to be sharing many essential genes as well as sensitization targets. Next, to assess the agreement between the MOON scores and the crispR sensitization scores, we computed the correlation between the features weights of the first factor of the MOON score PCA, which captures differences in regulation between the resistant and sensitive cell lines, (see section 2.4) and the minimum KO sensitization scores of the HCC1806 treated with CDK4/6i (CDK4/6 inhibitor) or CDK2/4/6i (CDK2/4/6 inhibitor, see Methods). The Pearson correlation coefficient was small but significant (R = 0.06, p-value = 0.02). The positive sign of the correlation coefficient can be interpreted as an agreement between genes that have negative KO sensitization scores, e.g. are sensitizing cells to treatment and MOON scores that are higher in HCC1806 cells compared to MCF7. Such genes are subsequently referred to as MOON/essentiality consistent genes. While small, the correlation coefficients between the feature weights of each of the components of the PCA and HCC1806 KO sensitization scores seemed to show an association with the separation between MCF7 and HCC1806 samples on each individual factor of the PCA (Pearson R = -0.52, p-value = 0.02), but not with MCF7 (Pearson R = -0.009, p-value = 0.96) (Figure 5B). Thus, while small, the correlation patterns are significantly consistent across every factor for the resistant cell line but not for the sensitive one. This agrees with the expectation that PCA factors that are specifically associated with resistant and sensitive cell separation would potentially be associated with resistance mechanisms that can be sensitization targets, but only in resistant cells (HCC1806) and not cells that are already sensitive (MCF7).

**Figure 5:**
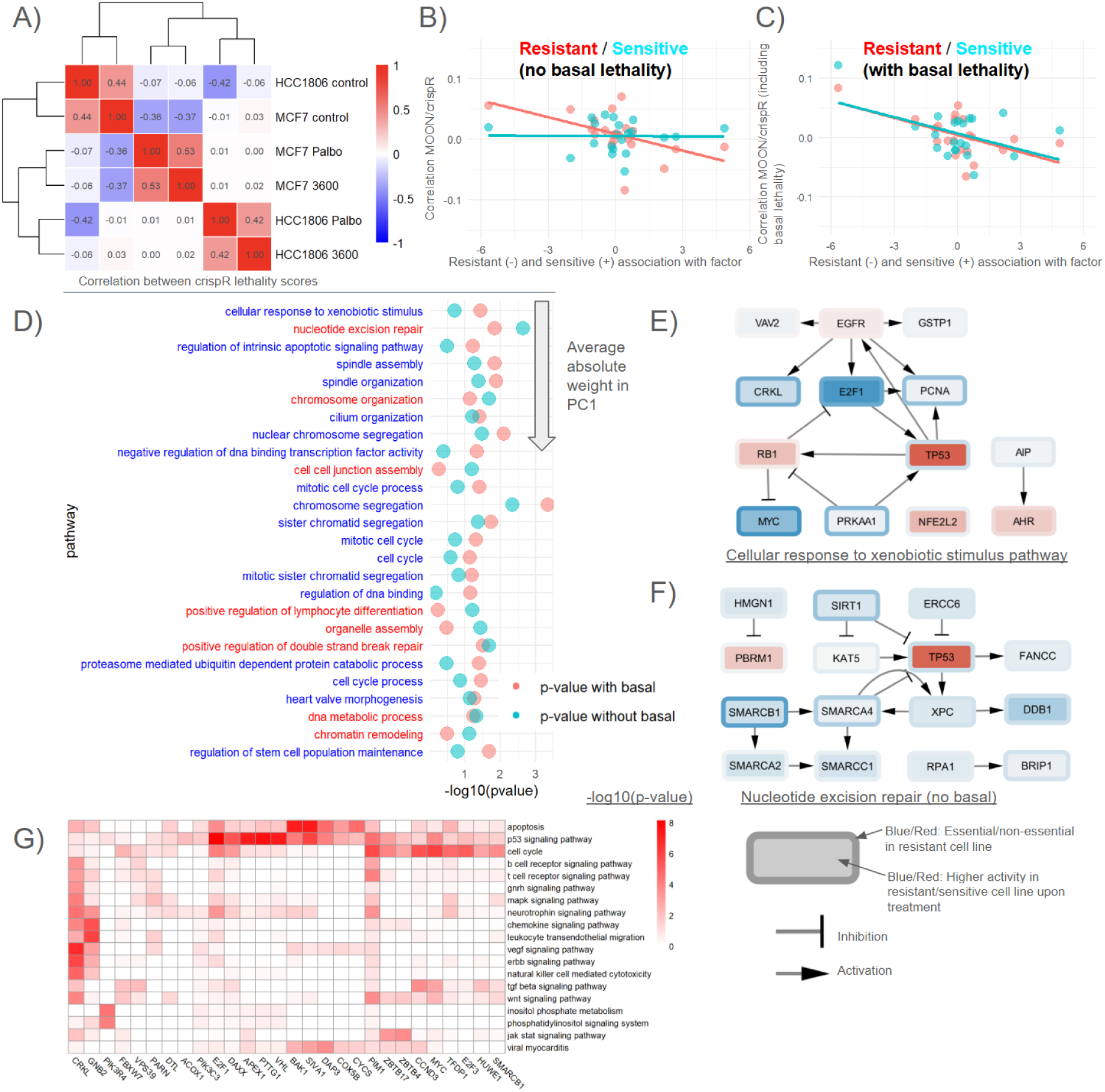
Joint analysis of MOON activity scores and crispR lethality. A) Correlation and hierarchical clustering between crispR genome wide KO lethality scores (Mageck z value) in 6 conditions: 2 cell lines, MCF7 (sensitive) and HCC1806 (resistant) and 3 treatments (control, Palbo and CDK2/4/6i). B) Each dot corresponds to a principal component of the MOON score matrix PCA (see <u>2.4</u>). X axis is the difference between the resistant cell (HCC1806) and the sensitive (MCF7) t-value of association with the respective PCs. Y axis is the correlation between the loadings (based on MOON scores) of a given PC and the crispR lethality scores (minimum value between the two drug treatmentS) in HCC1806 (red) or MCF7 (blue). Expectation is that correlation should be high when the difference is negative (resistant negative association, thus negative loading means high in resistance, thus they should have negative crispR score as well), and vice-versa, only in the resistant cell line. Correlation coefficients between the feature weights of each of the components of the PCA and HCC1806 KO sensitization scores for HCC1806 samples on each individual factor of the PCA (Pearson R = -0.52, p-value = 0.02), and MCF7 (Pearson R = -0.009, p-value = 0.96) C) Same as B, but here the crispR score is the minimum between the two drugs AND the control. The rational being that genes that are already lethal in control may not show up as more lethal in treatments. In this case, the correlation is significant for resistant (cor and sensitive cell lines. This indicates that the negative correlation in B was indeed probably lacking statistical power and that the basal lethal genes are confounding the sensitizing effect to drugs of the crispR KO. Thus, there seems to indeed be a significant correlation between the MOON scores of the drug perturbation and the lethality of gene crispR KO. Correlation between the MOON PCA weights/crispR scores correlations and the separation between MCF7 and HCC1806 samples on each individual factor of the PCA for HCC1806 (Pearson R = -0.55, p-value = 0.01) and MCF7 cells (Pearson R = -0.46, p-value = 0.04) D) Top pathways that show agreement between moon score difference (resistant - sensitive) and crispR lethality to treatment E) Network representation of the cellular response to xenobiotic stimulus pathway with PC1 scores (node fill) and crispR score (outer node border) including basal essentiality. F) Network representation of the nucleotide excision repair pathway with PC1 scores (node fill) and crispR score (outer node border) excluding basal essentiality. G) Pathway control analysis of the top genes where MOON weights from PC1 are in agreement with crispR scores (PC1 weight < -1 and crispR score < -2). Here only pathways where at least one gene significantly controls it (p-val < 0.001) and only genes that significantly control at least one pathway (p-val < 0.05) are shown.

To further characterize how MOON score PCA feature weights and crispR KO scores agree or disagree, we first considered genes that had a KO sensitization score of -2 at least as true positives, that is genes that are driving resistance to CDK4/6i or CDK2/4/6i. We could then compute the area under the precision curve (AUPRC) for genes that have negative weights in PC1 (that is, genes that are more active in the resistant cell line after treatment with CDK4/6i and CDK2/4/6i). The AUPRC is favored in this case to the area under the receiving operator curve (AUROC) due to 1) the imbalance of the ratio of true positive and true negative (only 7% of true positive) and 2) precision is a usual metric for a model that we assume does not inherently capture the full complexity of the underlying biological mechanism, and merely contextualize a part of it. The AUPRC computed in this manner was 0.11, against a random baseline AUPRC of 0.07. At a given significance threshold of -2 as well on the normalized PC1 weight (that is 2 standard deviations below the average of the PC1 weights), the probability of a gene with a significant weight also having a negative crispR sensitization score is 0.85, against a baseline probability of 0.72 (binomial p-value = 0.046). On the other hand, the probability of observing a gene with a negative PC1 weight among genes that have a crispR z-score below -2 is 0.59, against a baseline probability of 0.55 (binomial p-value = 0.26). Therefore, while the AUPRC is higher than a random baseline, this shows that the precision of the PC1 weight is better than the recall to capture sensitization drivers.

To account for the potential confounding effect of the general essentiality of genes that could mask their sensitizing effect when knocked out, we repeated the previous analysis but this time by including the general essentiality crispR scores (MageCK scores without treatments) in the correlation coefficient calculation. This time, the significance of the association between the MOON PCA weights/crispR scores correlations and the separation between MCF7 and HCC1806 samples on each individual factor of the PCA increase for both HCC1806 (Pearson R = -0.55, p-value = 0.01) and MCF7 cells (Pearson R = -0.46, p-value = 0.04) (Figure 5C). Therefore, the MOON/essentiality consistent genes, that is genes that have higher moon scores in the resistant cell line HCC1806 compared to the sensitive cell lines MCF7 when treated with CDK2/4/6i (CDK 2/4/6 inhibitor), seem to often be essential genes.

To further explore the biology of genes that are both essential/sensitizing and more active in resistant cells treated with CDK inhibitors, a pathway over-representation analysis was performed using genes that were showing a sign agreement between crispR scores and MOON PCA feature weights on PC1 as a significant set, and the genes that have a sign disagreement as background. This indicated that pathways related to chromosome segregation, cell cycle and apoptosis seemed specifically enriched with genes that have both up-regulated moon scores in the resistant cell (i.e. negative weight in PC1) and negative (i.e. sensitization) crispR scores (Figure 5D). The pathway that had the most extreme weights on average in the first component of the PCA while having an over-representation of MOON/essentiality consistent genes (p-value <= 0.05) was “cellular response to xenobiotic stimulus”. Among the pathways containing genes with the most extreme weights on average in PC1, several were more significantly enriched with MOON/essentiality consistent genes not including the basal essentiality, such as “nucleotide excision repair”, “chromosome organization” or “cell cell junction assembly”.

To better understand the mechanisms underlying such pathways, we generated a network representation for the “cellular response to xenobiotic stimulus” pathway members (Figure 5E) by extracting a subset of the prior knowledge network containing only genes belonging to this pathway. This showed that the pathway contains a crosstalk between E2F1 and MYC, which are both highly ranked in terms of negative weight in PC1 while having a negative crispR score (including minimum between general essentiality and sensitization) lower than -3.8 and -4.8, respectively. We also represented a subset of the “nucleotide excision repair” pathway in Figure 5F, this time with crispR score only considering the minimum sensitization score. This showed that this pathway contained a crosstalk between the SMARC protein family (SMARCB1, SMARCA2, SMARCA4, SMARCC1), XPC and DDB1, which all had negative weights in PC1 while also having negative crispR sensitization scores. In both subnetworks, TP53 was present with a very positive PC1 score (i.e. biased toward the sensitive cell line), while still having a negative crispR score (sensitizing the resistant cell to treatment). The score of TP53 in PC1 was also incoherent with several of its neighbors in these pathways (e.g. PCNA or FANCC).

While visualizing pathways that contain genes that agree between MOON scores and crispR scores helps identify relevant molecular mechanisms, we also wanted to assess which pathways were specifically deregulated under the influence of genes that had both the top negative PC1 weights and negative crispR scores. We ran a pathway control analysis (see <u>methods</u> and section 2.3) for those genes specifically, which showed E2F1 controlled deregulated pathways (associated with PC1) such as cell cycle, apoptosis, P53 signaling pathway and neurotrophin signaling pathway (Figure 5G). Other examples of such controls were GNB2 and the chemokine signaling pathway and leukocyte transendothelial migration, PIK3R4 controlling the inositol phosphate metabolism, PIM1 controlling the WNT signaling pathway and T cell receptor signaling pathway and CRKL controlling the ERBB pathway (Natsume *et al*, 2012; Li *et al*, 2023).

### 2.6 Patient specific MOON signatures

Cell line models are useful to help identify potential mechanisms of resistance to CDK inhibition, however it is important to evaluate how such mechanisms translate in patients and if COSMOS+ could bring complementary information to gene expression alone with respect to prognostic marker identification.

To explore the clinical application of COSMOS+ and benchmark with existing standard methods, we first reproduced the COX analysis of (Turner *et al*, 2019) on the Paloma3 RNA expression cohort (2 treatment arms, CDK4/6i + Fulvestrant and Fulvestrant + Placebo, Figure 6A). The COX survival analysis identified the same markers associated with worse response (according to progression free survival) in the treatment arm (Figure 6B). COSMOS+ was then used to estimate activity score profiles at a single patient level (each patient RNA data normalized across the cohort was used as an input for TF activity estimations, which then were used to estimate MOON scores) in the Paloma3 cohort. The resulting MOON score matrix could then be used as input instead of RNA counts for the COX survival analysis, identifying markers associated at the level of their estimated activities instead of their expression (Figure 6B). CDK2 and RB1 MOON scores were both found to be significantly associated with worse response in the treatment arm, but not at their expression level. Furthermore, RB1 coefficient relative direction is reversed between its expression and MOON score. A lower MOON score of RB1 in patients is significantly associated with worse patient response in the treatment arm, while a high expression was marginally associated with worse patient response (MOON interaction p-value = 0.008, RNA interaction p-value = 0.12). Since RB1 is a known inhibited target of CDK4 and 6 as well as a tumor suppressor (Knudsen *et al*, 2019), its MOON score direction is more consistent with expectation than its expression.

**Figure 6:**
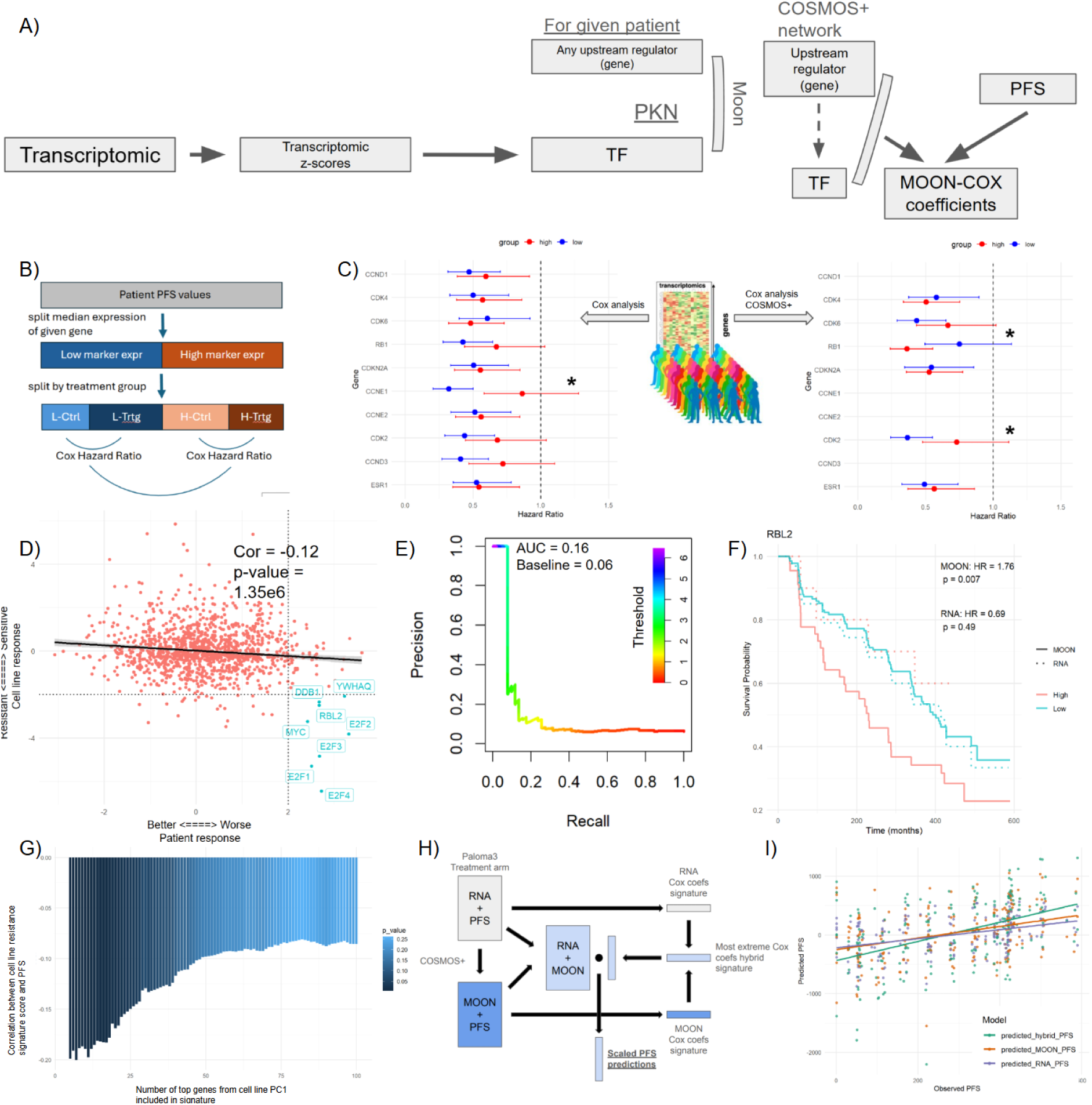
Clinical analysis of breast cancer treatment resistance with MOON mechanistic hypothesis generation. A) Schematic representation of the COSMOS+ pipeline followed by Cox survival analysis with the patient basal transcriptomic data. B) Schematic representation of the patient cohort splits to perform the Cox Hazard ratio computation with gene expression. For a given gene, the entire cohort is split between high and low expression (based on median expression). Then, each sub-group is further split between control and treatment arms. A Cox hazard ratio is then computed for each high/low expression group between treatment and control. Then, the significance of the difference between the Cox hazard ratio (control vs treatment) of high and low expression groups is estimated to determine if a gene is specifically associated with poor response in the treatment arm compared to the control arm. C) Cox hazard ratio confidence intervals for the gene that were part of the original signature of Turner et al. 2019. On the left, the Cox hazard ratios are computed based on gene expression values. On the right, the Cox hazard ratios are computed instead based on the patient specific MOON scores (computed through the COSMOS+ pipeline). Stars represent genes that have significantly (p-value < 0.05) different hazard ratio between the high and low expression group. D) Scatter plot between Cox coefficients (high/low activity score in treatment arm only) and PC1 (associated with difference between resistant and sensitive cell-line response) weights. For a given gene, higher Cox coefficient means that the MOON score is associated with worse patient response, while lower PC1 weight means that the MOON score is higher in the resistant cell line treated with CDK inhibitors than in the sensitive cell line. E) Precision-recall curve using the genes with MOON score-based cox coefficients greater than 2 (higher coefficient represent association with worse treatment response) as true positives and then computing the AUPRC of the PC1 loadings. F) Cox survival analysis curve for RBL2 in the treatment arm of the Paloma3 cohort. The high/low groups are defined based on MOON scores (full lines) or gene expression values (RNA, dashed lines). G) Correlation coefficient between patient resistance scores based on cell line derived signatures and observed PFS. The patient resistance score is computed with ULM where the input data is the patient specific MOON scores and the signature is defined as the top N lowest weight genes in the PC1 (associated with difference between resistant and sensitive cell-line response). We computed the coefficients for signatures comprising between the top 5 and top 100 lowest weighted genes of PC1. H) Schematic representation of the Cox coefficient signature based PFS prediction. For each gene, the Cox coefficient is computed between either high/low relative expression or high/low MOON score. Then, the Cox coefficients are used as a signature to compute a scaled PFS value by calculating the dot product between the Cox coefficient signature and the patient expression, for each patient, with either the RNA or MOON based Cox signature. Then, for each gene, the most extreme coefficient is selected between RNA and MOON score based coefficients to build an hybrid RNA/MOON signature that can also be used to estimate a scaled PFS value in a similar manner. I) Scatterplot and regression line between measured and scaled PFS values for each Cox coefficient signature (RNA, MOON score or hybrid based).

We also checked if the markers that were identified as differentially active between the resistant (HCC1806) and sensitive (MCF7) cell lines when treated with CDK4/6i or CDK2/4/6i (PC1 loadings) were also associated with worse outcomes in the treatment arm of the Paloma3 cohort. We focused on the treatment arm, where patients received CDK4/6i. The correlation between the cell line PC1 loadings and MOON score-based cox coefficients was significantly negative (Pearson correlation coefficient = -0.12, p-val = 1.35*10^-6^, Figure 6C). This indicates that genes that are more active in the resistant cell line compared to the sensitive one are also more active in patients with worse response to treatment. We also performed a precision-recall analysis by setting the genes with MOON score based cox coefficients greater than 2 (higher coefficient represent association with worse treatment response) as true positives and computing the AUPRC of the PC1 loadings. The AUPRC was 0.17, against a random baseline of 0.06. The precision is again high among the top PC1 loading, but the recall remains poor (for example, there are 7 true positives among the top 20 PC1 scores, which constitute a 5.83 fold increase compared to the random baseline).

One of the top MOON score markers for poor response in the Paloma3 cohort treatment arm was YWHAQ, also identified in the cell line models among the top differently active genes between sensitive and resistant cells. While YWHAQ expression wasn’t measured in the Paloma3 gene panel, COSMOS+ could still find such an association as the activity of YWHAQ is estimated from its downstream targets, rather than its actual expression (Figure 6D). RBL2 was another top consistent MOON marker between the cell line and patient response (Figure 6E). However, its expression doesn’t seem to be associated with PFS in the treatment arm, unlike its corresponding MOON score. Therefore, these further demonstrate the complementarity of information that can be gained by taking into account the activity of genes based on meta-footprint compared to their expression alone.

We also evaluated if the PC1 loadings from the cell line data could be used as a signature to predict poor patient response. Therefore, we selected the top n most negative loadings (n between 5 and 100) as a resistance signature and scored each patient accordingly using decoupleR’s ULM (using patient MOON scores as input data and the resistance signature as gene set). The best correlation between the patient resistance scores and actual PFS was observed in the range of 5 to 20 genes in the signature (Pearson correlation coefficient between -0.2 and -0.18, p-value between 5*10^−3^ and 2*10^-2^, Figure 6F). Beyond 20 genes, the absolute correlation coefficient decreases steadily until 55 genes, where it stabilized at around 0.08. This showed that there is a significant amount of information that can be extracted from a sensitive/resistance treatment response cell line model to inform patient response.

Finally, to further explore the complementarity of MOON scores with expression value, we created a hybrid Cox coefficient signature by combining the most extreme cox coefficients computed from the RNA data on the one hand and from MOON scores on the other hand (Figure 6G). That is, for each gene, the cox coefficient that was the most extreme between MOON and RNA was included. We then evaluated the ability of such a signature to estimate patient outcome by multiplying the hybrid Cox coefficient signature with the corresponding hybrid patient cohort of MOON scores and RNA measurements. The resulting vector essentially corresponds to scaled PFS predictions. The scaled PFS prediction was compared with the actual PFS for the hybrid signature, as well as with a signature based on RNA or MOON scores alone. The correlation for the hybrid signature PFS prediction was 0.45, while the RNA alone and MOON scores alone were 0.4 and 0.35, respectively (Figure 6H). This further supports that both MOON score and RNA values can bring complementary information to inform us about gene regulation events and processes that may be associated with better or worse outcomes in patients.

## 3. Discussion

In this study, we introduced COSMOS+, a flexible method to integrate multi-omic data and prior knowledge networks to extract interpretable mechanistic insights from complex experiments. COSMOS+ includes a novel and efficient network scoring procedure, meta-footprint (MOON), that can be used downstream of factor analysis. We show how MOON can identify biological mechanisms underlying cancer treatment resistance. COSMOS+ also introduces additional updates over the first version of COSMOS, such as a compression algorithm to remove redundant paths in the network and a method to highlight pathway control mechanisms. We further curated the metabolic reaction network used to build the prior knowledge network of COSMOS and refactored its building procedure within the Omnipath. All those updates enable the use of COSMOS+ across a wider range of applications with enhanced interpretation, including the ability to give insights at the level of single samples projected in factor spaces. We applied COSMOS+ on the Cytosig perturbation cohort transcriptomics dataset and the public NCI60 cell line multi-omic dataset to benchmark and demonstrate the potential of this new pipeline for multi-omic analysis. Then, we applied COSMOS+ on a novel multi-omic time course dataset of breast cancer cell lines treated with various CDK inhibitors to explore molecular response to CDK inhibition in resistant and sensitive cell lines. Finally, we used COSMOS+ on the PALOMA3 breast cancer patient cohort transcriptomic datasets to compare the breast cancer cell line analysis results with respect to patient outcome. These analyses demonstrate the flexibility and efficiency of COSMOS+ to generate mechanistic hypotheses across multi-omic layers and in diverse contexts. These results also illustrate how mechanistic hypotheses generated in the context of pre-clinical cell-line models can translate into a clinical setting and potentially help predict treatment efficacy as well as identify underlying molecular causes of treatment resistance.

First, we benchmarked the ability of COSMOS+ to score ligand perturbations with the Cytosig dataset. While the Cytosig dataset comprises more than 1300 perturbation experiments, the new lightweight network pruning and scoring heuristics made it possible to apply COSMOS+ to each cytokine perturbation experiment individually. This allowed us to benchmark the overall performance of COSMOS+ with respect to ligand scoring, and find and fix errors in the prior knowledge that affect the MOON scoring procedure. We showed that certain ligands such as IFNA1 could be very specifically predicted to be active in samples treated with it. We also showed that some other ligands seemed to be harder to capture, either due to errors in our prior knowledge or the measurements themselves. Finally, we showed how some simple error fixing the prior knowledge, like in the case of OSM, could lead to dramatic improvement in our ability to predict specific perturbations. Other methods such as Nichenet (Sang-aram *et al*, 2025) exist that can also score the activity of ligands. However, those approaches are not directly comparable as they differ with respect to one key aspect. COSMOS+ aims to provide a general way of leveraging prior knowledge networks by assessing which parts of it best fit the observed measurements. Methods like Nichenet (Sang-aram *et al*, 2025), on the other hand, aim to provide a more focused and accurate estimation of the activity of specific molecular features, in this case ligands. They do this by leveraging not only prior knowledge networks but also other experimental measurements to generate weighted signatures for a given set of molecular features. This precludes a direct comparison of both approaches’ performances since, for example, Nichenet learns its weights from the Cytosig dataset itself among data sources. While such approaches are expected to offer better performances with respect to the specific task they are trained for, they offer less flexibility than a complementary method like COSMOS+ that can be used to score any molecular feature as long as they are found within a prior knowledge network.

Then, we illustrated how footprints and meta-footprints based on prior knowledge networks can be applied downstream of factor analysis. To do so, we analyzed the transcriptome, proteome and metabolome data of the NCI60 cancer cell lines. We estimated transcription factor activity and ligand-receptor scores from MOFA factor weights and we showed how COSMOS+ points out a potential biological crosstalk between JAK-STAT, ITGB1, PRKCA and citrate metabolism to be a specific regulatory mechanism associated with leukemia-like cancer cell lines. Thus, we showed how the resulting mechanistic hypothesis network of COSMOS+ is coherent with expected regulatory mechanisms, while hypothesizing novel control mechanisms spanning metabolites and signaling occurring in leukemic cell lines, such as the MAPK pathway being controlled by Acetaminophen or 4-beta-oestradiol. Taken together, these results highlight the synergy between MOFA, footprint and network methods to derive interpretable and actionable biological insights.

The application of footprint methods in the context of factor-based variance decomposition (i.e. partitioning the variance of a variable – or a set of variables – into components attributable to different sources or factors) is in direct continuity of the use of pathway enrichment analysis to biologically interpret the result of factor analysis methods. Factor analysis has been used for a long time in natural sciences (Thurstone, 1931), and they are commonly used to analyze omics data (Velten & Stegle, 2023). They are particularly suited to handle large cohorts of samples such as cancer patient cohorts (Yang & Michailidis, 2016; Rau *et al*, 2022; Freeman-Cook *et al*, 2021a; Argelaguet *et al*, 2018; Quintero *et al*, 2020) and cell lines collections (Knowles *et al*, 2019; Wang *et al*, 2021; Barretina *et al*, 2012; Garnett *et al*, 2012) without explicitly grouping samples. Most of the current efforts to interpret the results of such analyses have focused on characterizing known meta-data variables (e.g. clinical variables) that may correlate with factors, or using pathway ontologies to interpret the feature weights of factors (Argelaguet *et al*, 2019; Consiglio *et al*, 2020; Gonçalves *et al*, 2022; Schlechte *et al*, 2023; Li *et al*, 2022; Monaco *et al*, 2022; Hamsanathan *et al*, 2022; Park *et al*, 2022; Kwok *et al*, 2023; Gambacorta *et al*, 2022; Mangiante *et al*, 2023; Meng *et al*, 2019; Pekayvaz *et al*, 2023). However, there are many more types of domain knowledge that can potentially be used to interpret feature weights of factors beyond pathway ontologies, such as prior knowledge in the form of footprints and signed-directed networks can help to provide interpretable insights from factor weights. COSMOS+ is agnostic to the variance decomposition method used, as long as they provide a matrix of factor weights (e.g. PCA, MOFA, MCFA, MEFISTO and others) (Argelaguet *et al*, 2018; Brown *et al*, 2023; Velten *et al*, 2022).

Footprints and LR analysis can be performed without further downstream network analysis, and various network methods can be used without prior footprint analysis (Liu *et al*, 2019; Cancer Genome Atlas Research Network, 2013; Bradley & Barrett, 2017; Chowdhury *et al*, 2022; Massacci *et al*, 2023; Rosenberger *et al*, 2024; Garrido-Rodriguez *et al*, 2022). Nonetheless, network methods that specifically model flows of activity in biological networks particularly benefit from inputs generated from prior footprint analysis (Dugourd & Saez-Rodriguez, 2019). Footprint and LR analysis using factor weights opens the door for factor-specific mechanistic network-level hypotheses, which we demonstrate using the MOON algorithm on the output of a multi-omic factor analysis of the NCI60 dataset. MOON relies on iterative scoring, representing a “greedy search” alternative to global network optimization algorithms based on integer linear programming, such as CARNIVAL (Liu *et al*, 2019). The iterative upstream scoring procedure of MOON has similarities to the scoring approach of causalR (Bradley & Barrett, 2017), the main difference being that MOON scores each node iteratively, by using only direct downstream interactions at each step, while causalR sequentially considers all the nodes that can be reached downstream of a given node (or upstream of a set of deregulated RNA measurements) within increasing maximum number of steps and then aggregates the results obtained with different maximum number of steps. Furthermore, the MOON scoring procedure relies on a linear model while causalR is based on a form of over-representation estimation. Finally, MOON was developed natively in the context of multi-omic mechanistic hypotheses generation within COSMOS+, which makes it particularly suited to use with the multi-omic factors of MOFA.

One of the main benefits of prior knowledge network based methods is that these networks are interpretable, which makes them particularly suited to extract actionable insights from data (Lipton, 2018; Lou *et al*, 2012). It allows for an intuitive interpretation with natural language, therefore allowing users to spot mistakes easily as demonstrated with the OSM ligand in the Cytosig dataset. For example, an OSM-IL6ST interaction can be interpreted as: “OSM binds and activates IL6ST receptor”. Thus, bridging factor-based analysis (e.g. MOFA) and network integration methods (e.g. COSMOS) allows us to put the relationship captured by factor weights in the context of known biological mechanisms such as transcriptional regulations, post-translational modifications (PTM) mediated regulations or metabolite/metabolic enzyme interactions. Paradoxically, the sheer amount of prior knowledge available about biological molecules can negate the expected interpretability of such an approach (Merico *et al*, 2009). Therefore, there are still significant challenges to extract actionable insights from contextualized prior knowledge networks. To address these challenges, various approaches have been developed such as summarizing metrics (e.g. network flow with node centrality (Borgatti, 2005)) and matrix representation of graphs (Henry & Fekete, 2006; Bae & Watson, 2011). In this work, we sought to design an approach that would abstract the network into a matrix format that could also directly inform us about the biological functions regulated by each node of the contextualized network and score the significance of such regulations. We built upon the ideas of network flow (Ahlswede *et al*, 2000) and guilt by association (Hou *et al*, 2014) while taking the sign of the interactions into account, to intuitively and intelligibly highlight the most significant regulation events captured by a contextualized prior knowledge network. We refer to such an analysis as a pathway control analysis and demonstrate its use with the NCI60 dataset. We saw that it was able to recover expected regulation mechanisms, as some of the top gene-pathway interactions were found to be e.g. JAK regulates the JAK-STAT pathway. This also highlighted less intuitive interactions, such as the fact that a significant number of deregulated members of the MAPK pathway in NCI60’s leukemic cell lines are down-stream of molecular targets of Acetaminophen (Iorga *et al*, 2021; Shen *et al*, 2023).

While the NCI60 dataset is very useful to demonstrate the use of COSMOS+ in the context of signaling and metabolomic cross-talks is limited to only basal state measurements. Therefore, we also assessed the ability of mechanistic hypotheses generated by COSMOS+ to support the identification of drivers of treatment resistance by applying it on a novel combined transcriptomic and phospho-proteomic dataset of breast cancer cell lines with different resistance profiles exposed to CDK inhibitor drugs measured at early (2-4 hours) and late time points (72-96 hours). CDK4/6i is approved for the treatment of hormone receptor–positive, human epidermal growth factor receptor 2–negative (HER2–) advanced breast cancer in combination with endocrine therapies (ET). While CDK4/6 inhibition combined with anti-hormonal treatments delay disease progression, patients develop resistance and ultimately progress. Predictive biomarkers of clinical CDK4/6i resistance have been challenging to identify. Recent transcriptional profiling of the CDK4/6i Paloma 3 patient samples indicated that patients with high CCNE1 expression were less responsive to the CDK4/6i and fulvestrant combination, potentially implicating CDK2 activity in resistance, but no conclusive evidence was found at the level of CDK2 expression alone (Turner *et al*, 2019). Combination treatment strategies that can simultaneously inhibit multiple compensatory signaling pathways mediating CDK resistance provide an appealing clinical treatment strategy. In this context, we sought to use COSMOS+ to examine both known and potentially novel mechanistic hypotheses mediating response and resistance to CDK4/6i. COSMOS+ recapitulated known signaling and transcriptional regulation components following CDK inhibition such as the CDK2,4,6, RB1 and E2Fs crosstalk, while also highlighting a wide set of other potentially important mechanisms that were differentially regulated between the sensitive and resistant cell line, such as a YWHAQ/E2F1/FOXO3 and a CDK2/NOTCH crosstalk. COSMOS+ was also able to highlight hypothetical mechanisms specifically different between the CDK2/4/6i and CDK4/6i treatments, such as the activation of CASP6 and HIPK2 downstream of CDK2 inhibition.

We then compared the mechanistic hypothesis identified by COSMOS+ after drug perturbation with essentiality and drug sensitizing potential of gene knock-outs (KO) data from genome wide crispR KO experiments. The poor recall suggests that there is a large gap between the ability to identify deregulated signaling and gene regulation between different cell lines treated with a drug and the actual identification of direct drivers of drug resistance. Of note, the crispR KO experiment itself has limitations because COSMOS+ can identify essential genes that will likely not be detected in a crispR KO sensitization assay. Indeed, it has been shown that general gene essentiality can confound sensitization assays because they cause cell death regardless of the context (Bock *et al*, 2022), which was further confirmed in our analysis. We show that when accounting for generally essential genes in the crispR KO screen comparison, the correlation between COSMOS+ hits and crispR hits increases, but also loses context specificity (no more differences between resistant and sensitive cells). Nonetheless we also were able to highlight biological pathways such as nucleotide excision repair that seemed to be particularly enriched with genes that were consistent between activity deregulation and being sensitization targets. We also showed how pathway control analysis could correctly recapitulate the control of E2Fs transcription factors over cell cycle processes in a breast cancer cell line dataset and how it is differentially regulated between sensitive and resistant cell lines treated with CDK inhibitors.

Finally, we demonstrated how COSMOS+ can be used with clinical baseline patient cohort data to complement biomarker predictions based on transcriptomics data alone. We showed how a Cox survival analysis performed with MOON scores instead of transcriptomic data allowed for the identification of expected markers that were missed by transcriptomic data alone. We then showed that the Cox coefficients based on MOON scores were correlated with MOON scores estimated from a cell line model of resistant and sensitive cell lines treated with similar drug regimens, suggesting that mechanisms identified in such a cell line model could support the identification of clinical biomarkers. While small (Pearson correlation coefficients = -0.12), the correlation was highly significant (p-value < 10-6), which is consistent with the idea that these cell line models can only capture a fraction of all the potential causes of treatment resistance in a patient cohort. We also demonstrated the complementarity of transcriptomic and MOON features to predict patient outcomes, further supporting the idea that mechanistic prior knowledge can increase the amount of information that can be extracted from omic data. For example, CDK2 and RB1 expression could not conclusively be related to worst prognosis at the level of their expression, but their MOON score could.

While the approach described here can be useful to help researchers to extract actionable insights from their data, it has several limitations. The first and foremost limitation comes from the use of prior knowledge to support those analyses. Indeed, prior knowledge can never be complete and is rarely error proof. This leads to two types of errors that methods that rely on prior knowledge can make. First, it will potentially miss interactions that underlie the observed data. COSMOS+ is fully constrained within the space of the prior knowledge that is used, and therefore will miss any interaction that isn’t covered in the prior knowledge, which is known to be incomplete and biased (Garrido-Rodriguez *et al*, 2022). For example, this is why CCNE1 MOON score was not estimated for the Paloma3 patients, even though its expression was measured. The second type of error comes from using interactions that are the product of miss-annotated database entries or flawed experimental sources to interpret the data at hand, such as the OSM ligand in the Cytosig benchmark. This can lead to methods highlighting false interactions as the potential cause of variations observed in the data. Due to the recursive nature of the MOON algorithm, errors in the PKN have a risk of being propagated upward and affect many upstream scores. As we show with the analysis of the Cytosig dataset, this error can be mitigated by cross-checking the source of any critical interactions highlighted by the analysis (consequently, making it easy to perform such cross-checking is crucial) and, if it becomes apparent that the interaction is fictitious, reporting it to the source database, removing it and repeating the analysis without its influence on the pipeline. As the coverage and quality of prior knowledge resources keep improving, COSMOS+ results will become more accurate.

The ability to handle and recover negative feedback loops in prior knowledge networks could be an interesting future direction to explore with such modeling strategies. Indeed, negative feedback loops cannot be recovered by this type of methods unless timepoints are explicitly considered by the network scoring procedure. Recovering a negative feedback loop requires the observation of a given node switching sign between two timepoints. Since COSMOS+ can be applied downstream of factor analysis, it can benefit from the fact that such type of analysis can be applied to time series and spatially resolved data. Beside finding mechanistic networks connecting data that already encode time, the network scoring procedure could be adapted to explicitly take time into account, e.g. using each previous iteration as a starting point for the next one, similar to the PHONEMES-ILP method (Gjerga *et al*, 2021). Of note, MOON recovers positive feedback loops, because it only prunes out edges from the network when there is an explicit incoherence between the activity sign of multiple interactors. However, an additional complexity of time-course data that is already problematic with single omic dataset but becomes particularly prevalent when dealing with multi-omic time courses is how different time points should be matched across multiple omics. For example, in the case of the phospho-proteomic and transcriptomic breast cancer cell line dataset that is introduced in this study, we showed that there was no clear indication that the 24h time point transcriptome should be matched with either the 2h or 24h phospho-proteomic when performing an integrative analysis.

Finally, once a hypothesis is formulated in the form of a mechanistic network representing a deregulated signaling pathway or gene regulation program, this hypothesis can be validated and translated into actionable insights. A common example of such actionable insight in pharmacological research is the identification of new drug targets, especially for cells and tissues that are resistant to existing treatments. Here, the mechanistic hypotheses that were generated by COSMOS+ for the breast cancer cell-line multi-omic dataset encompassed both expected mechanisms (such as the CDK2/4/6-RB1-E2Fs axis) as well as less known/studied mechanisms in this context, especially presented altogether in a mechanistic network such as crosstalk between YWHAQ, E2F1, PTTG1 and TP53. However, it requires care and crosschecking to bridge the mere identification of biological mechanisms that are different between cell lines and/or affected by the exposure to a given drug and the actual identification of targetable drivers of resistance (Khan *et al*, 2024; Lei *et al*, 2023). In this study, we show that there is partial consistency between differentially regulated molecular drug responses and molecular drivers of treatment sensitization. Furthermore, the good precision of COSMOS+ to capture sensitization targets but poor recall is consistent with the idea that such a model, while being able to accurately capture important biological mechanisms, is not yet able to generate predictive mechanistic models of cell signaling.

Nonetheless, the mechanisms hypothesized by COSMOS+ seem to complement the information in the omic dataset alone. Indeed, when COSMOS+ was applied at the level of the transcriptome measurements of the breast cancer cohort Paloma3 patients, it revealed markers of treatment outcomes that could not have been captured at the level of their expression alone. For example, RB1 activity estimated from its expected downstream regulated expression target was significantly associated with the worst outcome in the treatment arm of the Paloma3 cohort, while its own expression was marginally associated with the outcome in the wrong direction (that is, high expression of the RB1 tumor suppressor associated with worst outcome). This could be explained by the fact that RB1 expression is regulated by a negative feedback loop under the control of E2F transcription factor. While RB1’s normally leads to an inhibition of E2Fs activity, if E2F escapes such control, for example through an overwhelming activation of other upstream regulators, this may lead to an increased expression of RB1, even though its effective activity is lower than in context where E2Fs are lowly expressed. This complementarity is further illustrated by the increased correlation between a PFS prediction performed with RNA or MOON scores alone compared to a hybrid signature. Therefore, it will be interesting in the future to assess if COSMOS+ could be used in combination with omic features to improve the performance of predictive approaches.

COSMOS+ could in principle be applied to any type of omic data that can be related to activation or inhibition of node activities in a biological molecular network, such as chromatin accessibility (Grandi *et al*, 2022), mutational data (Wilkerson *et al*, 2014), high throughput protein conformation (Cappelletti *et al*, 2021; Mateus *et al*, 2020), microRNA (Sun & Zhang, 2022) or other PTMs beside phosphorylation (Sun & Zhang, 2022; Varland *et al*, 2023; Potel *et al*, 2025). For this, prior knowledge resources that would allow linking such measurements to functional readout are needed (e.g. non-ambiguous miRNA-target interactions, functional annotation of ubiquitination and acetylation, etc…). It is also important to keep in mind that the mechanisms that are proposed by methods like COSMOS+ are hypotheses and benchmarking them is critical and non-trivial. Toward this end, COSMOS+ new methods were also implemented within the Networkcommons (Paton *et al*, 2024) benchmarking framework and future work will focus on evaluating their accuracy with respect to various other scenarios beside the Cytosig benchmark presented in this study.

To conclude, we present an analytical workflow, implemented in the tool COSMOS+, that combines factor analysis, mechanistic features, and prior knowledge networks to extract meaningful biological insights from multi-omic data. The software and a step-by-step tutorial are available at https://github.com/saezlab/cosmosR.

## 4. Methods

### 4.1 NCI60 Multi-omic data processing

#### 4.1.1 Transcriptomic data

The NCI60 transcriptomic dataset was obtained through the https://discover.nci.nih.gov/cellminer/home.do portal. The data was obtained as log2(FPKM + 1) values. To filter out lowly expressed genes, we set any log2(FPKM+1) value that was lower than 1 (which corresponds to an original FPKM value of 1) as NAs. Then, any gene that had an NA value in more than ⅔ of samples was excluded. Next, genes were ordered by their standard deviation across samples, and the top 6000 genes were kept for further analysis with MOFA. To decide on the number of transcripts to keep, we tried to reduce it as much as possible while still having enough data to estimate as many transcription factor (TF) activities. We used CollecTRI regulons to determine which transcript was a target of a TF, then we checked how many TF would still have at least 10 targets when we progressively reduce the number of transcripts from 8354 to 0 (Figure 1D), from most variable to least variable across cell lines. We saw that at 6000 transcripts, >100 TFs still had at least 10 target transcripts. At 3000 transcripts, that number would go down to > 55 Tfs. Considering this, we settled for 6000 transcripts, so that we would still be able to estimate a good number of TFs for downstream analyses., at 6000 transcripts, the average number of targets per TF was > 90 (Figure 1D), ensuring relatively robust TF activity estimates.

#### 4.1.2 Proteomic data

The NCI60 proteomic dataset was obtained through the https://discover.nci.nih.gov/cellminer/datasets.do portal. The data was obtained as log10(intensity + 1) from Sequential Window Acquisition of all Theorical MS intensity values (SWATH). Duplicated protein entries were averaged to generate a set of unique proteins with corresponding intensity values. Any log10(intensity+1) value of 0 was converted to NA (which corresponds to an original intensity value of 0). Next, proteins were ordered by their standard deviation across samples, and the top 60% of proteins were kept for further analysis with MOFA.

#### 4.1.3 Metabolomic data

The NCI60 metabolomic dataset was obtained through the https://wiki.nci.nih.gov/display/NCIDTPdata/Molecular+Target+Data porta. The data was obtained as log2(MS intensity + 1). Triplicates were averaged to match the structure of the RNA and proteomic data. Values above 32 were considered outliers and were replaced by NAs. Metabolite identifiers were matched to primary HMDB identifiers manually, one by one.

### 4.2 MOFA

The different omics were assembled into a single mofa ready input file. For each type of omic data, cross-correlation between features were evaluated using Pearson correlation with pairwise complete observations. The correlation between each transcript and its corresponding protein was also evaluated using Pearson correlation with pairwise complete observations. Mofa was run with the following options: scale_groups = “False”, scale_views = “False”, likelihoods = c(’gaussian’,’gaussian’,,’gaussian’), spikeslab_weights = “True”, spikeslab_factors = “False”, ard_factors = “True”, ard_weights = “True”, iter = “1000”, convergence_mode = “fast”, startELBO = “1”, freqELBO = “1”, dropR2 = “0.001”, gpu_mode = “True”, verbose = “False” and seed = “1”. The maximum number of factors was set sequentially between 5 to 15, 20 and 58. The resulting models were compared by correlating their weight matrices (Pearson correlation). Since factors seemed to converge after setting the number of factors to 10 and above, the model with 10 maximum factors parameter was kept for subsequent analysis.

Clinical information about cell lines were discretized when necessary (such as age variable split between > 50 and <= 50 years old). The clinical variables were then converted into a sample-clinical variable edge table. The ULM method of decoupleR was used to model each factor Z values as a function of their belonging or not to each given clinical variable categories. The resulting top enriched clinical variable categories in each factor were thresholded using an absolute t-value of 2 (t-value of ulm’s linear coefficient).

### 4.3 Footprint and ligand-receptor analysis with MOFA weights

Ligand-receptor interactions were obtained from the consensus resource of the LIANA R package (v 0.1.5), and decomplexified using the decomplexify function. Then, TF target interactions were converted into a ligand-receptor set collection by associating each ligand and receptors to their corresponding ligand-receptor (LR) set. TF-target regulons were obtained using the get_collectri function of the decoupleR R package (v 2.5.2). For each factor, decoupleR’s ULM function was used for each TF and LR set to model the RNA MOFA weights as a function of their belonging or not to each TF and LR set, respectively.

For each factor, weights of transcripts and their corresponding protein were correlated using Pearson coefficient.

### 4.4 COSMOS

The weights of factor 4 were extracted to prepare them as input for the downstream mechanistic hypothesis generation. The prior knowledge network (PKN) was obtained from the COSMOS package (v 1.5.2). to select genes that were consistently regulated at both the level of RNA and protein in each factor, the weight of each gene that had an absolute value of less than 0.2 for their RNA weight and less than 0.05 for their protein weight were set to 0. Metabolite weights that had absolute weight values < 0.2 were also set to 0. The thresholds were estimated from visual inspection of the weight distributions in factor 4.

#### 4.4.1 COSMOS with Meta-fOOtprint aNalysis (MOON)

MOON is a way to score the nodes of a prior knowledge network using successive iterations of the Univariate Linear Model (ULM) method from decoupleR’s package over a network, layer by layer, starting from a set of downstream nodes. At the first iteration of the MOON algorithm, ULM is run using measurements (e.g. metabolomic abundances) or activity scores (e.g. TF activity scores) as input data, and the prior knowledge network as a regulatory network input. Since ULM only considers regulons that have measured downstream targets, only nodes that are directly upstream of the provided measurements of activity scores will be scored at this stage. Thus, the score assigned to nodes at the first iteration represents how their direct targets are significantly up or down-regulated (consistently with the sign of the interactions), compared to the rest of the downstream layer data points. Once those nodes have received a score, they become the input downstream layer for the next scoring iteration, and are removed from the list of possible source nodes of the network to avoid multiple scoring of each node. This process is repeated as long as (1) the iteration number is lower than a user defined limit, (2) there are still source nodes upstream of the last layer that was scored in the network. The output of the MOON function is a dataframe with all the nodes of the input network scored, as well as a corresponding layer index, indicating at which iterations they were scored (that is, how many steps from the original downstream layer input they are).

If an upstream layer input is provided, any node that has a score that is different from the data provided in the upstream layer input is removed from the output dataframe. The algorithm describing the base MOON function is as follow:

##### Pseudocode for the moon Algorithm

###### Algorithm 1

***moon***

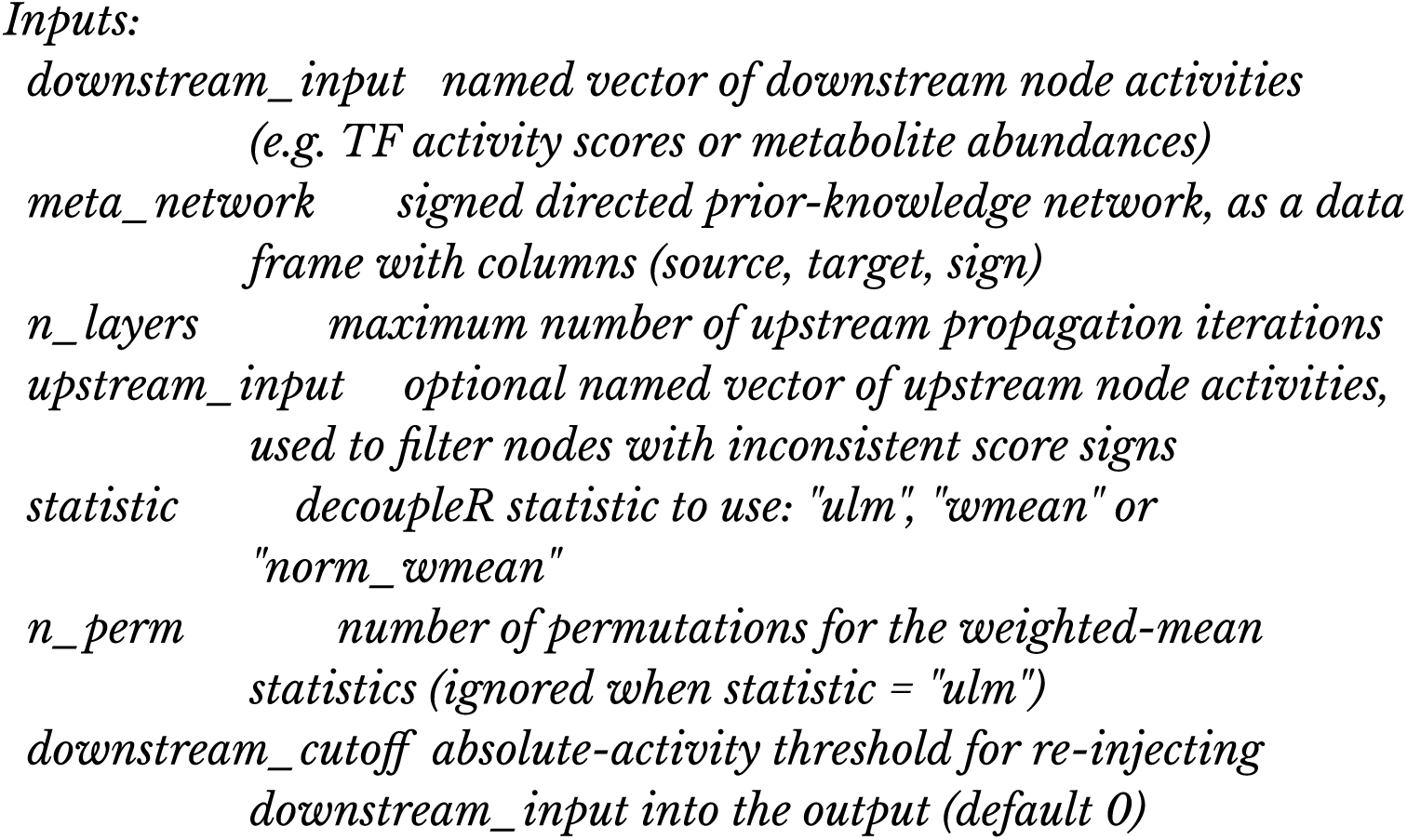
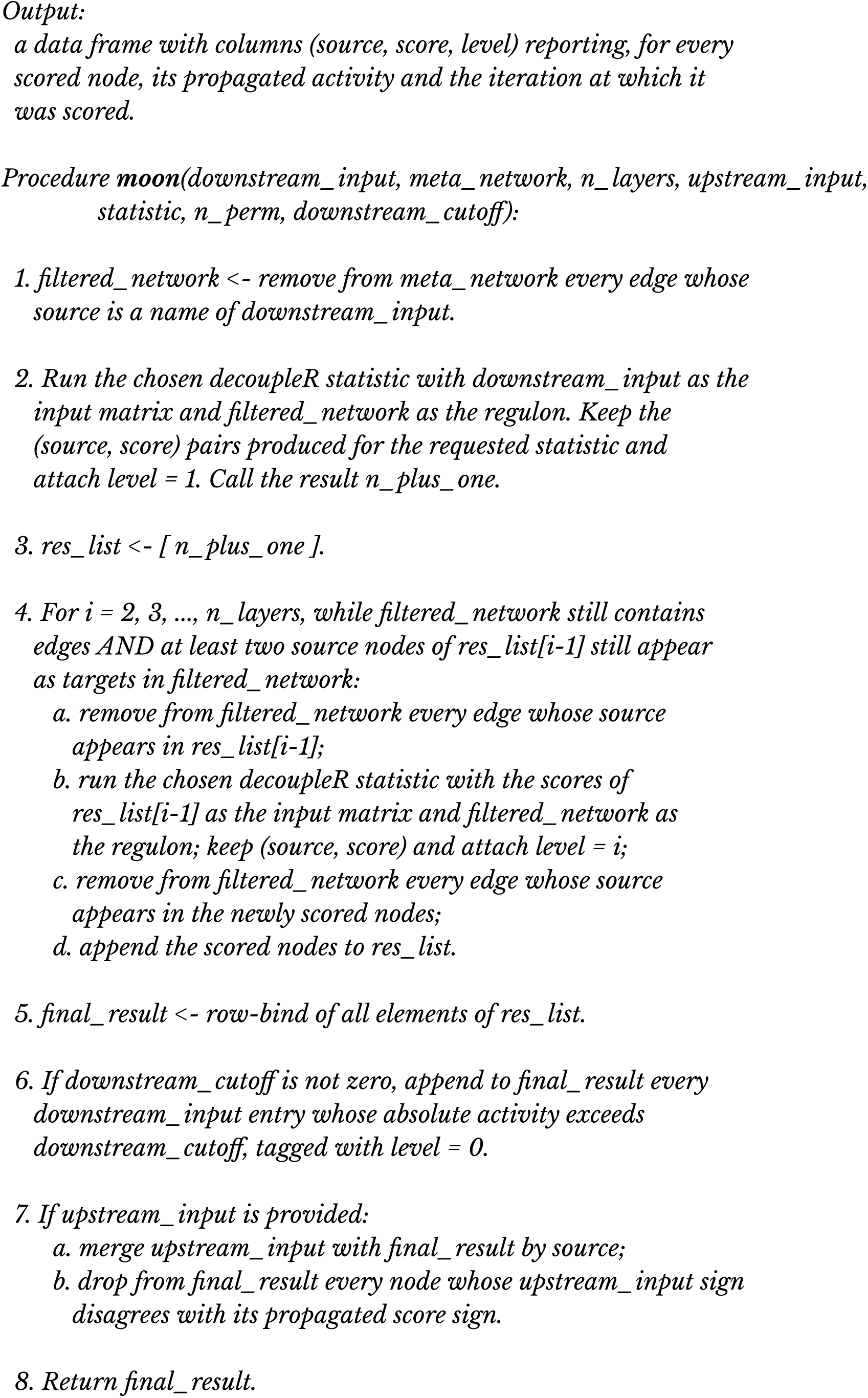

If RNA data points (here, mofa weights) are provided, a check can be performed to remove any interaction from the network that connects a TF and a downstream gene that has an incoherent expression sign with the TF MOON score. Thus, the moon function and the TF-target coherence check can be run in a loop until the output of the moon doesn’t contain any incoherence between TF scores and downstream targets. The algorithm to remove incoherent TF-target interactions is as follow:

##### Pseudocode for filtering incoherent TF target interactions

###### Algorithm 2

filter_incoherent_TF_target

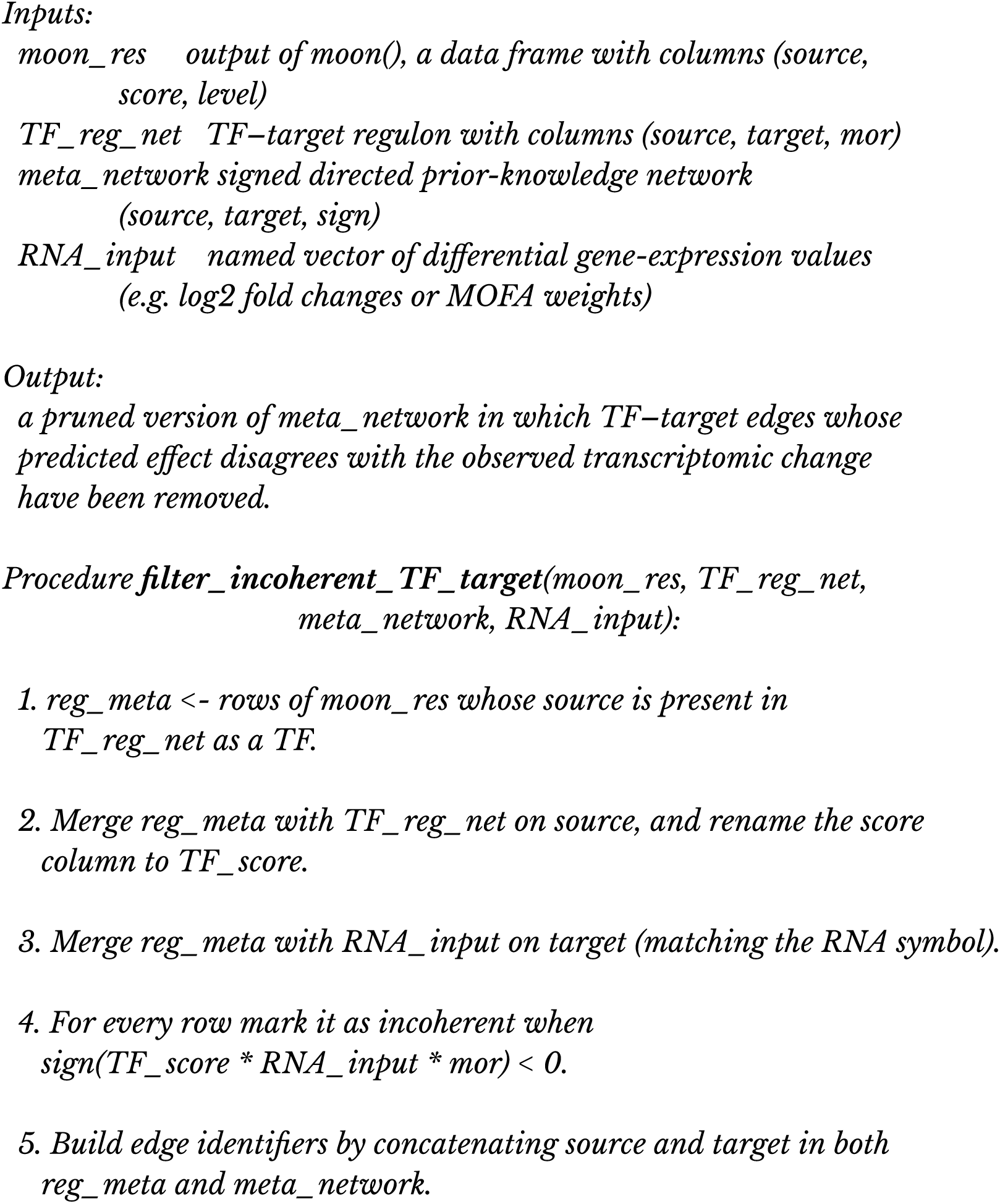
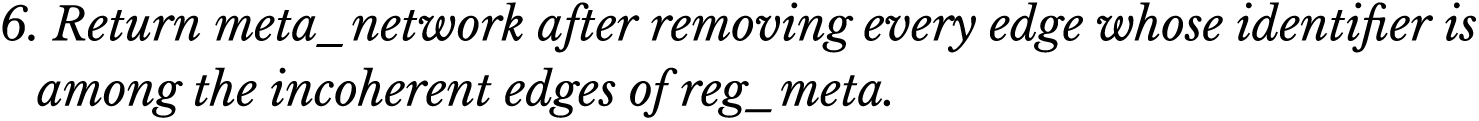

If there are multiple redundant parallel paths in the network, this can lead to score biases, because two nodes that have the same downstream targets will receive the same score, and thus will count double in the background of the next scoring iteration. To avoid this issue, the PKN can be compressed using a simple procedure that combines any set of nodes that have exactly the same direct downstream direct targets into single virtual nodes that can later be decompressed back into the original nodes using a hash table mapping virtual nodes with the original nodes of the network they are replacing (<u>Supplementary</u> <u>Figure S1B</u>). The algorithm to compress nodes with same children is as follow:

##### Pseudocode for compressing the network based on nodes sharing the same children

###### Algorithm 3

*compress_same_children*

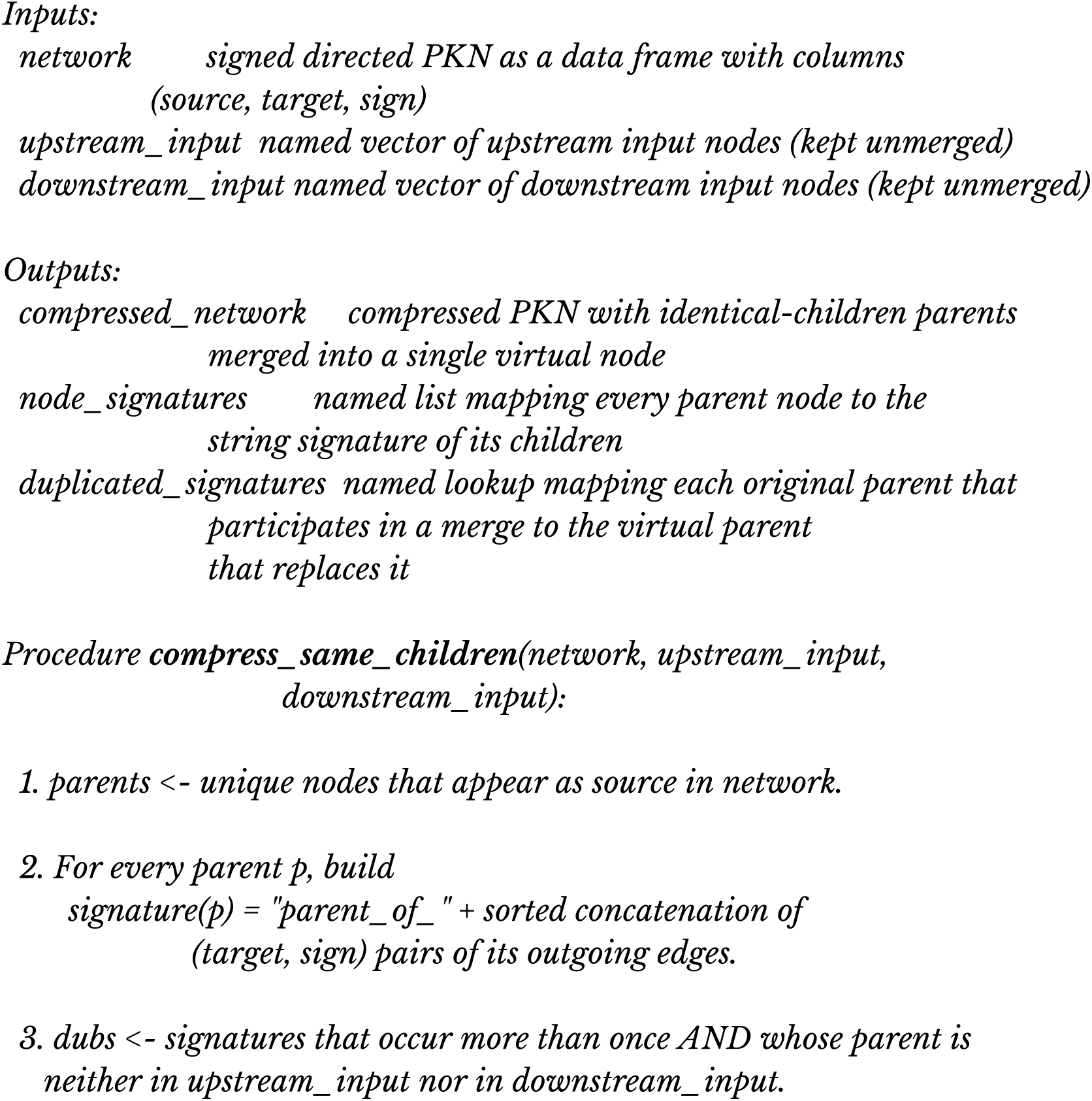
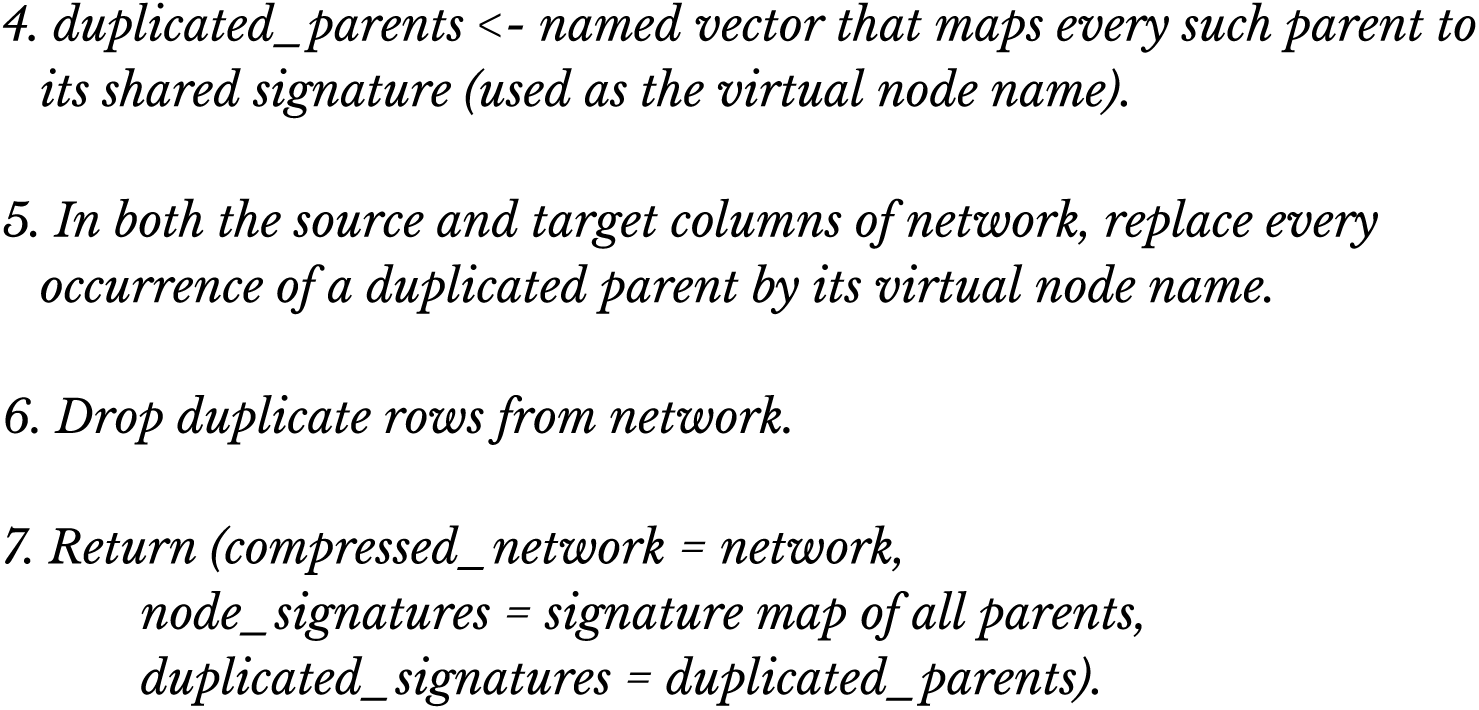

The algorithm to decompress the network is as follow, were SIF is a Simple Interaction Format network and ATT is a node Attribute data-frame:

##### Pseudocode for decompressing the solution network

###### Algorithm 4

decompress_solution_network Inputs:

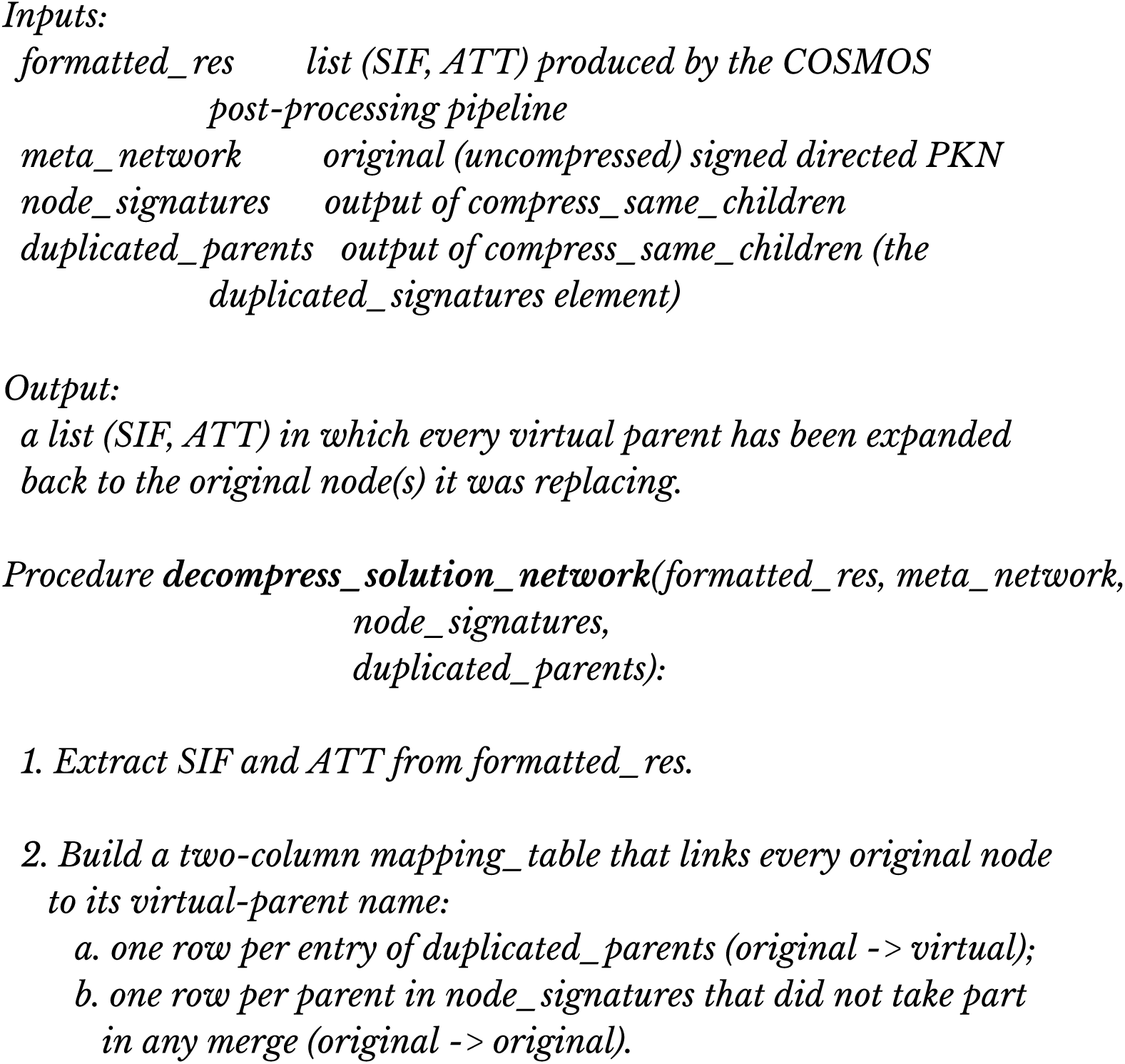
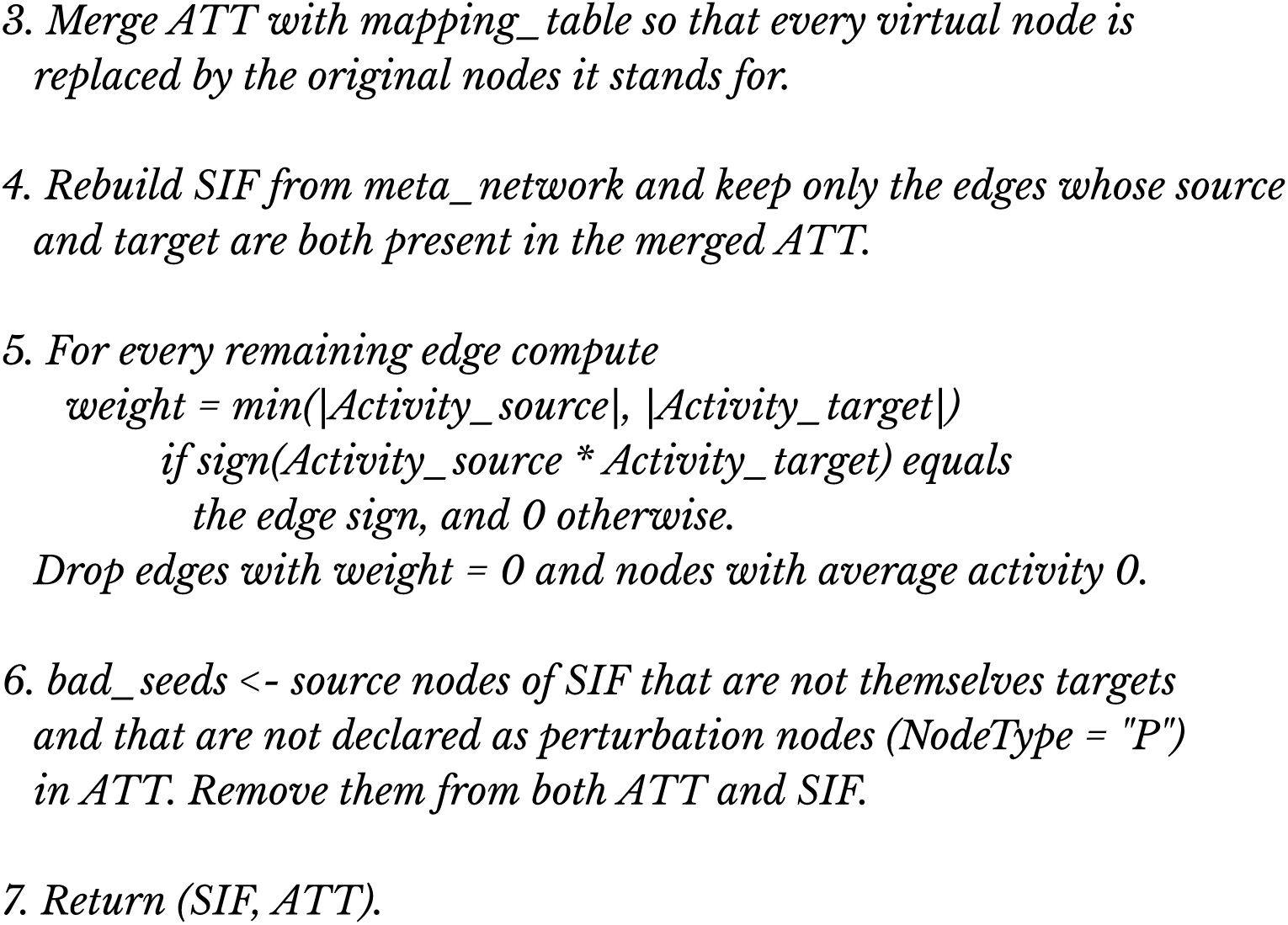

Finally, to generate a solution network with similar properties as the output network of the classic CARNIVAL pipeline, the node MOON scores can be mapped on the PKN, and each interaction in the network that is not consistent with the sign of the MOON score of it’s connecting node is removed from the PKN. It is also possible to define a score threshold to only keep paths connecting upstream and downstream input layer nodes that go through nodes with absolute scores that are strictly higher than the threshold. The algorithms are create a reduced solution network are as follows:

##### Pseudocode for the reduce solution network function

###### Algorithm 5a

*reduce_solution_network*

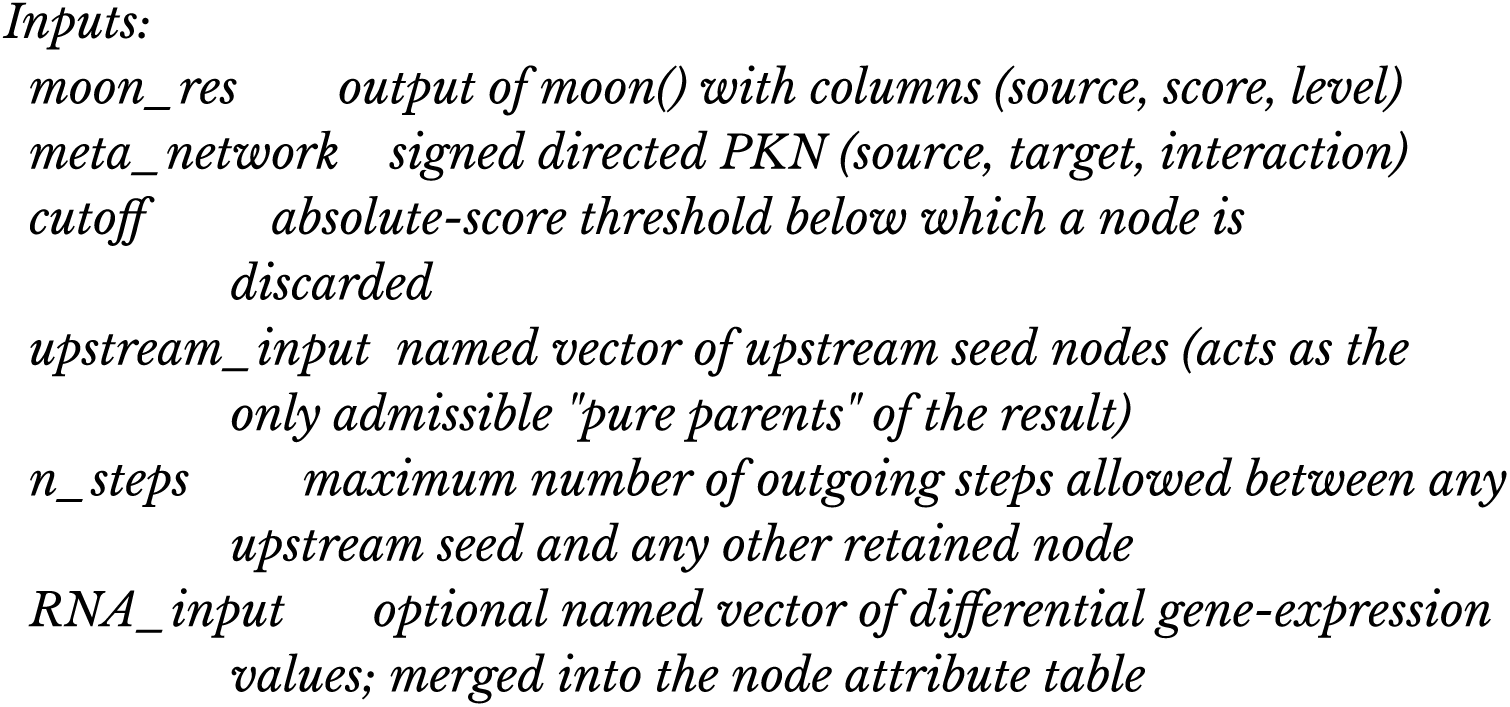
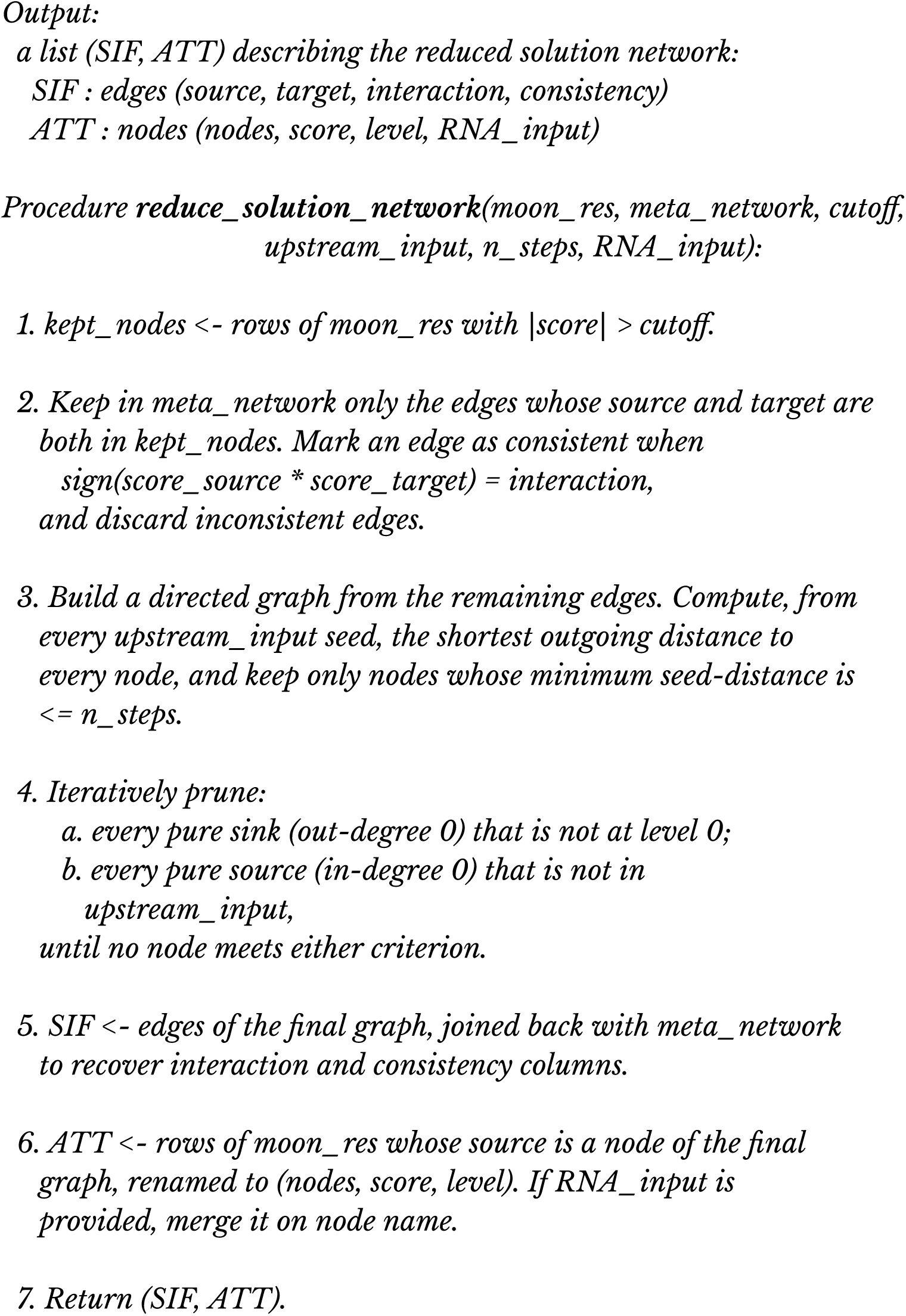

###### Algorithm 5b

*reduce_solution_network_double_thresh*

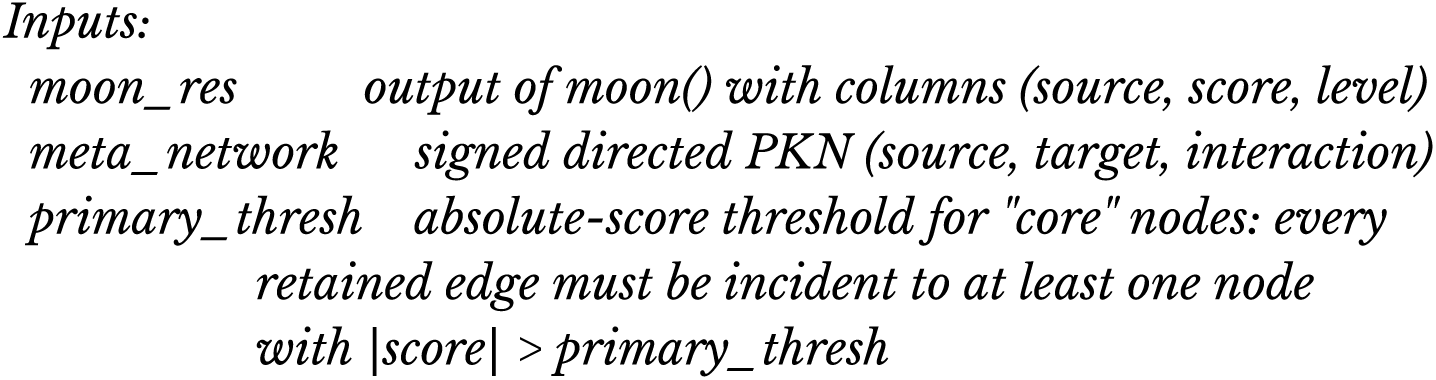
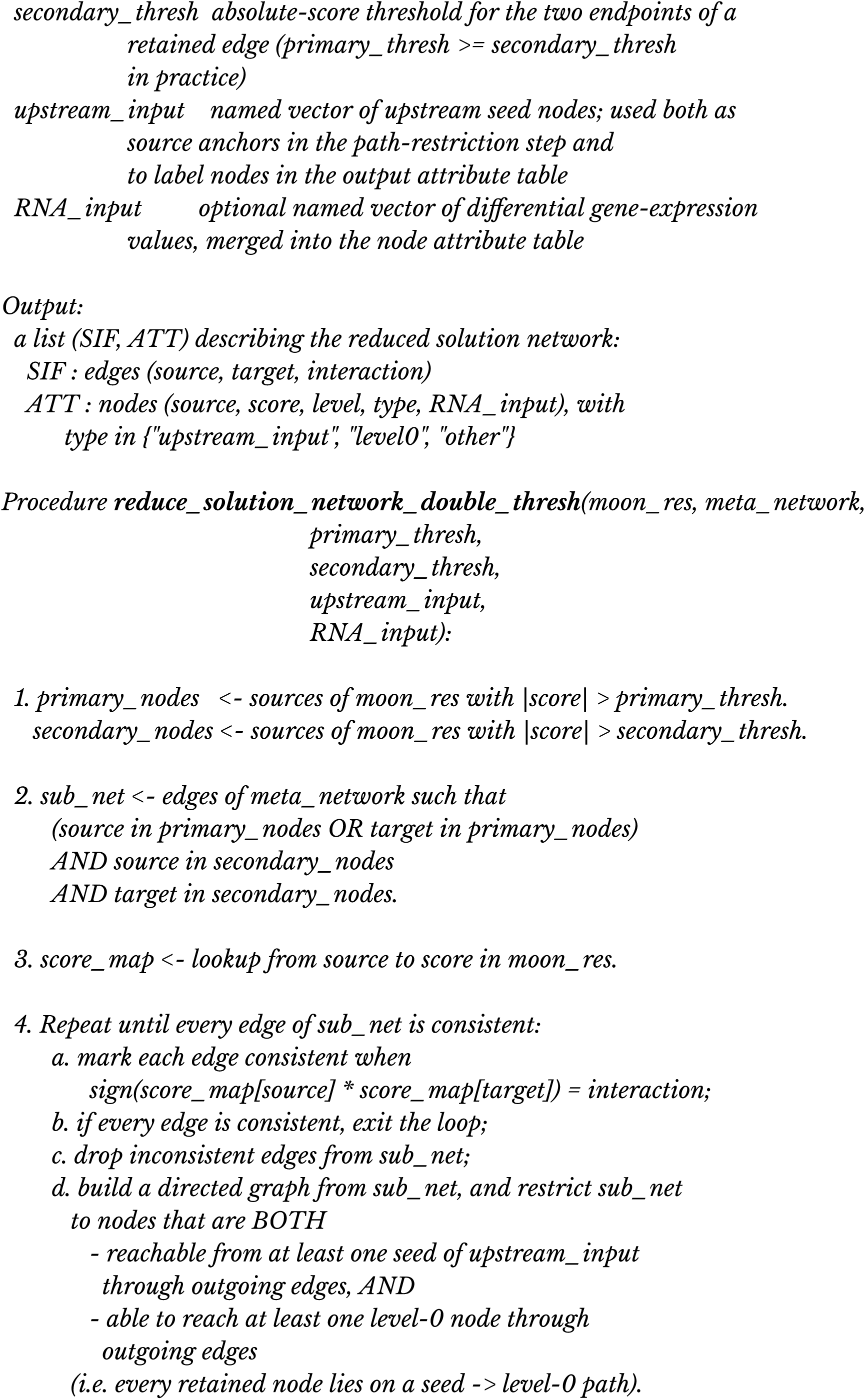
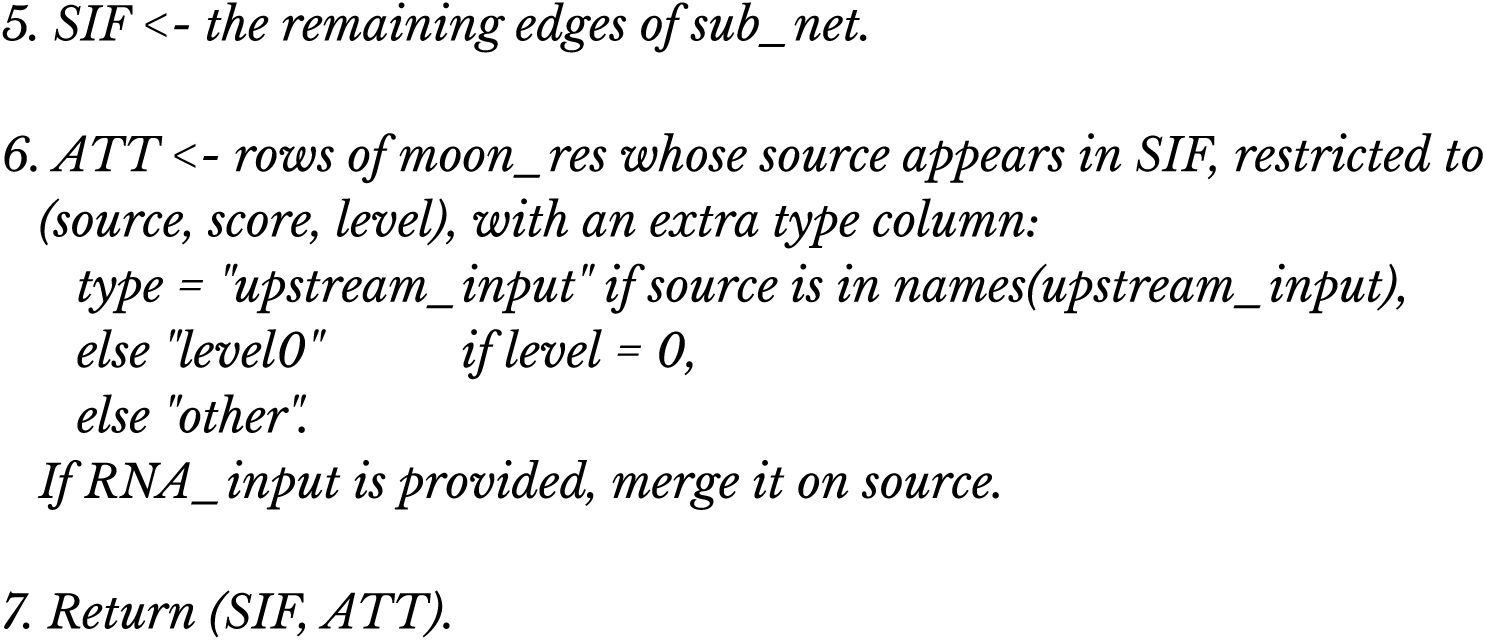

For the MOON run, the prior-knowledge-network was processed slightly differently. Since the MOON scores are estimated compared to a background distribution, we did not exclude non-regulated TF, genes and metabolites from the network or from the inputs. The PKN was only filtered by removing any gene that wasn’t part of the filtered Log2(FPKM+1) dataset, and we kept controllable and observable nodes that were within 6 steps of upstream and downstream inputs in the network. The network was then compressed using the procedure mentioned above.

We ran MOON first between receptors and downstream TF and metabolites, like the first CARNIVAL run. The resulting scores were mapped on the input network and an absolute score threshold of 1.5 (which represents a ULM t-value) was used to generate a reduced solution network comparable to the CARNIVAL solution network.

We then ran MOON a second time, between upstream TF and downstream ligands, using a limit on the number of steps of 1. This essentially allows us to generate a network that only contains direct interactions between TF and target ligands where the TF activity score is coherent with the sign of the downstream ligand. An absolute score cutoff of 1.5 was also applied to remove non-significant TF-ligand interactions (at least according to this threshold).

The two resulting networks of mechanistic hypotheses resulting from the first and second MOON run were combined by taking the union of their edge and attribute lists into a single network connecting ligand, receptors, TF and metabolites altogether.

### 4.5 MOON cytokine scoring with cytosig data

#### 4.5.1 Cytosig data collection

The sample-level preprocessed data was downloaded from the FDC platform (https://curate.ccr.cancer.gov/, accessed 22nd April 2022), using the code provided (https://github.com/data2intelligence/FDC_treatment_profile). For each treatment, we calculated a Z-score per gene as follows: the difference of the mean expression between treated and control samples was divided by a smoothed standard deviation of the gene expression of the control samples. The standard deviation was modeled using a locally-weighted regression (LOESS) based on the mean gene expression, using the scikit-misc python package. We kept only perturbations that had at least two matching control samples (as required for computing standard deviations).

#### 4.5.2 Cytosig filtering

We filtered out genes that had no expression measured (NA values) in > 1000 cytosig comparisons. We renamed the cytokines of cytosig to their corresponding gene symbols, or HMDB identifiers for metabolites. Then, we filtered out cytokines that were not present in the COSMOS meta-prior knowledge network.

#### 4.5.3 Cytokine scoring

For each cytosig gene expression signature (composed of cytosig z-scores), we filtered out any unmeasured genes (NA values), then we used decoupleR’s ULM method with the CollecTRI TF-target network to estimate the T-values of linear models for each TFs fitting gene expression z-scores as a function of their mode of regulation (-1, 0 or 1), referred to as the TF score. Then, for each resulting TF score profile, we filtered out specifically the COSMOS prior knowledge network for genes that were not measured in the corresponding gene expression profile. We then filtered out any node of the COSMOS prior knowledge network that couldn’t be reached within ten steps upstream of the TFs. Then the COSMOS prior knowledge network was compressed as explained in <u>(4.4.1)</u>. Moon scores were then computed using decoupleR’s ULM method as explained in <u>(4.4.1)</u>.

#### 4.5.3 MOON scoring networks

To visualize and interpret each MOON score, we implemented a function to recover the sub-network of the prior knowledge network that contains the direct and indirect downstream targets of a given node that were used for its MOON score computation. The algorithm of the function is detailed as follow:

##### Pseudocode for the Get Moon Scoring Network Function

###### Algorithm 6

*get_moon_scoring_network*

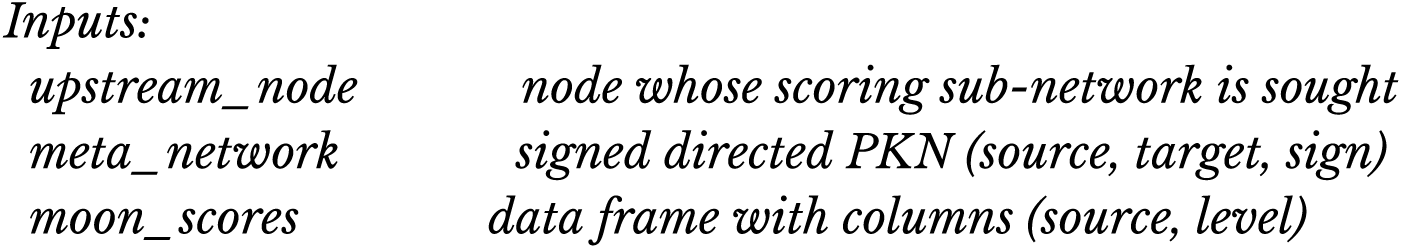
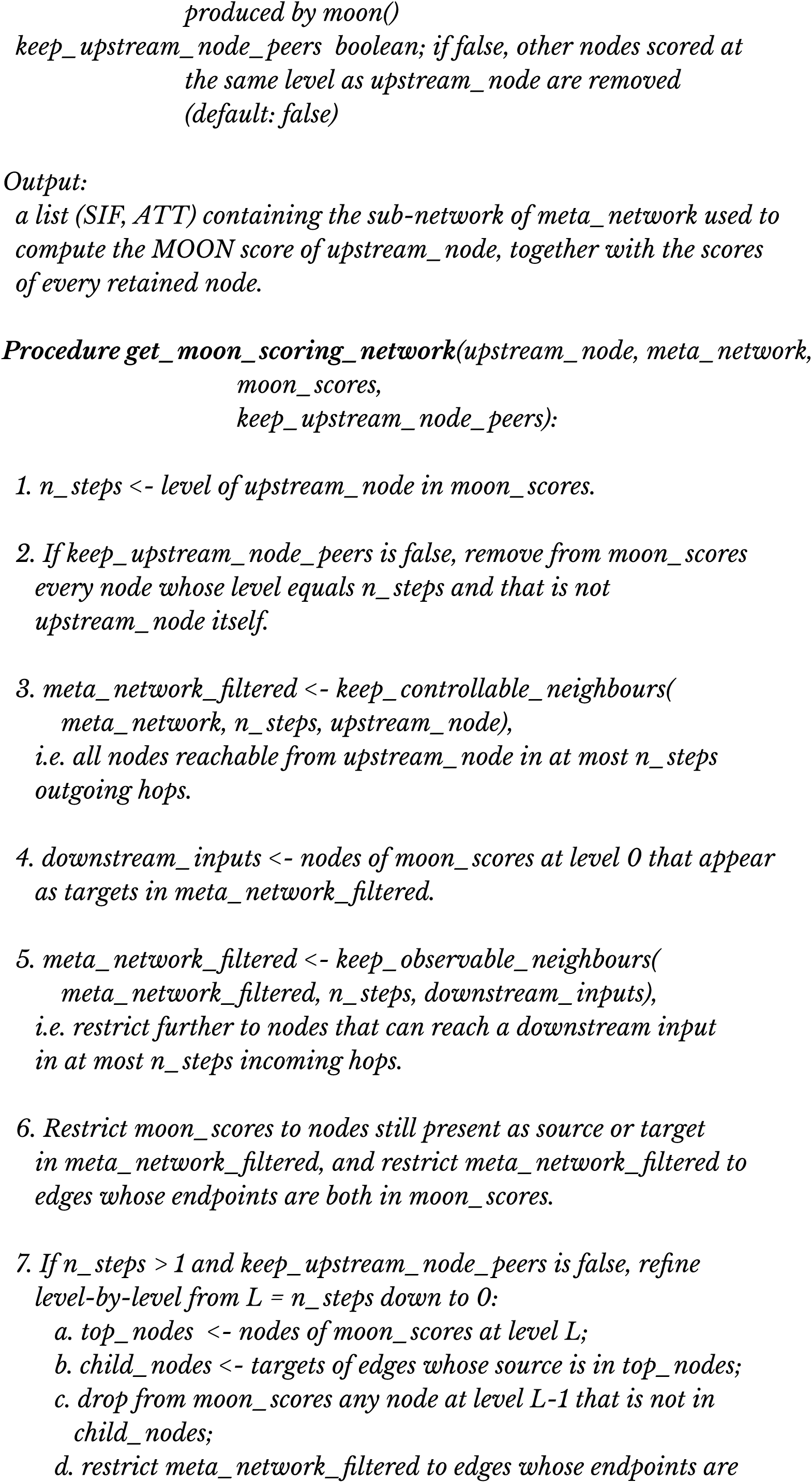
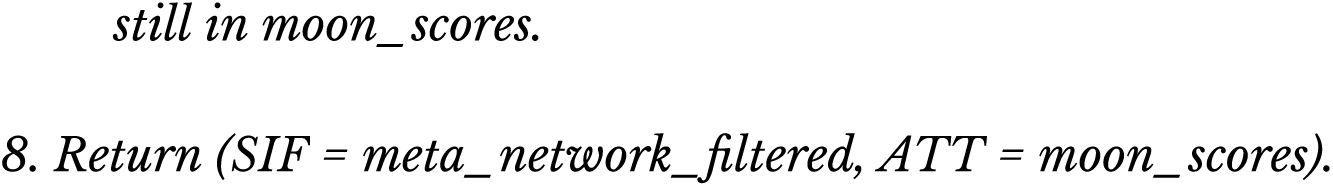

### 4.6 Pathway control analysis

To elucidate the potential pathways modulated by key regulatory nodes in our molecular network, we developed an algorithm termed “Find_Controlled_Pathways.” This algorithm commences by identifying a set of “top nodes” from the combined results of the moon propagation algorithm (full_moon_res_combined). These top nodes are distinguished based on an absolute score threshold greater than 1.5 (this value can be set differently at the user’s discretion). For each top node, the algorithm employs the COSMOS:::keep_controllable_neighbours function to ascertain downstream nodes within a maximum number interaction steps in the solution network (two steps in this study). If this filtered set of downstream nodes is not empty, an Over-Representation Analysis (ORA) is executed using the piano::runGSAhyper function, utilizing the NABA and KEGG pathway databases in this study. Other pathway set collections can be provided at the user’s discretion. The resulting p-values, along with a computed log2 fold ratio, are associated with each node of interest and pathway, and these are stored in a data frame.

The algorithm aggregates the data across all nodes of interest into a single data frame (pathway_control_df), which is then reshaped to feature each pathway as a row and each node of interest as a column, with the corresponding p-values populating the cells. This comprehensive table serves as a pathway control matrix, enabling the identification of pathways significantly influenced by specific nodes within the regulatory network, and vice-versa. The algorithm for the pathway control analysis is as follow:

#### Pseudocode to find controlled pathways

##### Algorithm 7

find_controlled_pathways Inputs:

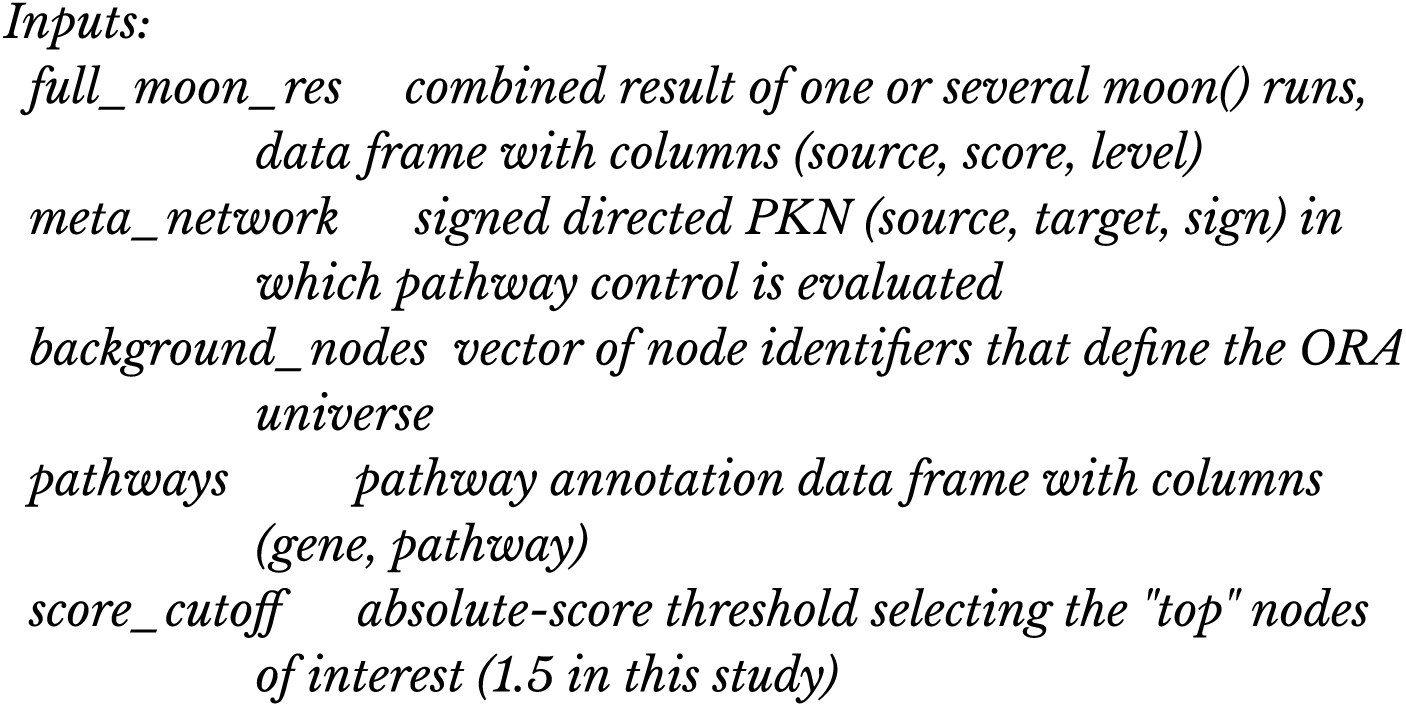
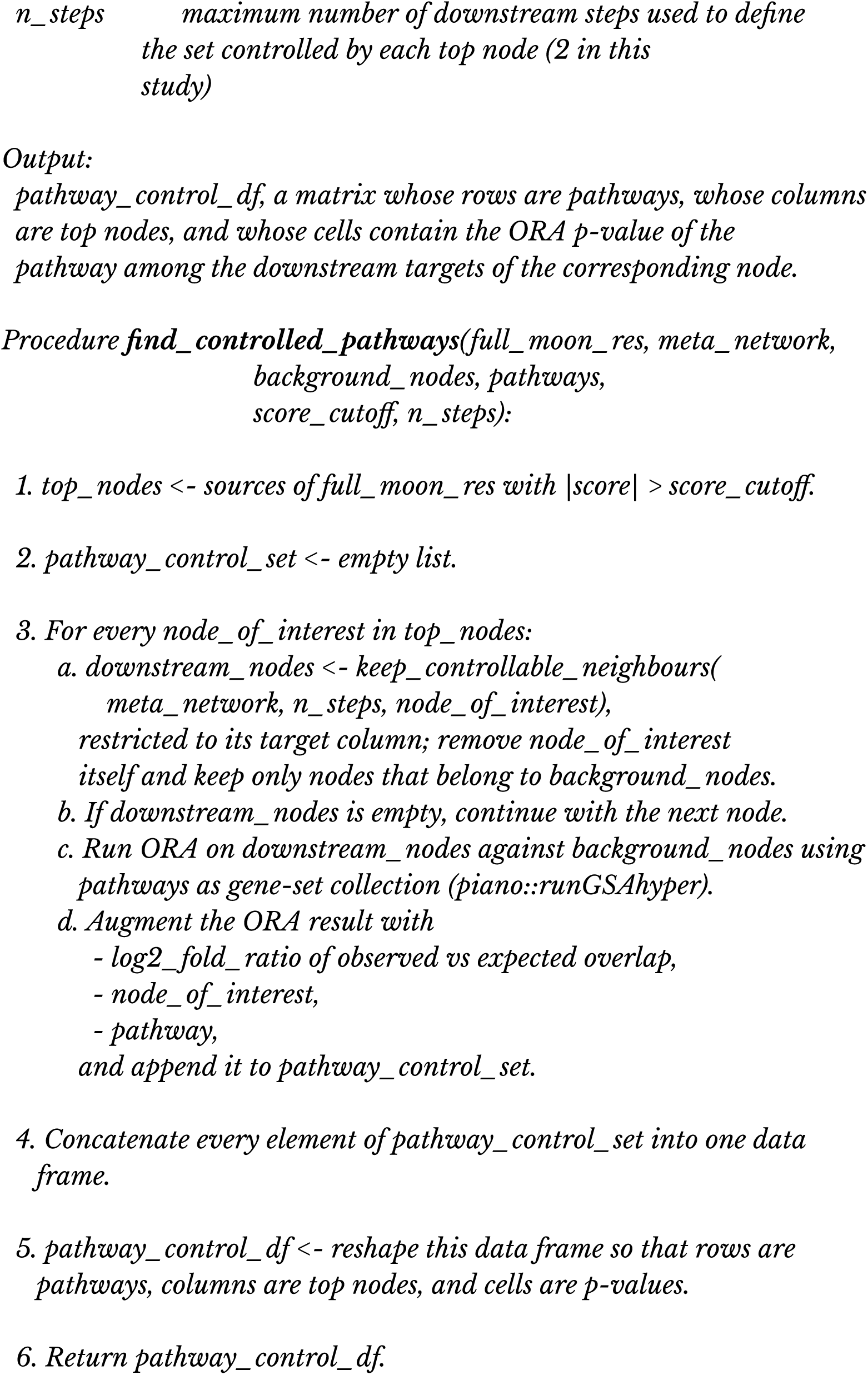

### 4.7 Cell line projection in factor space

To study how single samples behave in the factor space of MOFA, we reconstruct the RNA data of a given cell line with respect to each factor. To do so, we simply multiply the weights in a given factor with the corresponding value in the MOFA Z matrix for the corresponding cell line and the corresponding factor (Supplementary Figure S5). The algorithm describing this process is as follow:

#### Pseudocode to Decompose RNA Data for Specific Cell Line

##### Algorithm 8

decompose_rna_for_cell_line

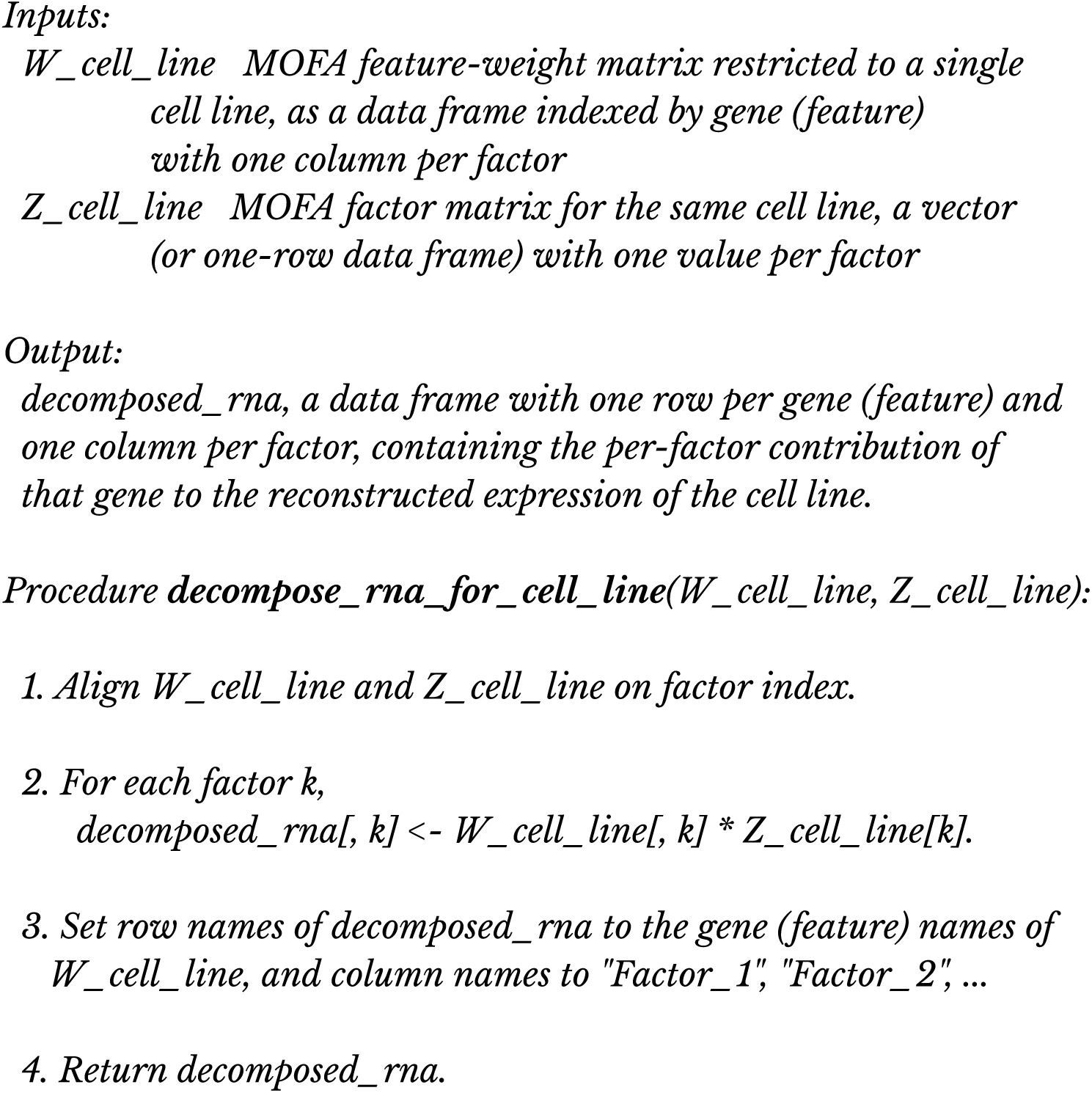

The data frame of decomposed RNA values for the cell line across factors can then be used seamlessly as an input for e.g. footprint analysis. In this case, we estimated TF activities and LR scores using the ULM function of decouleR, as described in Section 4.3.

### 4.8 Breast cancer cell line RNA/phospho data generation

The breast cancer cell line dataset was designed to probe the treatment effect of direct CDK2 inhibition in addition to CDK4/6 inhibition at multiple doses and timepoints provided. Three cell line models were selected for analysis as they provided a range of resistance to CDK4/6i:

- **Sensitive:** The MCF7 cell line model is a widely used breast cancer cell line model that responds robustly to CDK4/6i with potent RB1 dephosphorylation.
- **Partially Resistant:** The HCC1806 model is a CCNE_1_^amp^ TNBC partial responder derived cell line which shows partial response to CDK4/6i inhibition as indicated by RB1 phosphorylation modulation.
- **Resistant:** The MCF7SYL model is a fully resistant cell line to CDK4/6i with resistance possibly mediated by CDK2 and Myc which may be relevant to resistance mechanisms observed in patient data.

The cell lines were treated with five 5 inhibitors targeting the CDK pathway and estrogen receptor in monotherapy and in combination at multiple doses and time points, see Table XX. Specifically, the treatments corresponded to:

- Approved palbociclib a dual target CDK4 and CDK6 inhibitor,
- PF-06783600 a triple target CDK2, CDK4 and CDK6 inhibitor (Freeman-Cook *et al*, 2021b),
- PF-06974055, a single target CDK2 inhibitor, similar to Pfizer first in class CDK2 selective inhibitor PF-07104091 (Shen *et al*, 2024),
- approved fulvestrant, a selective estrogen receptor degrader (SERD),
- atirmociclib PF-07220060 a novel single target CDK4 inhibitor (Palmer *et al*, 2025)

This comprehensive list of both approved as well as novel CDK2,4,6 inhibitors and fulvestrant provided us with a diverse set of responses to CDK inhibition treatments. Measurements were performed in most cases at 4 time points: untreated control, early (2-4 hours), middle (24 hours) and late time point (72-96h).

#### 4.8.1 Cell culture

Parental cell lines were purchased from ATCC and cultured in RPMI-1640 medium supplemented with 10% FBS, 4 mM glutamine, and antibiotics (final concentrations: 50 U/mL penicillin, 50 µg/mL streptomycin) at 37°C in a humidified atmosphere with 5% CO2 using standard cell culture techniques. CDK4/6i resistant MCF7 model was generated with a starting exposure of 250 nM, and the concentration was increased 2-fold once cells reached parental growth rate. The final concentration for maintenance of CDK4/6i resistant lines is 4 µM. When cell growth rates in the presence of inhibitors returned to that of parental lines, resistance was evaluated in a standard proliferation assay.

#### 4.8.2 RNA-seq data generation

Cells were collected by trypsinization and washed with PBS (Phosphate Buffered Saline). Cell pellets were fast frozen with liquid nitrogen and then sent to Azenta life sciences for RNA extraction, library preparation and RNA sequencing (Illumina 2×150bp).

#### 4.8.3 Proteomics data generation

Cells were lysed in freshly prepared buffer (50 mM HEPES pH 8.5, 9 M urea, PhosSTOP), and protein concentration was quantified via BCA assay. Approximately 1.2 mg of protein per sample was digested with Lys-C (1:200 w/w, 2 hr, RT), diluted, and further digested with trypsin (1:100 w/w, overnight, RT). Digestion was quenched with 1% TFA, desalted (Sep-Pak C18), dried, and labeled using 18-plex TMT reagents. Equal amounts of labeled peptides were pooled, desalted, and fractionated via high-pH reverse-phase HPLC (XBridge C18, 60 min gradient, 0.4 mL/min), yielding 72 fractions. 1% of each was combined into 24 fractions for proteome analysis; the remaining 99% into 12 fractions for phosphopeptide enrichment using TiO₂ and Fe-NTA kits (Thermo Fisher). LC-MS/MS was performed on an Orbitrap Ascend with a 2-hour gradient (6–50% B), using data-dependent acquisition with MS1 (120K resolution) and MS2 (60K resolution) scans. Precursor ions (+2 to +7) were isolated (0.7 Da window), fragmented at 36% NCE for proteome and 34% for phosphopeptides, with dynamic exclusion of 20 s and ±10 ppm mass tolerance.

### 4.9 RNA and phospho data analysis of breast cancer cell lines

RNA counts were normalised using the VSN R package, differential analysis was performed using the limma R package. Each comparison for each time point was made with respect to time matched untreated cell line samples. Ligand-receptor scores were calculated using a ligand-receptor collection obtained from the Liana R package and the ULM function of the decoupleR R package. Transcription factor (TF) scores were calculated using the CollecTRI TF-target collection and the ULM function of the decoupleR package (with a minimum target set size of 5.

TMT phospho intensities were normalised separately for each TMT batch using the VSN R package, differential analysis was performed using the limma R package. Each comparison for each time point was made with respect to time matched untreated cell line samples. For each differential analysis resulting in a complete (no missing values) set of t-values, kinase/phosphatase scores were calculated using a custom set of kinase-target interaction (see below) and the ULM function of the decoupleR package (with a minimum target set size of 5). The custom set of kinase-target was obtained by merging a kinase-target set obtained from omnipathR using the Phosphosite, DEPOD and SIGNOR databases and high confidence (>0.95 score) kinasePhos kinase-target predictions. In order to generate the kinasePhos kinase-target predictions, kinasePhos3 (Ma *et al*, 2023) (https://github.com/tom-209/KinasePhos-3.0-executable-file) was run by selecting all kinase families available on the human reviewed set of protein sequences of Uniprot (https://www.uniprot.org/uniprotkb?query=*&facets=reviewed%3Atrue%2Cmodel_organism%3A9606). We used kinasephos3 instead of other similar tools (such as GPS6 or networkin) because it was the only one we managed to run in such a manner.

In order to compare the effect of synchronising or shifting the transcriptomic and phosphoproteomic data (asynchronous), samples were aligned by either matching the 24h RNA time point with the 24h phospho time point, or by matching the 24h RNA time point with the 2 or 4 hours phospho time points. Then, Pearson correlation matrices were generated to systematically compare phosphosite/gene pairs. For each phosphosite and gene, median correlation across all other gene/phosphosite was calculated, respectively, for both synchronous and asynchronous time point alignment. Gene Ontology was obtained from msigDB (c5.go.bp.v2024.1.Hs.symbols.gmt), and pathway scores were calculated using the ULM decoupleR function with median gene correlation coefficients as input, for both synchronous and asynchronous set up. The difference between pathway scores in each set up was obtained by a simple subtraction of the pathway scores.

COSMOS was used to integrate the transcription factors, ligands and receptors calculated from the transcriptomic data with the kinase activities calculated from the phosphoproteomic data. The prior knowledge network was obtained using the import_omnipath_interactions function of the omnipathR R package with default parameters on 12/01/2024. Receptor and ligand scores were obtained by splitting the ligand-receptor pairs and duplicating the ligand-receptor score over the ligand and receptors. Then, the resulting multiple scores for a given ligand or receptor (if a ligand is paired with multiple receptors, it would therefore have multiple scores) were averaged to obtain a single value for each ligand and receptor. Receptor scores were only kept if they didn’t already have a score based on kinase activity or TF activity estimations (as their activity is assumed to be more accurately determined by the expression/abundance of their targets rather than their own expression). The omnipath prior knowledge network was filtered to only keep genes that had measured expression in the RNA data set. When multiple concentrations were available in the phospho data but not in the RNA data, the closest matching concentrations were used (e.g. 100 nM in phosho data and 150nM in RNA data). For each contrast (treatment with a given inhibitor at a given time point), Cosmos was first run from receptors and kinases toward transcription factors. Receptors and kinases were kept as upstream input if their absolute score was above 2. The MOON algorithm was run with a maximum of 4 steps between receptor/kinases and downstream TFs, as described in <u>4.4.1 COSMOS with</u> <u>Meta-fOOtprint aNalysis (MOON)</u>. The MOON algorithm was also run with a maximum of 1 step between TF and ligand, to extract pairs of TF and downstream ligands that were sign-coherent. The two resulting networks for each contrast were then combined by keeping only the union of their edges where the sign of the MOON score of their common node where consistent.

### 4.10 Analysis of crispR KO data and validation of resistance mechanisms

Due to the large combination of timepoints, treatments and cell-line (22 contrast and therefore 22 combined MOON networks) to analysis, a PCA was performed on the combined MOON scores across 22 integrated phospho/RNA contrasts. The contrast PCA scores (that is the PCA coordinates) were then tested for enrichment of the various metadata variables (i.e. timepoints, cell lines and treatments) using the ULM decoupleR function. The strength of association of PCA factors was roughly estimated by the difference between the enrichment score of the resistant cell line HCC1806 and the sensitive cell line MCF7. crispR genome wide knockout data was obtained in three different conditions: control (i.e. basal essentiality or lethality), CDK4/6i (i.e. sensitisation to CDK4/6i treatment) and CDK2/4/6i (i.e. sensitisation to CDK2/4/6i treatment) for two cell lines, HC1806 (resistant to CDK4/6i and CDK2/4/6i) and MCF7 (sensitive to CDK4/6i and CDK2/4/6i). The crispR dataset was processed using the Mageck (Li *et al*, 2014) package to obtain z-scores for each gene representing the difference of cellular fitness with respect to specific gene knockouts. Then for each gene, in each cell line, the minimum value between 1) basal lethality, sensitisation to CDK4/6i and sensitisation to CDK2/4/6i, representing the most lethal potential of the gene or 2) sensitisation to CDK4/6i and sensitisation to CDK2/4/6i, representing the sensitisation potential of the gene. The values, referred to here as “most lethal”, from 1) and 2) were correlated with corresponding factor weights in each factor of the PCA performed on the MOON score matrix (see above), in order to determine if genes that tend to be differentially activated/inhibited between the resistant and sensitive cell lines are also important to sensitise/protect the resistant cell line from CDK4/6i and/or CDK2/4/6i. The resulting 21 factor/crispR correlation coefficients are then themselves respectively correlated with the difference between the association score of the resistant and sensitive cell of each factor. The resulting correlation coefficient therefore represents if the PCA factors that are associated with difference between resistant and sensitive cell lines are also associated with genes that are consistently more active/inactive in the resistant cell line compared to the sensitive one. It can then be interpreted as e.g. an indication of how much does a gene that is more active in a resistant cell line treated with a CDK inhibitor such as CDK4/6i actually is directly responsible for the resistance to the treatment.

In order to find which pathways showed an over-representation of genes that were both deregulated (in terms of activity computed by MOON) in resistant compared sensitive cell and sensitising (including/not including basal essentiality), a hypergeometric test was performed using the piano package, using as those genes as a set of significant genes, and using all other genes that did not agree between MOON and crispR background, specifically for the first component of the PCA (which showed the most difference of association between the sensitive cell MCF7 and the resistant cell HCC1806).

### 4.11 Paloma3 cohort analysis

Patient level RNA counts were z-transformed across patients. The ULM method of decoupleR was used with CollecTRI to estimate patient specific transcription factor activity signatures (XXX min target per TF). Patient specific COSMOS networks were generated using MOON with XXX steps up-stream from TFs. Using patient specific progression free survival values, COX survival models were computed for each gene of the one hand and each MOON score on the other hand, by splitting first patients by median gene expression or median MOON score, and then computing the COX hazard ratio between control and treatment group first, and second between high and low expression/moon scores.

Pearson correlation coefficient was computed between the MOON based COX coefficients and PC1 loadings from the cell-lines MOON score PCA. Precision-recall analysis was performed by setting the genes with MOON score based cox coefficients greater than 2 (higher coefficient represent association with worse treatment response) as true positives and computing the AUPRC of the PC1 loadings.

A hybrid COX coefficient signature was generated by combining gene expression based and MOON score based cox coefficients. The combination was defined as the union of both COX coefficient signatures, and for each gene that had both a gene expression based and MOON score based COX coefficient, the most extreme coefficient was selected. A scaled PFS prediction for each patient was computed by making a dot product between their gene expression value, MOON scores and the respective corresponding COX coefficient in the hybrid signature. A Pearson correlation coefficient was computed between the scaled PFS predictions and the original PFS values.

## 5. Acknowledgements

We acknowledge funding to J.S.R. by the German Federal Ministry of Education and Research (Bundesministerium für Bildung und Forschung BMBF) to support A.D. and D.T. (BMBF, [031L0181B]); by MSCoreSys research initiative research core SMART-CARE (031L0212A) to support A.D.; by Pfizer to support A.D. and R.F.; by HPC/Exascale Centre of Excellence for Personalised Medicine in Europe [PerMedCoE; European Union Horizon 2020 program, grant no. 951773] to support D.T.; D.M. acknowledges funding by the Spanish Government, which supports him through a predoctoral grant (PRE2020-092578 MCIN/AEI/10.13039/501100011033).

Thanks to Attila Gabor for his help maintaining the COSMOS package and setting up the documentation. Thanks to Ricardo Ramirez, Martín Garrido Rodríguez-Córdoba, Pablo Rodríguez Mier and Todd VanArsdale for their insightful advice on the manuscript.

## 6. Code and data availability

CosmosR package and NCI60 mofa-cosmos analysis markdowns can be found at https://saezlab.github.io/cosmosR/, while the source scripts and data can be found at https://github.com/saezlab/Factor_COSMOS. The cytosig benchmark analysis can be found at https://github.com/saezlab/MOON_benchmark. The python implementation of MOON can be found as part of a broader collection of tools at https://github.com/saezlab/networkcommons. The cytosig data was downloaded from the FDC platform (https://curate.ccr.cancer.gov/, accessed 22nd April 2022). The NCI raw data was accessed through NCI60 cellminer: transcriptomics (RNA-seq PMID:31113817), proteomics (SWATH (Mass spectrometry) PMID:31733513} and metabolomics (LC/MS & GC/MS (Mass spectrometry) DTP NCI60 data).

## 8. Authors contributions

AD and JS-R designed the method. AD and PL coded the pipeline. AD ran the analysis, DM, VP and DT reimplemented parts of the pipeline. RF and ACK helped with the data analysis. YL, MT, DN, SD, CS, JDL generated and processed cell line and clinical datasets and helped with the data analysis. BS and JS-R supervised the project. AD wrote the manuscript with help from PL and JS-R.

## Supplementary Materials

**Supplementary Figure S1.**
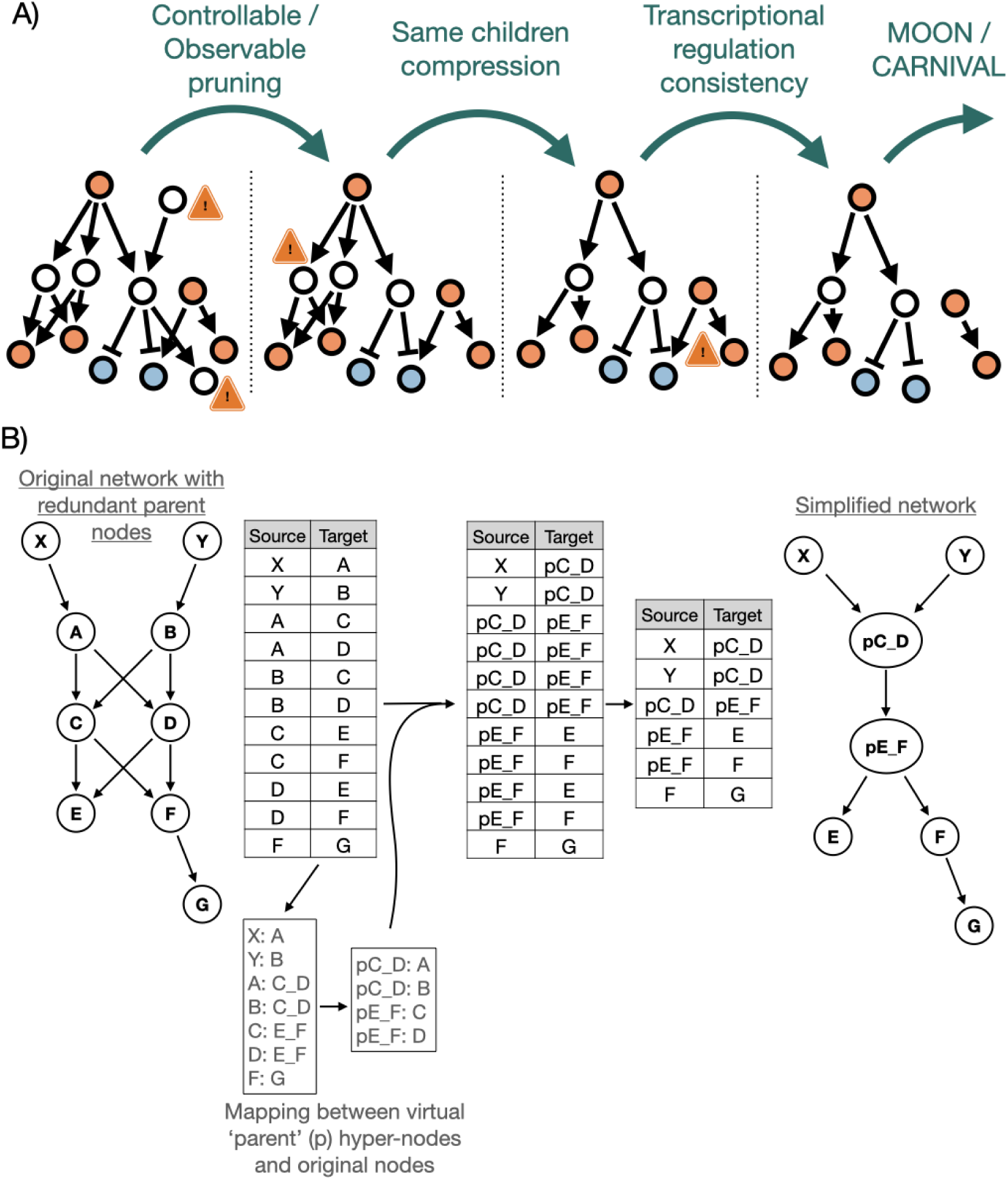
- A) Representation of the prior knowledge pre-processing steps. First, nodes that cannot be reached downstream of the upstream input (controllable) and nodes that cannot be reached upstream of downstream inputs (observables) are pruned out of the network. Then, any nodes that have strictly the same set of direct children nodes are compressed into virtual nodes, preventing redundant paths in the network. Next, a transcriptional consistency check is performed, where any node that is regulated directly by a transcription factor is pruned out if its transcriptional regulation direction doesn’t match the activity of said transcription factor. If proteomic data is available, both the transcriptional and proteomic regulation direction have to be consistent. The transcriptional consistency check is also performed after the network scoring, and the network scoring procedure is repeated until the transcriptional consistency check doesn’t find any inconsistency anymore. B) Schematic representation of the network compression algorithm.

**Supplementary Figure S2.**
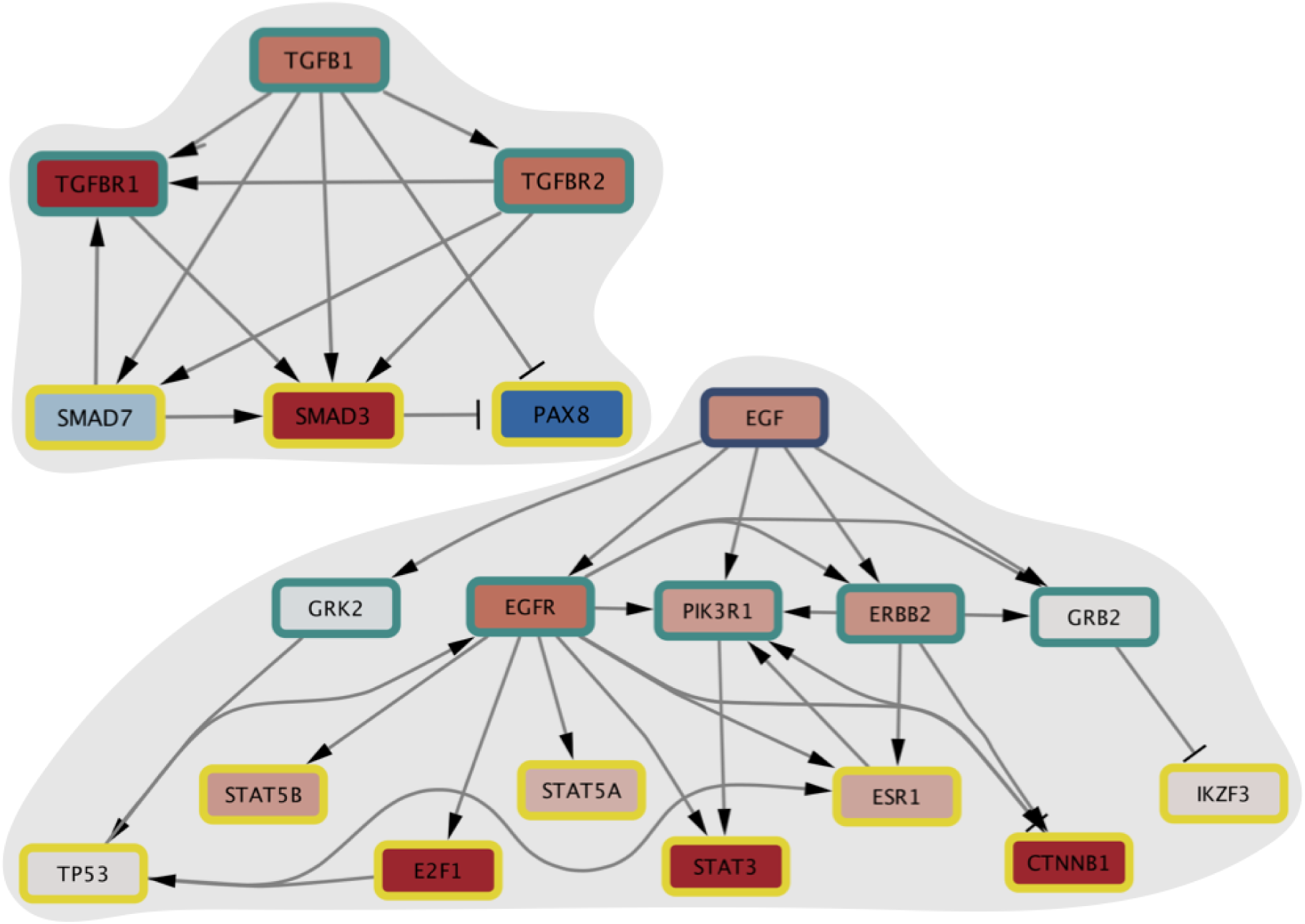
- TGFB1 also had relatively heterogeneous MOON scores across experiments, despite having a positive score on average (TGFB1: mean = 0.57, SD = 0.8).TGFB1 and EGF are examples of ligands with more complex MOON score estimation. Indeed, with respect to MOON heuristic, TGFB1 score depends on the activity estimation of SMAD3, SMAD7 and PAX8. While TGFB1 is relatively well scored (69th average quantile in experiments where it’s applied and 0.5 in experiments where it’s not), SMAD7 activity is systematically inconsistent with respect to the activity of SMAD3 and PAX8. Indeed, SMAD3 is expected to be activated by TGFB1 while PAX8 is expected to be inhibited, and their activity estimation is consistent with this in the example experiment of Figure 2B. However, while SMAD7 should seemingly be activated by TGFB1 according to the prior knowledge network of Omnipath, its activity isn’t increased when TGFB1 is applied. According to scientific literature **(Yan et al, 2009; Monteleone et al, 2001**), SMAD7 is in fact known to be an inhibitor of TGFB1 signaling via negative feedback mechanisms. EGF MOON score, on the other hand, appears to be depending on the activity of STAT5A, STAT5B, E2F1, STAT3, ESR1, CTNNB1 as well as TP53 and IKZF3 (Figure 2B). While STAT5A, STAT5B, E2F1, STAT3, ESR1, CTNNB1 all appear to be coherently deregulated, TP53 and IKZF3 do not seem to follow that trend. According to the prior knowledge network, TP53 and IKZF3 are regulated through GRK2 and GRB2 respectively. The EGF-GRK2-TP53 is documented in the scientific literature **(Gambardella et al, 2020; Chen et al, 2008**), albeit suggesting that GRK2-TP53 interaction should be an inhibition rather than an activation. EGF-GRB2 is also supported by scientific literature **(Yamazaki et al, 2002)**, but the GRB2-IKZF3 does not appear to have direct and targeted documented evidence in the scientific literature.

**Supplementary Figure S3.**
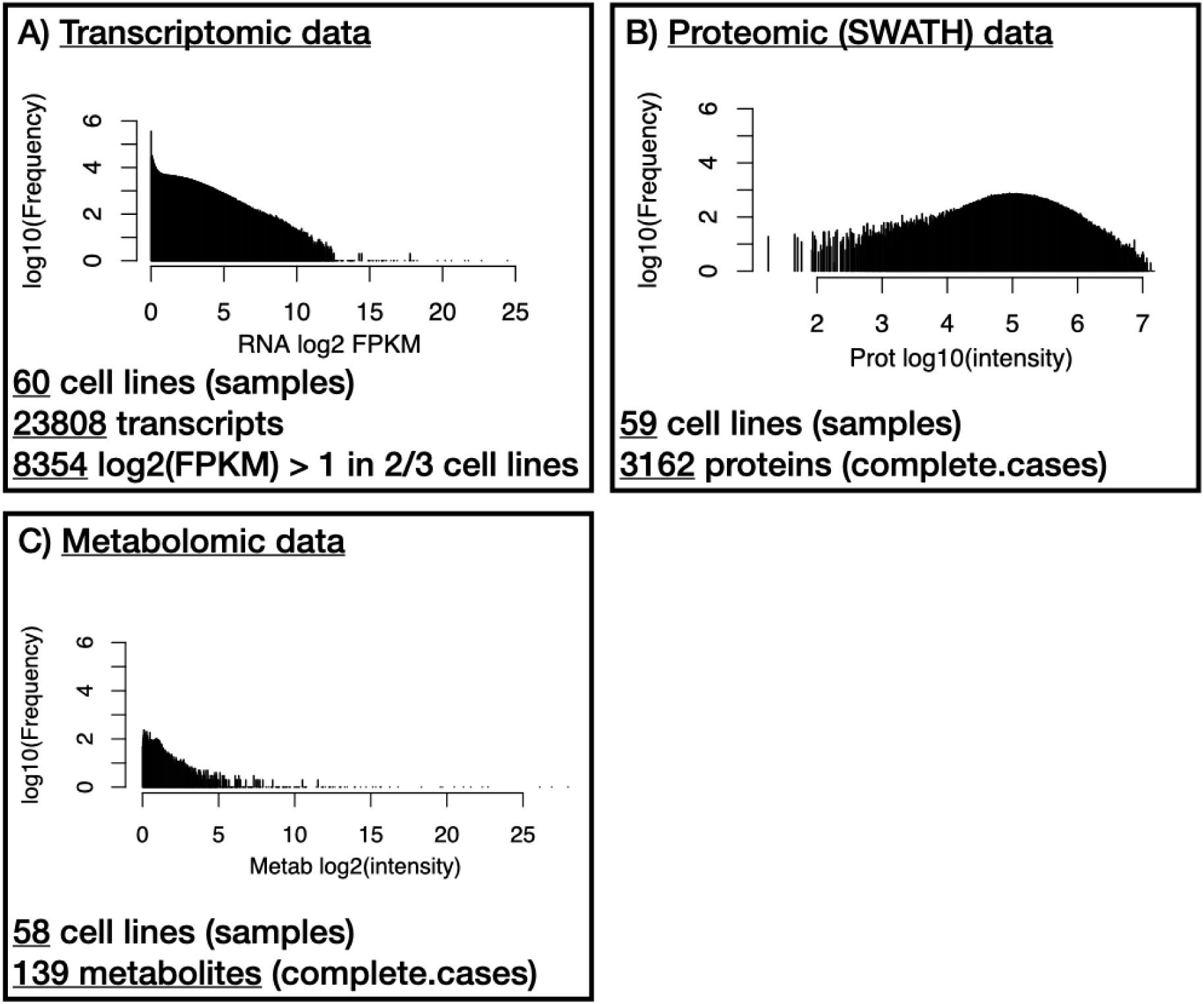
- NCI60 multi-omic data and MOFA. A) Histogram of all the log2 transformed FPKM+1 values of the NCI60 transcriptomic datasets. 60 cell lines have transcriptomic data in single replicates. 23808 transcript have annotated HUGO gene symbols, of which 8354 have log2(FPKM+1) values > 1 in at least ⅔ of the samples. log2(FPKM+1) values < 1 were excluded from the rest of the analysis (see “Prepare_RNA.R”). B) Histogram of all the log10 transformed intensities of the NCI60 proteomic dataset. 59 cell lines have proteomic data in single replicates. 3162 proteins are consistently detected in every cell line (complete cases). Data was obtained directly from cellminer and no further preprocessing was considered necessary (see “Prepare_proteomic.R”)). C) Histogram of all the log2 transformed intensity values of the NCI60 metabolomic datasets. 58 cell lines have metabolomic data in triplicates. To remain consistent with the other omic data, triplicates were simply averaged into single cell-line samples. A few extreme outliers (log2(intensities) > 32) were removed. 125 metabolites are consistently detected in every cell line (complete cases) (see “Prepare_metabolomic.R”).

**Supplementary Figure S4.**
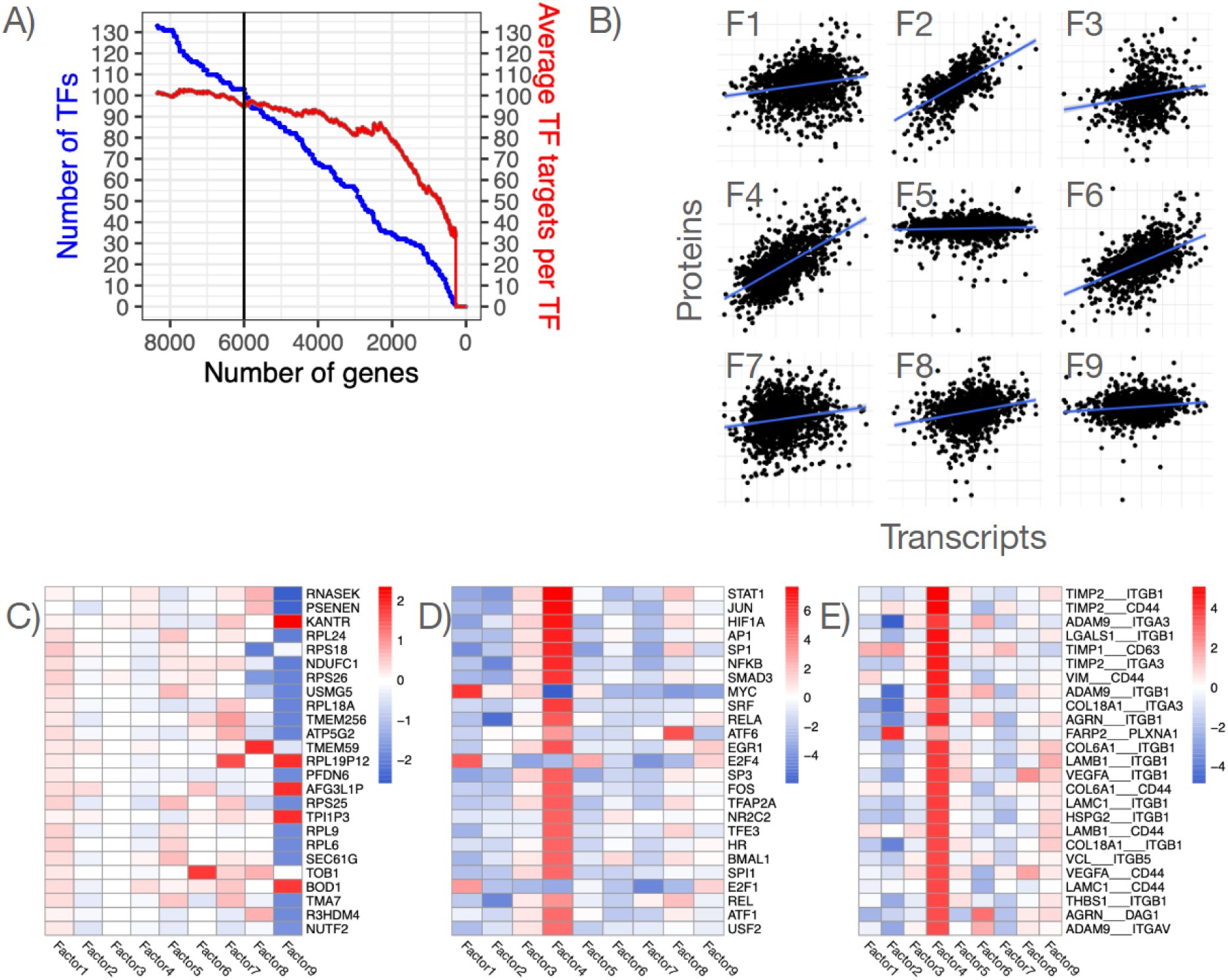
- MOFA and footprint analysis results. A) This plot shows how many TFs have enough measured targets (>10) to estimate their enrichment scores, given how many top variable genes are kept in the analysis. Number of TFs enrichment scores that can be estimated as a function of the number of top variable genes (blue line) and average number of targets per TFs as a function of the number of top variable genes (red line). B) Correlation of RNA and protein MOFA weights for each factor respectively. C) Heatmap of the top MOFA RNA features weights across 9 factors. The weights are ordered by their absolute maximum across all factors. D) Heatmap of the top transcription factors associated with MOFA RNA feature weights across 9 factors. The scores are estimated from RNA weights using the ULM method of decoupleR with the CollecTRI TF-target regulons and are ordered by their absolute maximum across all factors. E) Heatmap of the top ligand receptors associated with MOFA RNA feature weights across 9 factors. The scores are estimated from RNA weights using the ULM method of decoupleR with LIANA’s consensus ligand receptor prior knowledge and are ordered by their absolute maximum across all factors.

**Supplementary Figure S5.**
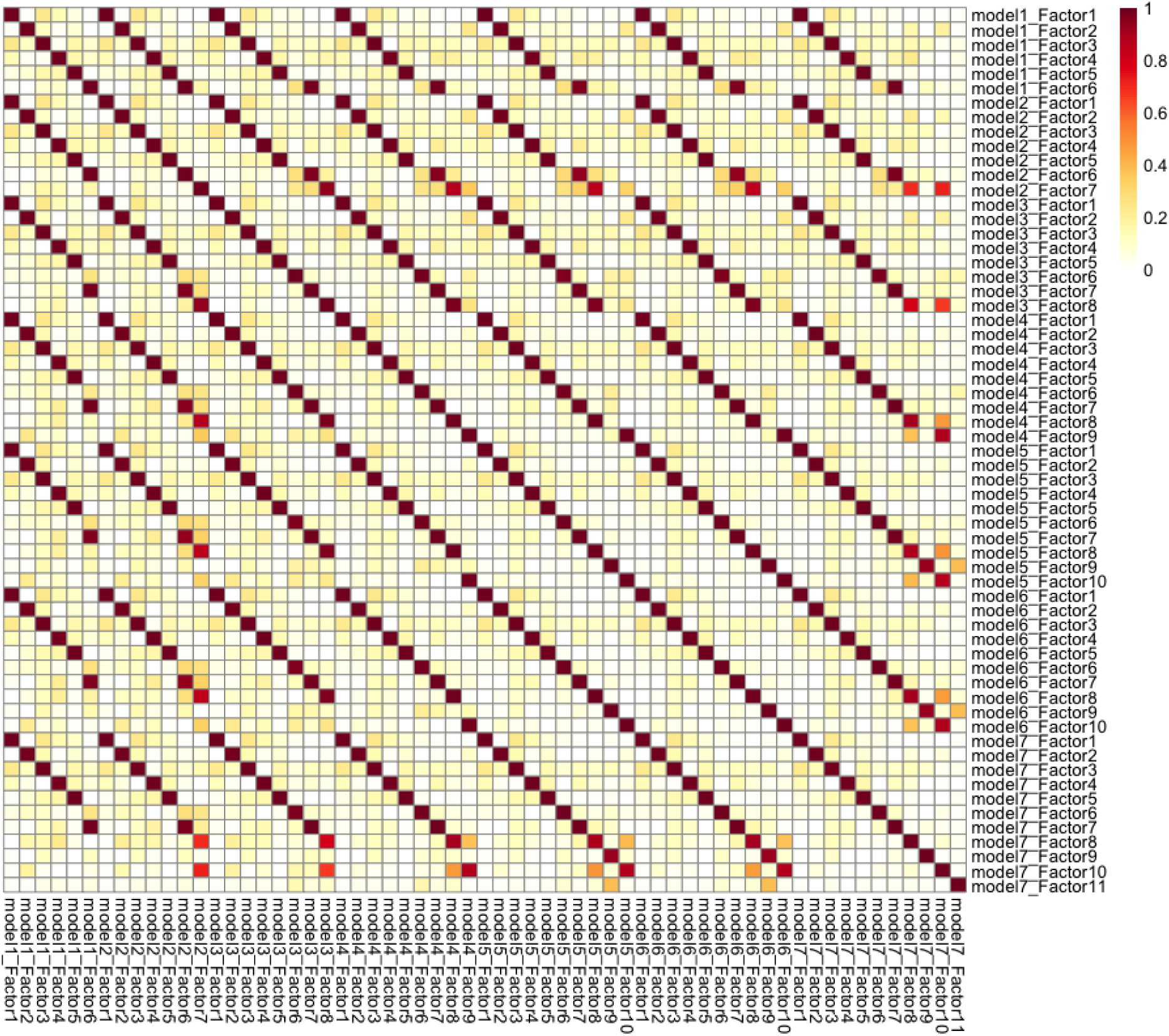
- Correlation heatmap between the factor weights of mofa models with different maximum number of factors (7 to 13 maximum factors).

**Supplementary Figure S6.**
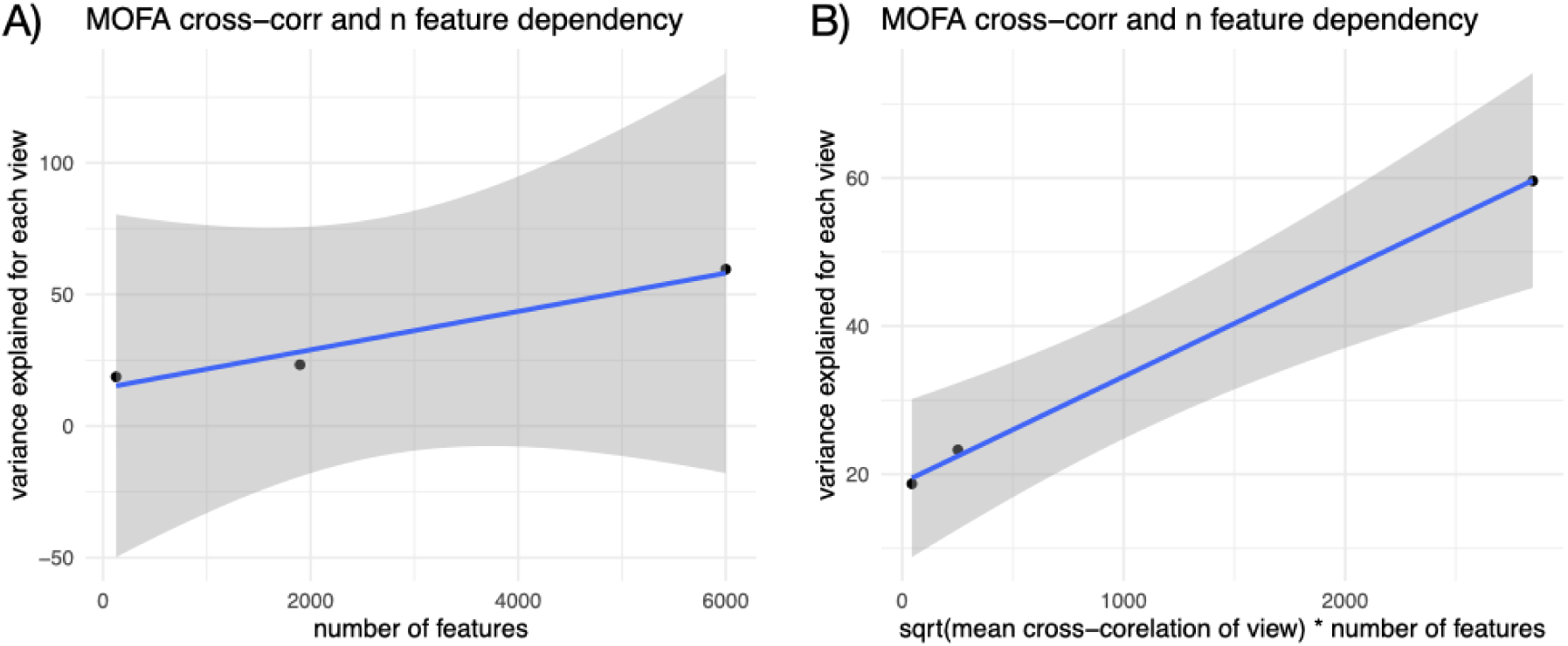
- Linear regression between the variance explained by the 9 MOFa factor for each omic view and A) the number of features of each view and B) the number of features of each view multiplied by the square root of the average cross correlation values of each view.

**Supplementary Figure S7.**
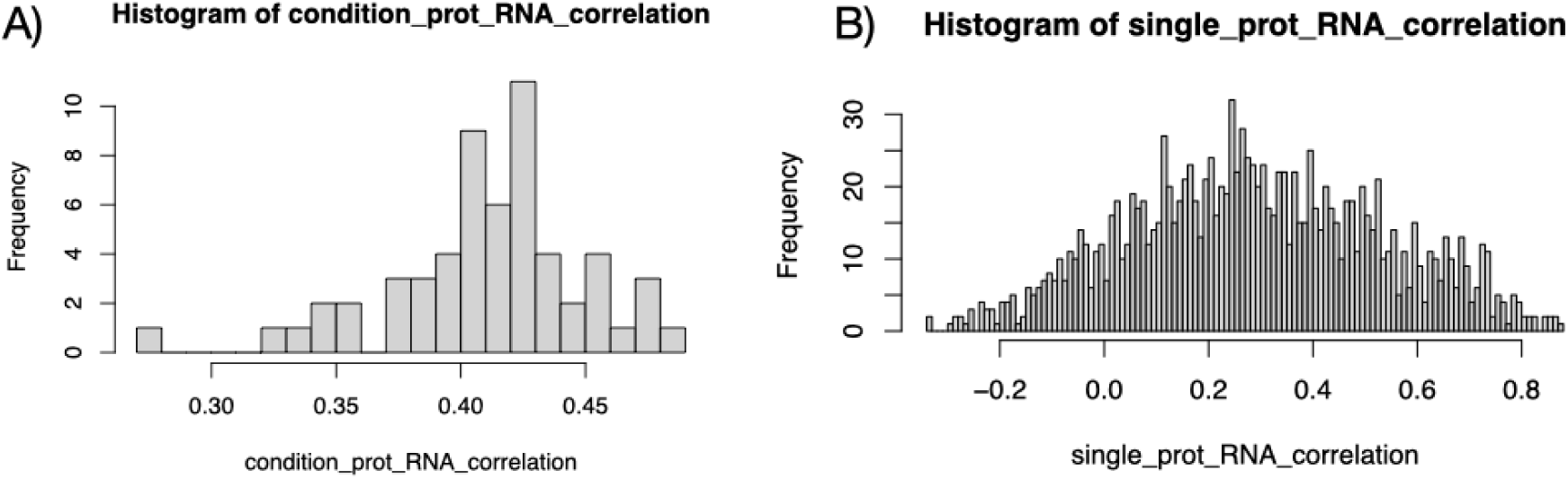
- A) histogram of correlation coefficients for each cell-line across genes between RNA and protein value. B) histogram of correlation coefficients for each gene across cell-lines between RNA and protein value

**Supplementary Text S1: Time point alignment for multi-omic samples** For CDK4, CDK2/4/6 and CDK2 inhibitors, phosphorylation measurements are available at both 2h and 24h while RNA measurements were available only at 24h. Therefore, phospho and RNA datasets can be aligned in synchronous (24h phospho -> 24h RNA) or asynchronous (2h phospho -> 24h RNA) manner. Since it is not in fact necessarily expected that matched time points are an optimal setting to integrate different types of omic data, we assessed the relative influence of the phosphosites measured at 2 or 24 hours on the expression of genes measured at 24 hours after treatment. To do so, we first computed the average correlation of each phosphosite with every gene across samples, and vice versa, in the synchronous and asynchronous sample alignment settings. Because different RNA samples are aligned with different phospho samples, the correlation coefficients are expected to be different between the two settings. Indeed, genes and phosphorylation sites were very slightly more correlated across samples (delta correlation coef = 0.003, t-test p-value = 10^-16^) on average in the synchronous setting. Averaging correlation coefficient between RNA and phosphorylation sites at the level of individual genes showed that gene-wise correlations were more conserved between synchronous and asynchronous settings (Supplementary Figure S8A Pearson r^2^ = 0.86) compared to phosphorylation site-wise correlations (Supplementary Figure S8B, Pearson r^2^ = 0.82). We compared which pathway enrichment scores were the most different between synchronous and asynchronous settings (using the difference of correlation t-values between synchronous and asynchronous correlations), which revealed that among the top 9 most enriched pathways, 3 of them were directly related to cell cycle (Supplementary Figure S8C). This indicated that while the differences between synchronous and asynchronous alignments are small, they seem to be preferentially localized within pathways that are directly relevant to the experimental set up (CDK inhibitor treatments). As the pathway enrichments for cell cycle seemed to be higher in the synchronous setting nonetheless, we chose the synchronous setting for the rest of the analysis.

**Supplementary Figure S8.**
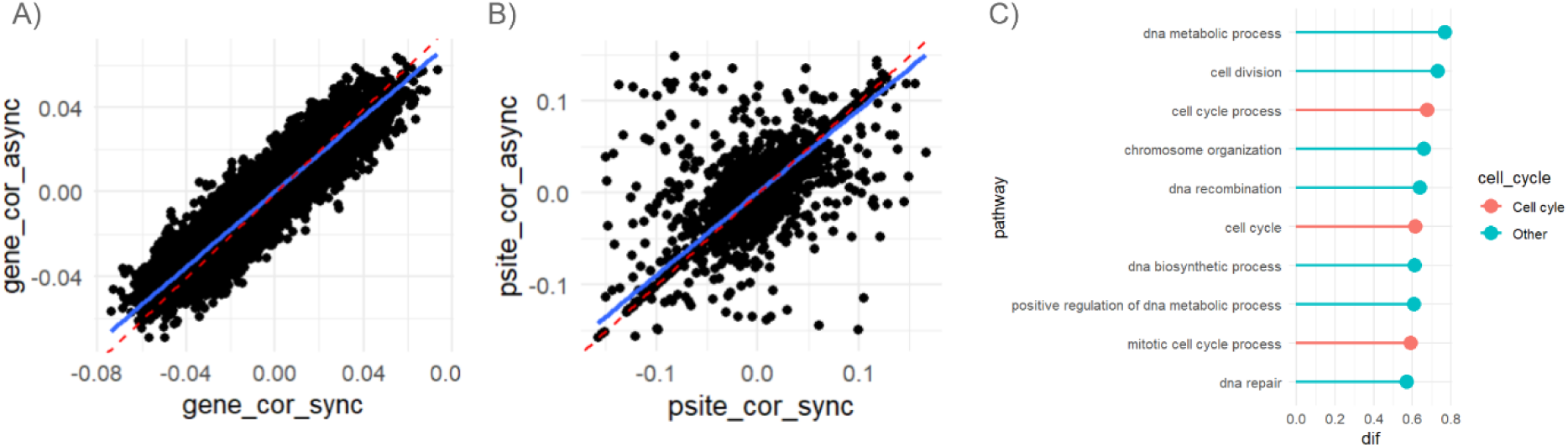
- A) Median correlations of genes with each psites, when 24h time points are synchronized (x axis) and when 24h RNA time points are matched with early (2h) phospho measurements (y axis). Red dashed line represent a unbiased expectation while the blue line represent the actual linear regression line of both set-ups. The steeper red line compared to the blue line indicates slightly higher correlations on average for the asynchronous alignment. B) Median correlations of psites with each gene, when 24h time points are synchronized (x axis) and when 24h RNA time points are matched with early (2h) phospho measurements (y axis). Red dashed line represent a unbiased expectation while the blue line represent the actual linear regression line of both set-ups C) Top 10 enriched pathways with differences (sync – async) in gene/phosphorylation correlation between the synchronous and asynchronous setting.

**Supplementary Table S1:** Summary of conditions where RNA and phospho-proteomic contrasts can be aligned. Each condition represents a contrast performed between the indicated cell line, drug and time point against a cell line/timepoint matched untreated sample.

| Condition | Cell | Resistance | Phospho<br>time point | RNA time<br>point | Drug |
| --- | --- | --- | --- | --- | --- |
| mcf7_fulv_2_4h | mcf7 | - | 2h | 4h | fulv |
| mcf7_fulv_72_96h | mcf7 | - | 96h | 96h | fulv |
| mcf7_fulv_cdk4i_2_4h | mcf7 | - | 2h | 4h | fulv_cdk4i |
| mcf7_fulv_cdk4i_72_96h | mcf7 | - | 96h | 96h | fulv_cdk4i |
| mcf7syl_3600_2_4h | mcf7syl | + | 2h | 4h | pf3600 |
| mcf7syl_3600_72_96h | mcf7syl | + | 96h | 72h | pf3600 |
| mcf7syl_fulv_2_4h | mcf7syl | + | 2h | 4h | fulv |
| mcf7syl_fulv_72_96h | mcf7syl | + | 96h | 96h | fulv |
| mcf7syl_fulv_cdk4i_2_4h | mcf7syl | + | 2h | 4h | fulv_cdk4i |
| mcf7syl_fulv_cdk4i_72_96h | mcf7syl | + | 96h | 96h | fulv_cdk4i |
| mcf7syl_palbo_2_4h | mcf7syl | + | 2h | 4h | palbo |
| mcf7syl_palbo_72_96h | mcf7syl | + | 96h | 72h | palbo |
| mcf7_3600_24h | mcf7 | - | 24h | 24h | pf3600 |
| mcf7_3600_72_96h | mcf7 | - | 96h | 72h | pf3600 |
| mcf7_4055_24h | mcf7 | - | 24h | 24h | pf4055 |
| mcf7_4055_72_96h | mcf7 | - | 96h | 72h | pf4055 |
| mcf7_palbo_24h | mcf7 | - | 24h | 24h | palbo |
| mcf7_palbo_72_96h | mcf7 | - | 96h | 72h | palbo |
| hcc1806_3600_24h | hcc1806 | + | 24h | 24h | pf3600 |
| hcc1806_3600_72_96h | hcc1806 | + | 96h | 72h | pf3600 |
| hcc1806_palbo_24h | hcc1806 | + | 24h | 24h | palbo |
| hcc1806_palbo_72_96h | hcc1806 | + | 96h | 72h | palbo |

